# Immunoengineered Chitosanase-Produced Chitosan Oligomers for Elevating Plant Resistance to Viral Infection

**DOI:** 10.64898/2026.06.09.731087

**Authors:** Soofia Khanahmadi, Ratna Singh, Judith Ryll, Naivy Y. Nava Cruz, Stefan Cord-Landwehr, Carolin Richter, Alireza Rafieerad, Bruno M. Moerschbacher

## Abstract

Chitooligomers can act as plant biostimulants or biopesticides, but today’s chitosan-based agro-biologics often lack sufficient efficacy. This is due to a lack of scalable production processes for structurally well-controlled chitosans combined with a limited understanding of structure-function relationships. Chitosans differ in their degree of polymerization (DP), fraction and pattern of acetylation (FA and PA). While the influence of DP and FA on antimicrobial and phytostimulatory properties is at least partially known, this is not yet the case for PA. PA can be partially controlled by using enzymatic rather than acid hydrolysis for oligomer production. We have used recombinant chitinases and chitosanases to hydrolyse a well-characterised chitosan polymer, and purified oligomers with different DP. We have structurally characterised the products and tested their abilities to protect tobacco from viral disease. Chitinase products were dominated by GlcNAc units at their reducing and non-reducing ends, with GlcN units dominating their centers, and v.v. for chitosanase products. While the chitinase-derived hydrolysates were inactive, the chitosanase-derived oligomers possessed elicitor and priming activities and protected plants from disease, and their activity increased with increasing DP. Clearly, the *Bacillus* chitosanase used is well-suited to set up a scalable production process for chitosan oligomers with promising agro-biologic properties.

**Graphical Abstract:** 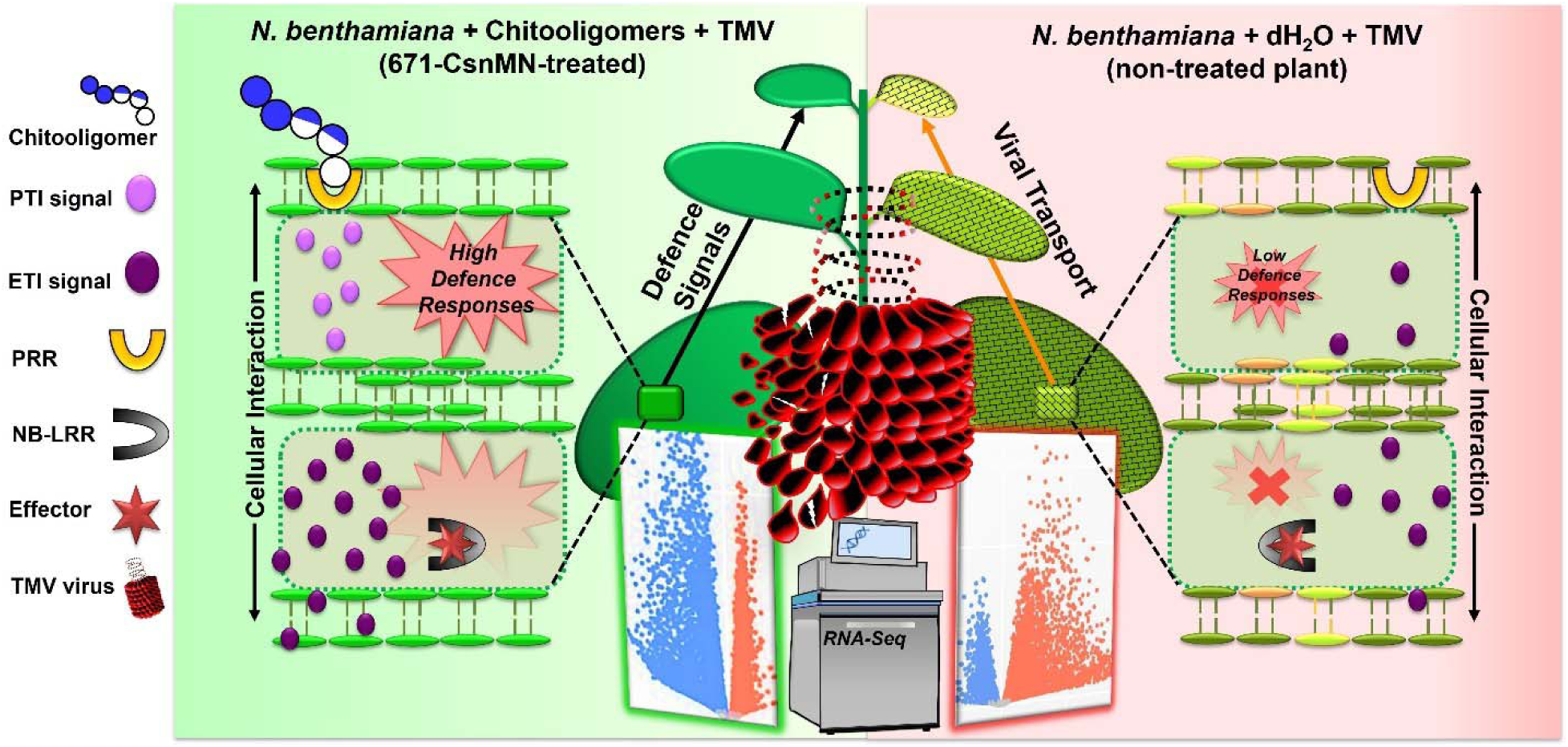

## 1. Introduction

Plants are under constant attack by pests and pathogens such as insects and nematodes, fungi, oomycetes, bacteria, and viruses, causing reductions in the yield and quality of crops. Pesticides are a proven way of limiting these losses, but they are environmentally problematic, potentially cause health problems with farmers and consumers, and are becoming increasingly inefficient due to resistance developments in the pests and pathogens. More sustainable alternatives for plant protection are resistance breeding and plant immunoengineering. While the former approach introduces resistance genes e.g. from wild relatives of the crop plants into modern, high-yielding crop varieties, the latter aims to develop tools to modulate and harness the potential of the plants’ own immune system.

Natural biopolymers such as alginates and ulvans from algal cell walls or chitins and chitosans from fungal cell walls or arthropod exoskeletons are known to possess plant eliciting or priming properties (Kahsay Meresa et al., 2024). While elicitor-active compounds trigger acute disease resistance mechanisms such as phytoalexin biosynthesis or hypersensitive cell death, priming-active compounds act like vaccines, increasing the plant’s ability to quickly and efficiently mount resistance reactions once a pathogen attacks (Liu et al., 2022). Often, the same compound can act as priming agent at low concentration, while exhibiting elicitor activity at high concentrations. Clearly, priming activity is more suitable for plant protection, but it is often associated with a growth-defense trade-off (He et al., 2022).Interestingly, this does not always seem to be the case, e.g. when a priming-active compound leads to increased phytosynthetic capacities providing extra resources for disease resistance reactions, as recently reported for chitosan-treated potato plants (Lemke et al., 2020).

However unfortunately, chitosan-based products, like many other agro-biologics, often suffer from low efficacy, poor reliability, and incompatibility with other agro-biologics or agro-chemicals (Bellich et al., 2016; Ebert, 2019; Hadwiger, 2013). These problems are particularly prominent when structurally and functionally ill-defined raw chitosans are used, which often exhibit high batch-to-batch variability (Wattjes et al., 2020). When such chitosans are used, as is the case very often in literature, experiments cannot be reproduced faithfully and results cannot be verified reliably. Chitosans are linear aminopolysaccharides composed of *N*-acetyl-D-glucosamine (GlcNAc, A) and D-glucosamine (GlcN, D) units linked by β-1,4-glycosidic bonds. They are structurally characterized by their degree of polymerization (DP), their fraction of acetylation (*F*_A_), and their pattern of acetylation (PA), all of which crucially influence their bioactivities (Cord-Landwehr et al., 2020; Wattjes et al., 2020). The situation is further complicated by the fact that chitosans can have both antimicrobial and phytostimulatory activities, so that it is not easy to distinguish between plant immunoengineering effects versus direct antimicrobial effects. As a consequence, molecular structure-function relationships as well as cellular modes of action of chitosans’ phyto-and immunostimulatory activities are still only poorly understood.

While it is clear that sufficiently large (DP ≥6) and highly acetylated (*F*_A_ ≥0.6) chitin and chitosan oligomers, provided they have a suitable PA, can be perceived by their interaction with pathogen-recognition receptors (PRRs) at the surface of plant cells, such as the heterooligomeric chitin receptors AtCERK1/AtLyk4/AtLyk5 in *Arabidopsis thaliana* or OsCEBiP/OsCERK1 in *Oryza sativa* (Narusaka et al., 2013; Shimizu et al., 2010), the perception of smaller or more strongly deacetylated oligomers is not yet understood. Yet, even small chitin and chitosan oligomers can possess bioactivities, such as plant growth-promoting chitin tetramers (Winkler et al., 2017) and plant priming-active monoacetylated chitosan tetramers (Basa et al., 2020). In addition, chitosan polymers may exert their effects on microbial and, potentially, also on plant cells by less specific electrostatic surface interaction between positively charged chitosans and negatively charged target molecules or structures, such as cell walls or membranes, possibly impairing cellular integrity and triggering stress reactions or even cell death (Kumaraswamy et al., 2018). It also needs to be taken into account that chitosan polymers applied to a plant or to soil will invariably be enzymatically processed in situ, yielding partially acetylated chitosan oligosaccharides (paCOS) which typically differ in all three properties from the original polymers(Cord-Landwehr et al., 2020). In infected plants, thus, receptor-mediated and target-mediated responses most likely occur in parallel, possibly even acting synergistically (Attjioui et al., 2021).

As a consequence, mixtures of chitosan polymers and oligomers are particularly promising, and these can easily be obtained by partial depolymerization of chitosan polymers using physical, chemical, or enzymatic processes (Gonçalves et al., 2021; Poshina et al., 2018). The latter have the advantage of yielding products with partially controlled PA, namely possessing non-reducing and reducing end units equivalent to the GlcN-and GlcNAc-preferences of the plus-and minus subsites of the enzymes, and lacking cleavage sites for the enzyme used in their production. Surprisingly, despite a multitude of studies on using chitosan oligomers as an agro-biologic, including oligomers produced by enzymatic hydrolysis (Boamah et al., 2023), very few studies have exploited this possibility of improved structural control of the enzymatic products by using enzymes with distinct cleavage specificities to understand the influence of PA on bioactivities. In an early report, Rahman et al. (Rahman et al., 2014) showed that products of a GH46-chitosanase (cleavage preference-X-D | X-X-; with D = GlcN specificity or very high GlcN preference, X = GlcN or GlcNAc, | = cleavage site; thus possessing GlcN-units at their reducing ends (Heggset et al., 2010)) inhibited fungal germination more strongly than products of a GH18-chitinase (-X-A | X-X-; with A = GlcNAc specificity or very high GlcN preference; thus possessing GlcNAc-units at their reducing ends (Brurberg et al., 1994). As already mentioned above, Basa et al. (Basa et al., 2020) then showed that small GH8-chitosanase (-D-D | X-X-) products, but not GH18-chitinase (-X-A | X-X-) products have priming activity in rice cells. And recently, Richter et al. (2025) reported that enzymatic hydrolysates using a GH8-chitosanase (-D-D | X-X-) or a GH18-chitinase (-X-A | X-X-) possess distinct antibacterial and antifungal activities as well as elicitor and priming activities in *Arabidopsis* plants. The latter study also revealed that this was not the case when the hydrolysate was produced chemically using acetic acid, which cleaves the polymer at random sites.

Here, we have built on these results, producing chitosanase hydrolysates using the above described GH8-chitosanase (-D-D | A-X-) and GH18-chitinase (-X-A | X-X-), but also a GH19-chitinase (-A-X | A-X-), or hydrochloric acid (rather than acetic acid) as a control. We assessed the elicitor and priming activities of these hydrolysates in *Nicotiana benthamiana* plants as well as their ability to protect the plants against tobacco mosaic virus (TMV). Combining the results of the current with the former study (Richter et. al 2025) reveals that the chitosanase hydrolysate has the strongest plant immunoengineering activity, both in *A. thaliana* and in *N. benthamiana*, has the strongest antimicrobial activity against bacteria and fungi, and induces local and systemic resistance against viral disease. Clearly, the chitosanase-produced chitosan hydrolysate is well suited for the development of a potent agro-biologic for sustainable agriculture.

## 2. Materials and Methods

### 2.1. Enzymatic hydrolysis of chitosan

We depolymerized a commercially available, microcrystalline chitosan prepared by partial heterogeneous alkaline de-*N*-acetylation of chitin extracted from shrimp shell waste (*F*_A_ ca. 0.2; DP ca. 800; *Ð* ca. 1.5; coded “651”) (Gillet Chitosan, Plumaudan, France) to produce three different hydrolysates (coded “671”). The enzymes used for depolymerisation were produced recombinantly as described in Table 1. Briefly, 0.1 g of chitosan 651 was dissolved in 100 mL of deionized water (dH2O), including 25 µL of acetic acid 100 % (pH ca. 5.5). The solution was mixed thoroughly using a magnetic stirrer and then filtered using 0.22 µm pore size filters. The resulting solutions were incubated with 1 µg of chitosanase (Regel et al., 2018b) or 10 µg of either of two chitinases (Bußwinkel et al., 2018; Heggset et al., 2009a) at 37 °C for 24 or 96 hours, respectively. The products were filtered using Vivaspin centrifugal concentrators with 10 kDa molecular weight cutoff polyethersulfone membranes (Sartorius, Göttingen, Germany). Filtrates were lyophilized overnight, dissolved in autoclaved pure water at a concentration of 1 mg mL^□1^, and stored at 4 °C until further use. A partial hydrochloric acid hydrolysate of the same 651 chitosan polymer (coded “661-Cl”) used for comparison was prepared and provided by the same producer. Fig. 1 gives an overview of the production processes for the different hydrolysates.

**Figure 1:**
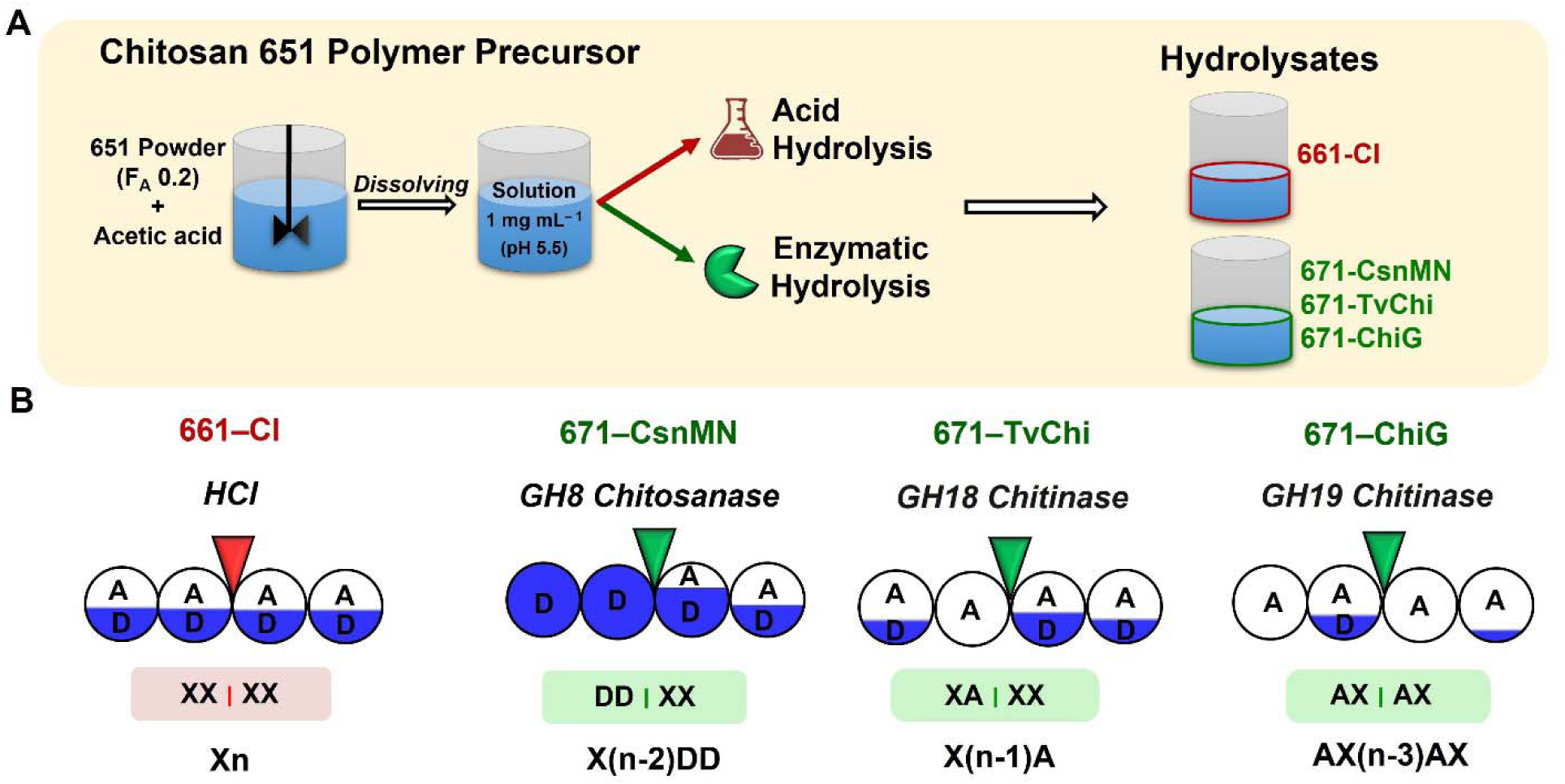
**Preparation of the chitosan hydrolysates**. A) Overview of the production process for the acid hydrolysate 661-Cl and the enzymatic hydrolysates 671-CsnMN, 671-TvChi, and 671-ChiG, produced from chitosan polymer 651 (*F*_A_ ca. 0.2; DP ca. 800; *Ð* ca. 1.5). B) Cleavage preferences during chemical and enzymatic hydrolysis and sequence of the chitosan oligomers obtained. The red and green triangles indicate the cleavage site; balls left of it indicate subsites-2 and-1 towards the substrate’s non-reducing end, those right of its subsites +1 and +2 towards its reducing end; subsite preferences for GlcNAc (A) and GlcN (D) are indicated by the white and blue background, respectively (see Table 1 for references). Simplified cleavage sites and the sequence of products with DP n found in the hydrolysates are indicated below the pictograms (A = GlcNAc, D = GlcN, X = GlcNAc or GlcN).

**Table 1:** Specifications and production parameters of enzymatically produced chitosan hydrolysates. For all samples, chitosan polymer 651 (*F*_A_ 0.2, DP 800) was incubated with chitosanase or chitinase to produce the different 671 hydrolysates.

| Sample Name | Enzyme used for the production | Type of enzyme | GH family | Enzyme origin | Reference describing enzyme production and specificity |
| --- | --- | --- | --- | --- | --- |
| 671-CsnMN | CsnMN | Chitosanase | GH8 | *Bacillus* sp. MN | Regel et al., 2018a |
| 671-TvChi | TvChi | Chitinase | GH18 | *Trichoderma virens* | Bußwinkel et al., 2018 |
| 671-ChiG | ChiG | Chitinase | GH19 | *Streptomyces coelicolor* A3(2) | Heggset et al., 2009b |

### 2.2. Preparation of DP pure oligomer fractions using size-exclusion chromatography (SEC)

Size exclusion chromatography was used to separate and collect the chitosan oligomers in the previous produced 671 samples according to their DP. A SECurity GPC system (Polymer Standards Service PSS, Mainz, Germany) with three HiLoad 26/600 Superdex 30 preparatory grade columns (GE Healthcare, Chicago, IL USA), a refractive index detector (Agilent 1260 infinity, Mainz, Germany), and a fraction collector FRAC-2000 (Pharmacia, Uppsala, Sweden) was used to achieve this. Briefly, 5 mL of the hydrolysates at a concentration of 10 mg mL^□1^ were injected into the SEC system. Ammonium acetate buffer (0.15 M) with 200 mM acetic acid at a pH of 4.5 was used as a mobile phase with a flow rate of 0.6 mL per minute. Starting 10 h after injection, fractions were collected every 10 min. Fractions containing paCOS with the same DP were pooled, freeze-dried overnight, dissolved in pure dH_2_O, and freeze-dried again to eliminate excess ammonium acetate.

### 2.3. Hydrophilic interaction liquid chromatography electrospray ionization mass spectrometry (HILIC-ESI-MS)

DP and *F*_A_ of the chitosan oligomers were analyzed using a Dionex Ultimate 3000RS UHPLC system (ThermoFisher Scientific Co., Waltham, MA, USA) coupled to an amaZon speed ESI-MS^n^-detector (Bruker, Bremen, Germany) according to Hamer et al. (Hamer et al., 2015). Separation was done using hydrophilic interaction liquid chromatography (HILIC) with a Waters Acquity UPLC BEH Amide column (2.1 mm × 50 mm, 1.7 µm) combined with a 1.7 µm VanGuard pre-column (2.1 mm × 5 mm, Waters Corporation, Milford, USA). The flow rate and column oven temperature of the system were set to 0.4 mL min^-1^ and 40 °C. Samples (2 µl at a concentration of 1 mg mL^-1^) were injected and eluted from the column using a gradient of solvent A (acetonitrile/water, 80:20 (v/v)) and solvent B (acetonitrile/water, 20:80 (v/v)), both containing ammonium formate (10 mM) and formic acid (0.1% v/v). The following gradient of solvent A and B was used to separate the chitosan oligomers in the different fractions: 0 to 0.833 min isocratic 100 % A; 0.833 to 4.5 min linear from 0-100 % B, 4.5 to 8.66 min 100 % B, 8.66 to 9 min linear from 100 to 0 % B, 9 to 10 min 100 % A). Samples were analysed in positive mode in the MS with an end plate voltage of 500 V, a capillary voltage of 4 kV, a nebulizer pressure of 15 psi, a dry gas flow rate of 8 l min^-1^, and a dry temperature of 200 °C. The detector’s target mass was set to m/z 700. Mass spectra were acquired using the ultra-scan mode with a scan rate ranging from m/z 160 to 2000. The determined *m/z* values of the HILIC-MS analysis were used to identify the oligomers in the mixture with the help of Data Analysis 4.1 software (Bruker, Bremen, Germany) and the monoisotopic masses of the chitin and chitosan oligomers.

### 2.4. Cultivation of N. benthamiana

All plant cultivation and maintenance processes were carried out under standard conditions of 14/10 h photoperiod at the University of Münster’s greenhouse facilities. Wild-type *N. benthamiana* seeds were first grown to seedlings, potted in soil, and grown at 24. Four-week-old plants were transferred to a standard climate chamber under the same conditions and kept for 24 hours before starting experiments to avoid uncontrolled stresses.

### 2.5. Eliciting activity of chitosans

#### 2.5.1. Measurement of elicitor activity in N. benthamiana leaf discs

Leaf discs were cut from the fully-developed middle leaves of the plants using a 5 mm cork borer. The leaf discs were then gently immersed into 96-well microtiter plates (Nunc 96-Well MicroWell, Thermo Fisher Scientific, Waltham, USA) containing 100 µL of pure dH_2_O per well. The plates were incubated overnight in the dark in a climate chamber at 24 to decrease stress levels. Elicitor activities were quantified by measuring the H_2_O_2_ released from the leaf discs, using a chemiluminescence oxidative burst assay (Albert & Fürst, 2017; Nishinaka et al., 1993). One day after transfer, the water in the wells was replaced by 200 µL of a 1:1 mixture of elicitor solution and 0.5 mM of luminol derivate L-012 (8-amino-5-chloro-7-phenylpyrido[3,4-d]pyridazine-1,4(2H,3H)dione) in 10 mM of 3-morpholino-propane-1-sulfonic acid/potassium hydroxide (MOPS/KOH) buffer at pH 7.4. The plates were placed into a microtiter plate reader (Lumat Varioskan LUX, Thermo Scientific), and luminescence was measured for 1 s per well every 2 min at room temperature for 90 min. The values of the maximum relative light units (RLUmax) reached during this interval were used to compare the elicitor-activities of the compounds.

### 2.5.2. Measurement of elicitor activity in N. benthamiana seedlings

Seeds of *N. benthamiana* were surface-sterilized using 70 % ethanol for 1 min followed by soaking in 3 % sodium hypochlorite for 2 min and washed five times with dH_2_O before seeding them on a Petri dish containing half-strength Murashige and Skoog medium (Duchefa Biochemie Co). The plates were sealed with parafilm, transferred to a climate chamber, and gently shaken at 24 for 7 d. Single 6-to-7-day-old seedlings were transferred to each well of 96-well microtiter plates to which 165 µL of MGRL culture medium(Naito et al., 1994) containing 1 % sucrose was then added. The plates were sealed and incubated in a climate chamber with 14/10 h photoperiod for 24 h. Elicitor activities were quantified using H_2_O_2_ measurements as above, adding 55 µL of a 1:1 mixture of elicitor solution and L-012 (0.25 mM in 50 mM potassium phosphate buffer (KPP) at pH 7.9) to each well.

### 2.6. Priming activity of chitosan hydrolysates

#### 2.6.1. Measurement of priming activity in N. benthamiana leaf discs

Test solutions (500 µL, 100 µg mL^□1^) were sprayed onto the adaxial side of a fully grown leaf with airflow at a maximum air pressure of 3.1 bar (SPARMAX Air compressor AC-55, DiscQver). Plants were kept for 2 h in a climate chamber (14/10 h photoperiod at 24) before harvesting discs from the treated “local” leaf and from the “systemic” leaf immediately above the local leaf, as described above. The oxidative burst was triggered by the addition of flg22 (EZBiolab, Carmel, USA) at a final concentration of 0.1 µg mL^-1^. Alternatively, the chitosans at concentrations of 10, 30, and 100 µg mL^-1^ were added to a Murashige-Skoog (MS) agar (Murashige & Skoog, 1962) in Incu Tissue vessels (72 × 72 × 100 mm, SPL) and placed on a sterilized bench for solidification. Then, 10-day-old seedlings were transferred to the agar using sterilized tweezers. After three weeks, leaf discs were harvested for oxidative burst measurements as above, using flg22 as an elicitor.

#### 2.6.2. Measurement of priming activity in N. benthamiana seedlings

Seven-day-old seedlings were transferred into Petri dishes containing chitosan solutions in non-solidified MS medium. The Petri dishes were then gently shaken inside a climate chamber. One day later, the seedlings were transferred to a microtiter plate containing 165 µL of MGRL nutrient medium complemented with 1 % sucrose (w/v). The plates were sealed and incubated in a climate chamber for 24 h, and oxidative burst measurements were performed as mentioned above. The concentration of flg22 was 0.025 ng mL^-1^ in this assay.

### 2.7. In planta antiviral properties of chitosans

To assess the antiviral activity of the samples against TMV in planta, we sprayed the chitosans at a concentration of 100 µg mL^-1^ onto the adaxial side of a middle “local” leaf of *N. benthamiana*. After 24 hours, the abaxial side of the next lower leaf beneath the local leaf was syringe-infiltrated with 100 µL of a TMV suspension at a dose of 0.1 mg mL^-1^. The TMV particles were freshly extracted from infected tobacco by grinding plant material frozen in liquid nitrogen using a mortar and pestle. The final dose of TMV was adjusted by adding 10 mM phosphate-buffered saline solution (pH 7.4). On day-7 after TMV inoculation, viral infection symptoms were observed in the newly grown leaves at the plant apex. The titer of TMV in these young leaves was quantified spectrophotometrically (Multiskan GO, Thermo Fisher Scientific, Waltham, USA) using a double-antibody sandwich enzyme-linked immunosorbent assay (DAS-ELISA) TMV Fast Kit (LOEWE Biochemica GmbH, Sauerlach, Germany). Infection rates are calculated relative to the viral titer reached in control plants.

### 2.8. Transcriptome sequencing of chitosan-treated N. benthamiana

Transcriptional changes in 4-week-old *N. benthamiana* plants were evaluated after treatment with chitosan in two different groups, including plants before and after TMV inoculation, using RNA-Seq analysis. In the first group of plants, a middle “local” leaf was sprayed with chitosan 671-CsnMN at a concentration of 100 µg mL^-1^, followed by harvesting the treated leaf two hours post-treatment (2 hpt). The expression profile of these samples was compared to that of the control plants which had only received dH_2_O. In the second group, plants were treated as above and 24 h later, the plants were inoculated with TMV as described above, and their apex leaves were harvested seven days post-inoculation (7 dpi). Their expression level was normalized to the control groups (dH_2_O-treated plants inoculated with TMV). RNA was extracted from these samples using a commercial column-based innuPREP RNA Mini Kit (Analytik Jena Co, Germany), and its quantity and quality was assessed using a nanodrop UV-Vis 2000 spectrophotometer (ThermoFisher Scientific Co, Waltham, USA) coupled to a Bioanalyzer 2100 machine (Agilent Technologies, Santa Clara, USA).

### 2.9. RNA-Seq transcriptome analysis

RNA-Seq measurements were performed by collecting a minimum of 150 base pairs (bp), paired-end reads on one HiSeq lane using a NovaSeq 6000 system (Illumina, San Diego, USA) at Novagene Company, United Kingdom. After controlling the quality of the raw sequencing data, the abundance of transcriptomic data was aligned to the genomic reference database and analysed by the Salmon aligner (Patro et al., 2017). FastQC was used to check the quality of the raw sequencing data (Andrews, 2010). Adapter trimming was performed with Trim Galore! (Krueger, 2018). The *N. benthamiana* genome and the associated annotation file were obtained from http://plantregmap.gao-lab.org/download.php#comparative-genomics. To corroborate the analysis, data were aligned to the *N. attenuata* genome as the most similar reference genome to *N. benthamiana* (http://ftp.ensemblgenomes.org/pub/plants/release-52/gff3/nicotiana_attenuata/).

### 2.10. RNA-Seq data analysis

Data obtained from gene-level read counts and alignment process were imported to the R package DESeq^2^ software and summarized for normalization and differential gene expression analysis of each sample using the tximport method (Love et al., 2014). Visualization of the clustering of the samples was performed using principal component analysis (PCA) and a Pearson’s correlation heatmap analysis. The volcano plotting of the DEG analysis was performed using pheatmap and visualized using ggplot2 (Wickham, 2011). The RNA-Seq results of the experimental groups were compared and normalized to the control (dH_2_O instead of chitosan treatment). Genes exhibiting p-values of less than 0.05 and absolute log2-fold changes of higher than 1 were considered statistically significant differential expressions. Multiple data analyses, including gene ontology (GO), KEGG, and over-representation analysis (ORA), were performed to examine the correlation and accuracy of the obtained data and to define whether these pathways or functions are well associated with the DEG analysis based on enrichGO and enrichKEGG functions from the clusterProfiler packages. Also, a second GO analysis was performed using the Plantregmap database (https://plantregmap.gao-lab.org/) to further confirm the results obtained using the R script. Lastly, the cnetplot function was collected from the enrich-plot R package to generate the concept network data.

### 2.11. RNA-Seq validation using quantitative reverse transcription PCR (RT-qPCR)

Next, to validate the RNA-Seq results, RT-qPCR was performed. Complementary DNA was synthesized based on total RNA (500 ng) using a commercial PrimeScript RT-Master Mix Kit (Takara Bio Inc, Kusatsu, Japan). A list of the primers used for RT-qPCR analysis is given in **Supplementary Table S1**. Glyceraldehyde 3-phosphate dehydrogenase (GAPDH) was used as a reference housekeeping gene. Five µL of a commercial KAPA SYBR FAST qPCR Master Mix Kit (Sigma-Aldrich, St. Louis, USA) was added to 2.5 µL of a 1:50 mixture of cDNA and dH_2_O and 2.5 µL of a 1:250 mixture of primer-pair and dH_2_O. RT-qPCR was conducted using a CFX96 Touch Real-Time PCR Detection System (Bio-Rad Inc, Hercules, USA). RT-qPCR data were quantified using the 2^-ΔΔCt^ method.

### 2.12. Statistical analysis

All data are presented as mean ± standard deviation. Statistical analysis was done using one-way ANOVA followed by Tukey’s post-hoc multiple comparison tests, and two-tailed T-tests between two groups unless stated otherwise in the figure legends. Data were analysed using GraphPad PRISM versions 8.0 and 9.02 (GraphPad Software, Inc., San Diego, California, USA). Outlier data of antiviral measurements were identified using the standard ROUT outliers test (Q = 0.1). Asterisks show statistically significant differences between the experimental groups (*P < 0.05, **P < 0.01, ***P < 0.001, ****P < 0.0001), “ns” indicates “not significant”.

## 3. Results and Discussion

### 3.1. Preparation and characterization of chitosan hydrolysates

A chitosan polymer (coded “651”) with *F*_A_ of 0.2 (DP ca. 800, *Ð* ca. 1.5) was used as a substrate for producing hydrolysates containing paCOS mixtures with different PA. As described in more detail below, we enzymatically hydrolysed the parent chitosan using recombinant chitosanase and chitinase enzymes from different glycosyl hydrolase (GH) families as defined by the Carbohydrate Active Enzymes (CAZy) database and compared the products (coded “671”) with those of acid hydrolysis using hydrochloric acid (coded “661-Cl”). In detail, the GH8 chitosanase CsnMN from *Bacillus* spec. with the cleavage specificity-D-D | X-X-(Weikert et al., 2017)(Weikert et al., 2017), the GH18 chitinase TvChi from *Trichoderma virens* with the cleavage specificity X-A | X-X (Bußwinkel et al., 2018)(Bußwinkel et al., 2018) and the GH19 chitinase ChiG from *Streptomyces coelicolor* A3 with the cleavage specificity-A-X | A-X-(Heggset et al., 2009b)(Heggset et al., 2009b) were used to create three hydrolysates named “671-CsnMN”, “671-TvChi”, and “671-ChiG”, respectively. A schematic model depicting the preparation process of the hydrolysates from chitosan polymer 651 is given in **Figure 1** above.

First, we separated the hydrolysates according to size using semi-preparative SEC. All hydrolysates were characterized by a broad range of chitosans from small polymers to small oligomers (**Figure 2**). While the acid hydrolysate 661-Cl was dominated by polymers, the oligomeric fractions were more pronounced in the enzymatic 671 hydrolysates which all showed a strong over-representation of paCOS with DPs of a multiple of 3, i.e. DP 3, 6, 9, etc. As shown recently (Hellmann et al., 2024), this effect is caused by the subsite specificity of the enzymes in combination with the typical, more regular than random PA of the chitosan polymer generated by the heterogeneous deacetylation process used for its production from chitin. At the same time, the missing of such an overrepresentation in the acid hydrolysate shows that under the conditions used, the hydrochloric acid hydrolyzed the glycosidic linkages of GlcN and GlcNAc at about equal rates, leading to random-PA paCOS. Overall, the composition of the chemical and enzymatic hydrolysates of the former (Richter et al. 2025) and of the current study are very similar, confirming the reproducibility of the processes and suggesting that under the conditions used, acetic and hydrochloric acid hydrolysis yield similar products.

**Figure 2:**
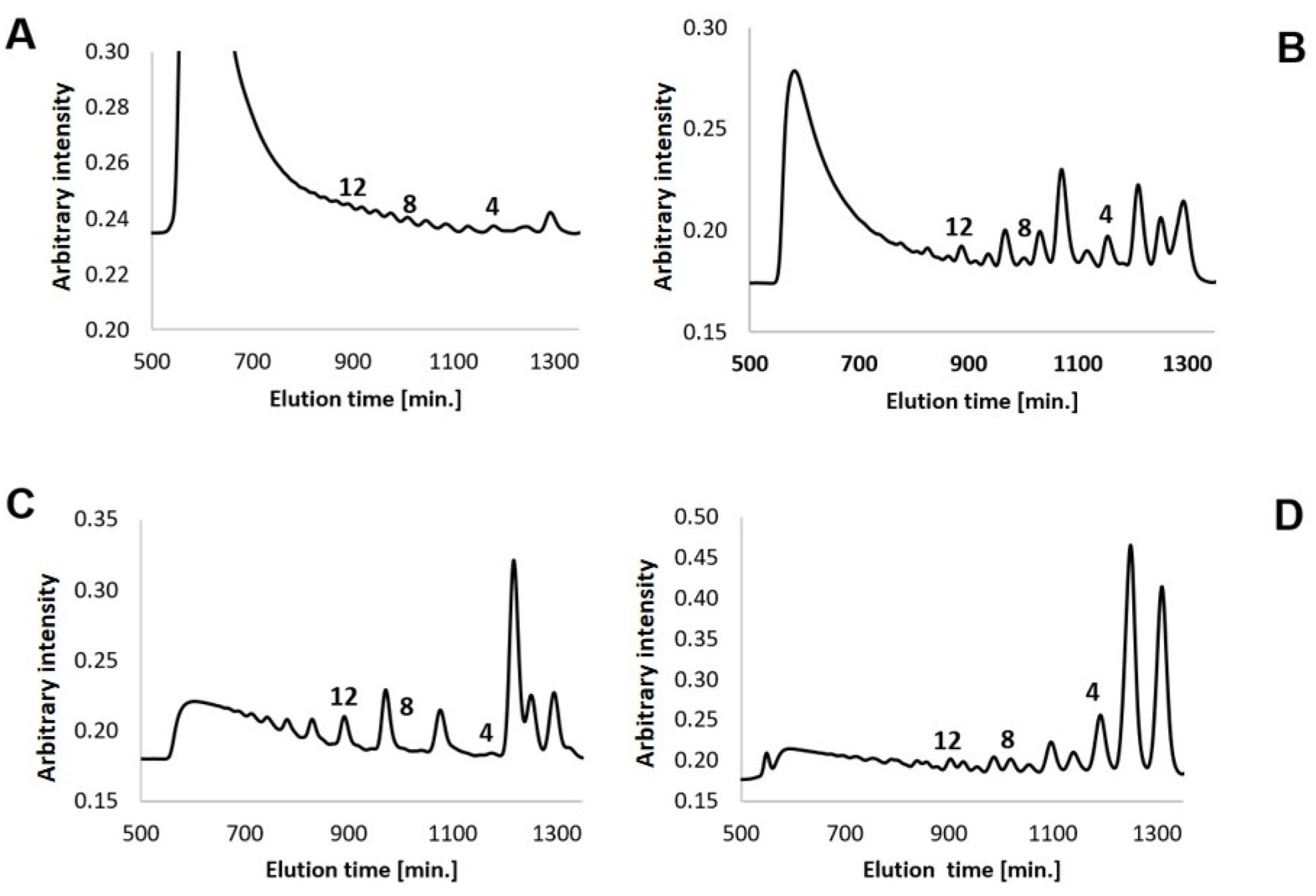
Size-exclusion-chromatograms of A) 661-Cl, B) 671-Tv Chi, C) 671-ChiG, and D) 671-CsnMN chitosan hydrolysates. The samples were separated by size exclusion chromatography and detected by a refractive index detector. The enzymatic hydrolysates (B-D) clearly show the overrepresentation of oligomers with DPs 3, 6, 9, 12, etc. brought about by the more regular than random PA generated by the heterogeneous alkaline deacetylation process used for the production of the chitosan polymer that served as a substrate for the hydrolyses. The position of oligomers purified and used in further experiments (DP 4, 8, 12) are indicated in the chromatograms.

Mass spectrometric analysis of the oligomeric products revealed the highest complexity in terms of different paCOS in the acid hydrolysate 661-Cl, followed by 671-TvChi (**Supplementary Figure S1**). While the GH18 chitinase TvChi has only one subsite (-1) with an absolute specificity for GlcNAc, the GH19 chitinase ChiG has two (-2 and +1) sites with very high preference for GlcNAc, and the GH8 chitosanase CsnMN has two sites with very high preference for GlcN. Consequently, the composition of the latter two enzymes’ products is less complex. The broad diversity of oligomers seen in the hydrochloric acid hydrolysate 661-Cl is similar, though not identical, to that seen in an acetic acid hydrolysate (661-Ac) of the same chitosan polymer (Richter et. al 2025). Similarly, the oligomers in the GH8-chitosanase hydrolysates 671-CsnMN and in the GH18-chitinase hydrolysates 671-TvChi, which were prepared independently for both studies, have similar but not identical compositions in both studies. In the current study, we also included a GH19-chitinase to produce 671-ChiG which we had not used in the previous study and which gave the least complex oligomer mixture.

This difference in complexity caused by the specificity of each enzyme reflects the level of definition in PA of the paCOS produced: the higher the substrate preferences of the enzyme’s subsites, the less complex the composition, and the more defined the paCOS produced, as shown schematically in **Figure 1B**. CsnMN with subsite preferences-D-D | X-X-mainly produces oligomers with two GlcN units at their reducing ends. TvChi with subsite preferences-X-A | X-X-mainly produces oligomers with a GlcNAc unit at their reducing end. ChiG with subsite preferences-A-X | A-X-mainly produces oligomers with GlcNAc units at their non-reducing end and next to their reducing end (Hellmann et al., 2024).

In the former study (Richter et al. 2025), we had separated an oligomeric fraction containing oligomers of DP 1-15 from a polymeric fraction of DP >15. To obtain deeper insight into the DP-dependency of the oligomers concerning the bioactivities tested, we now collected the fractions containing oligomers of DP 4, 8, and 12 from each hydrolysate, and analyzed them by mass spectrometry. DP 8 was chosen as the optimum DP for ligand binding to the chitin receptor CERK1 (T. Liu et al., 2012; Vander et al., 1998), DP 4 because of the known bioactivities of chitin (Winkler et al., 2017)and monoacetylated chitosan (Basa et al., 2020) tetramers, and DP 12 because of the increasing elicitor activities of partially acetylated chitosan oligomers with DP even beyond DP 8 (Basa et al., 2020; Gubaeva et al., 2018). The MS chromatograms showed the successful separation of oligomers according to their DP and confirmed that they comprise mixtures of paCOS with different *F*_A_ (**Supplementary Figure 2A-D**). Due to the different cleavage specificities, the paCOS are also characterized by specific PAs, as described above.

### 3.2. Eliciting activities of the chitosan hydrolysates

An oxidative burst is the rapid and transient accumulation of reactive oxygen species (ROS) such as hydrogen peroxide (H_2_O_2_). In plants, the oxidative burst is an early hallmark of defence responses against pathogens and it can also be elicited by pathogen-derived compounds (Bi et al., 2022; Kaur et al., 2021). Some bioactive chitin and chitosan oligomers possess the innate ability to bind to specific plant cell surface receptors and elicit resistance responses in plants, including the oxidative burst (Levine et al., 1994; Mittler, 2017). We evaluated the eliciting activity of the different hydrolysates and paCOS described above and compared them with that of the parent chitosan polymer 651 by performing an oxidative burst assay in *N. benthamiana* leaf discs (**Figure 3**A and **3**B) and seedlings (**Figure 3**C and **3**D).

**Figure 3:**
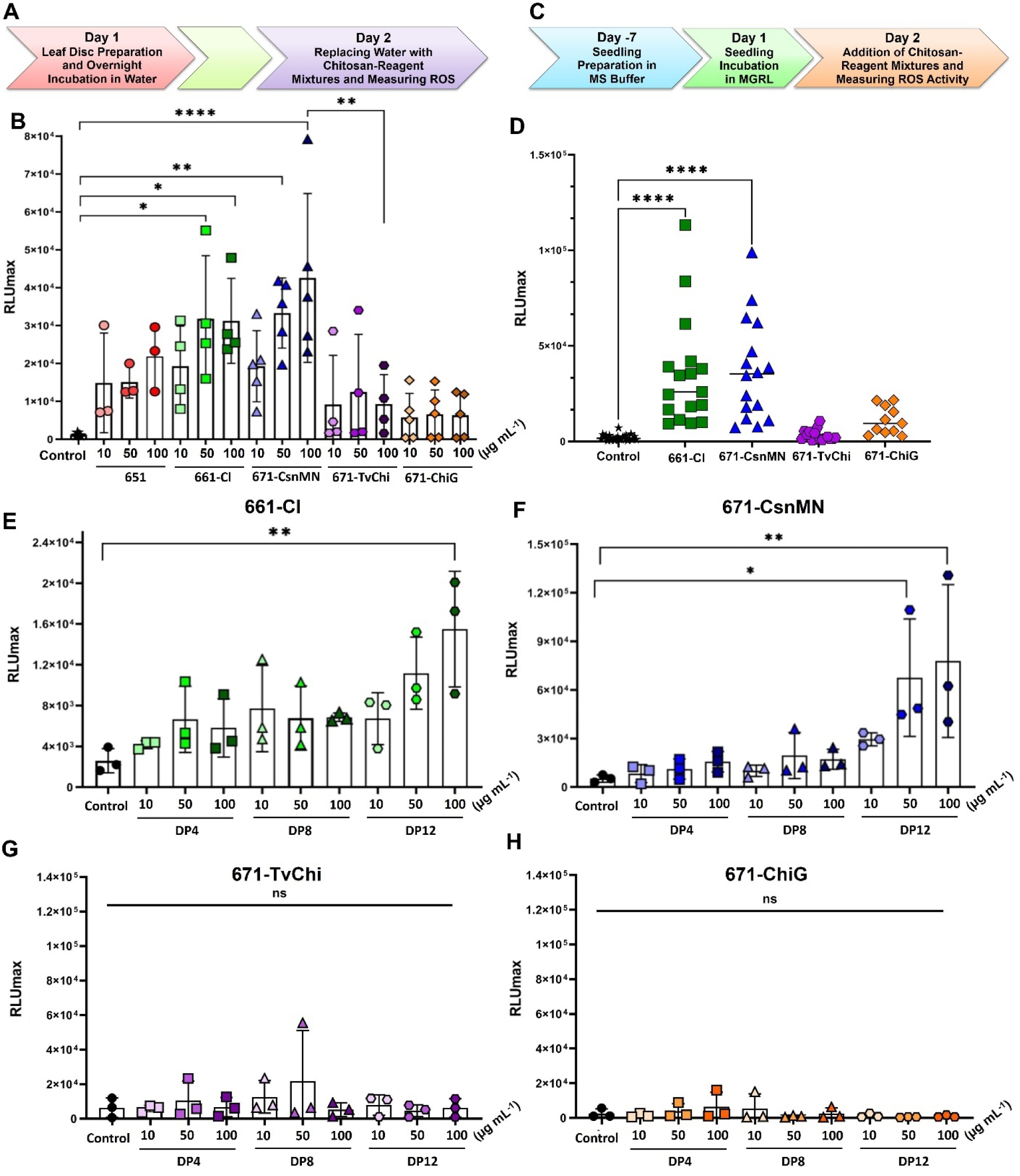
Elicitor activities of the chitosan polymer (651), its acid (661) and enzymatic (671) hydrolysates, and their oligomers with different DP in *N*. *benthamiana*. A) Timeline of the eliciting assay in leaf discs. B) Leaf discs were treated with different concentrations of chitosans or ddH_2_O as a control, and the production of ROS was measured as relative light units (RLU) in the medium continuously over 90 min using a luminol-based bioassay. The bars represent the means ± SD of the maximum values reached, individual data points give the means of the independent replicates performed (N = 3-5, n = 6-9). C) Timeline of the eliciting assay in seedlings. D) Oxidative burst measurements of the samples at the concentration of 10 µg mL^□1^, compared to control groups (ddH_2_O) (N = 3, n = 6). E-H) Effect of DP on the oxidative burst production in the leaf discs system; E) 661-Cl, F) 671-CsnMN, G) 671-Tv Chi and H) 671-ChiG (N = 3, n = 6-9).

In the leaf disc assay, the chitosan polymer 651 showed a dose-dependent tendency to elicit an oxidative burst which, however, was statistically not significant (**Figure 3B**). In contrast, the chemical hydrolysate 661-Cl showed clear elicitor activity. The chitosanase hydrolysate 671-CsnMN showed even stronger elicitor activity, while the chitinase hydrolysates 671-TvChi and 671-ChiG were almost elicitor-inactive. These results corroborate and extend our earlier observations on elicitor activities of GH8-and GH18-hydrolysates in seedlings of *Arabidopsis*. We assume that the observed strong differences in the eliciting activities of the enzymatic hydrolysates are largely due to their different PAs. As discussed above, chitosanase produces paCOS with GlcNAc units dominating their internal positions while their ends are dominated by GlcN units. This specific PA of chitosanase products may favor their binding to the plant’s chitin receptors, probably promoting receptor dimerization and ensuing elicitation of an oxidative burst. In contrast, the specific PA of chitinase products with central GlcN units flanked by GlcNAc units may prevent receptor binding and, thus, elicitor activity. To verify the results of the leaf disc assays, we tested the eliciting activity of the hydrolysates in intact *N. benthamiana* seedlings (**Figure 3D**). Again, the chemical hydrolysate 661-Cl and the chitosanase products 671-CsnMN possessed elicitor activity, while the chitinase products 671-TvChi and 671-ChiG did not significantly elicit ROS production, showing that the effects were not dependent on the wounding involved in the leaf disc assay.

We then tested the elicitor activities of the purified oligomers with DP 4, 8, and 12 using both assays (**Figure 3E-H** and **Supplementary Figure S3**). From the chemical hydrolysate, only the highest concentration of the DP 12 oligomers showed significant, but weak elicitor activity. The chitosanase-produced oligomers of DP 12 had much stronger activity, while the chitinase-produced oligomers were elicitor-inactive. In a direct comparison of the DP4 fractions, only the chitosanase-produced tetramer 671-CsnMN showed significant elicitor activity at the concentration of 10 µg mL^-1^; interestingly, this fraction contained mostly the fully deacetylated tetramer, but also a small amount of mono-acetylated tetramer(s) (Regel et al., 2018b) (**Figure 1D** and **Supplementary Figure S4**). Based on the cleavage specificity of the enzyme used, the latter are expected to be a mixture of ADDD and DADD (Basa et al., 2020). The mixtures of oligomers in the DP 8 and DP 12 fraction were clearly more complex, not allowing comprehensive sequencing, and with defined sequences only at and near their ends, as described above.

Clearly, the paCOS produced by chitosanase-catalyzed hydrolysis exhibited the strongest elicitor activities in tobacco. While the elicitor activity increased with increasing DP of the oligomers, as expected, even the tetramer fraction showed slight, but significant activity in the more sensitive seedling assay. These results are in agreement with our earlier work reporting the elicitor activity of a GH8-chitosanase hydrolysate of a *F*_A_ 0.35 random-PA chitosan and oligomers purified from it, but not of a GH18-chitinase hydrolysate of the same chitosan, in suspension-cultured rice cells (Basa et al., 2020). They also align with a parallel study on the elicitor activity of chitosan hydrolysates in the model plant Arabidopsis thaliana (Richter et al., 2025), extending the earlier findings i.a. regarding the DP-dependency of the effects.

### 3.3. Priming activities of the chitosan hydrolysates

In plant physiology and defence, priming is a sophisticated process which prepares plants to more quickly and more strongly respond to biotic and abiotic stresses(Andersen et al., 2018; Tilocca et al., 2020). Primed states can be triggered either by plant-beneficial microorganisms or by treatment with naturally existing or synthetic compounds (Laura et al., 2018). Priming and priming-enhanced defence responses have been described repeatedly in plants, but investigation of the underlying mechanisms is in its infancy (Conrath et al., 2002). As an example, chitosan treatment increased jasmonic acid levels and enhanced the expression of defence genes in tomato plants, promoting tolerance against *Botrytis cinerea* (Conrath et al., 2015; de Vega et al., 2021). In our parallel study, we had shown that elicitor and priming activities in *Arabidopsis* were unchanged by acetic acid, destroyed by GH18-chitinase, and increased by GH8-chitosanase hydrolysis, using an oxidative burst assay in seedlings. Increased ROS generation in primed compared to naïve plants upon elicitor treatment is regarded as a standard marker for measuring priming activity (Pastor et al., 2013; Roychoudhury et al., 2022; Srivastava et al., 2009).

Therefore, we assessed the priming activity of the chitosan samples in three different bioassays, eliciting an oxidative burst in chitosan-primed *N. benthamiana,* using flagellin (flg22) as an elicitor. **Figure 4A** gives a schematic representation of the priming assay using intact plants. Briefly, a single middle-aged leaf of the plant was sprayed with the chitosan solution or water as a control and discs from the treated leaf (local response) and from an upper, younger leaf (systemic response) were collected 2 h after the treatment. One day later, the leaf discs were treated with flg22, and H_2_O_2_ production was measured immediately. As can be seen in **Figure 4B**, the elicitor-active chitosanase hydrolysate 671-CsnMN had the strongest local priming activity and, surprisingly, also the elicitor-inactive GH18 chitinase hydrolysate 671-TvChi showed significant, though less pronounced priming activity locally. In line with their low or missing elicitor activities shown above, the chemical hydrolysate 661-Cl and the GH19 chitinase hydrolysate 671-ChiG had no significant local priming activity. None of the hydrolysates exhibited significant systemic priming activity (**Figure 4C**). As the locally priming-active chitosan hydrolysate 671-CsnMN also exhibited the strongest tendency towards systemic priming, we also analysed the local and systemic priming activities of the purified DP 4, 8, and 12 oligomers of this fraction. Both the octamer and the dodecamer significantly induced priming both locally (**Figure 4D**) and systemically (**Figure 4E**) at appropriate concentration (100 µg mL^□1^).

**Figure 4:**
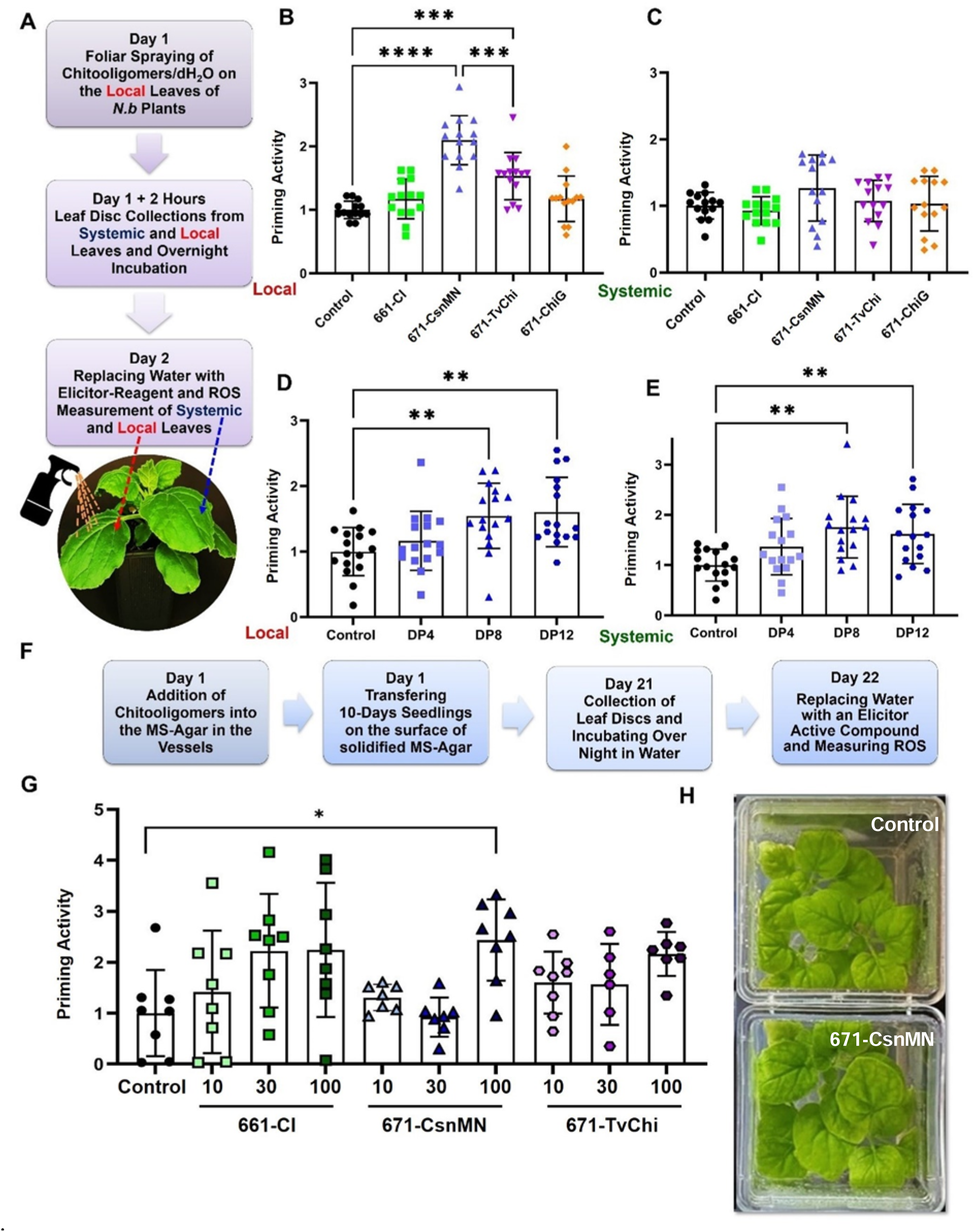
Priming activities of the acid (661) and enzymatic (671) chitosan hydrolysates and their oligomers with different DP in *N*. *benthamiana.* A) Timeline of the priming activity assessment in *N. benthamiana* plants. Priming response of the hydrolysates at a concentration of 100 µg mL^□1^ on B) local and C) systemic leaves was evaluated after foliar spraying chitosans on the local leaf. D, E) Influence of DP on the priming activity of chitosan oligomers SEC purified from the 671-CsnMN hydrolysate at 100 µg mL^□1^ (N = 3, n = 3-4). F) Experimental timeline of the priming activity of the hydrolysate-treated culture box *N. benthamiana* plants. G) Assessments of the priming activity of the hydrolysates at the concentrations of 10, 30, and 100 μg mL^□1^. (N = 1, n = 8). H) Optical observation of the representative 3-week plants inside the culture boxes (top: control and bottom: 671-CsnMN at 100 µg mL^□1^). In all cases, the oxidative burst was triggered by flg22 as an elicitor-active compound. Data are normalized to the control plants sprayed with water instead of chitosan, and presented as mean ± SD.

In an independent bioassay, 10-day-old tobacco seedlings were transferred to chitosan-containing agar, and leaf discs were prepared three weeks later followed by flg22-elicitation another day later (**Figure 4F**). All the hydrolysates tested (671-ChiG not tested) showed a tendency towards dose-dependent priming, but only the priming by the chitosanase hydrolysate 671-CsnMN was significant in this assay (**Figure 4G, H**). In a third, microtitre-plate-based priming bioassay, intact 6-day-old tobacco seedlings were treated with the 671-CsnMN hydrolysate or its purified oligomers 24 h prior to flg22-elicitation and H_2_O_2_ quantification, where the total hydrolysate showed significant priming activity while the increased oxidative bursts in the DP4, DP8, and DP12 oligomer-treated seedlings were statistically not significant (**Supplementary Figure S5**).

Again, these results corroborate and extend our studies in suspension-cultured rice cells (Basa et al., 2020) and in the model plant *Arabidopsis* (Richter et al. 2025). Importantly, the use of different bioassays showed that also priming activities can be observed in different assays, i.e. are independent of e.g. wounding which is inherent in the leaf disc assay.

Interestingly, when we assessed the root growth of the plants grown in chitosan containing agar, the chitosan polymer 651 as well as the chitinase hydrolysates 671-TvChi and 671-ChiG led to increased dry weights (although statistically not significant), while the chemical hydrolysate 661-Cl and the chitosanase hydrolysate 671-CsnMN did not (**Supplementary Figure S6**).

### 3.4. Resistance-inducing activities of the chitosan hydrolysates against viral disease

The studies on elicitor and priming activities described above and in the previous studies suggest that the chitosanase hydrolysate 671-CsnMN, but not the chitinase hydrolysate 671-TvChi, may have the potential to protect plants from disease. To experimentally test this hypothesis, we next investigated the ability of the chitosan polymer 651 and its different hydrolysates to systemically induce resistance in *N. benthamiana* against TMV. The chitosan solutions were sprayed onto a single, middle-aged leaf of a tobacco plant and one day later, a lower, older leaf was inoculated with TMV. Symptoms of viral infection were assessed five days later at the apex of the plants, and another three days later, the uppermost, youngest leaves were harvested for quantification of the viral titre (**Figure 5A**, B and **Supplementary Figure S7**). As expected from the elicitor and priming assays described above, the chemical hydrolysate 661-Cl exhibited a weak and the chitosanase hydrolysate 671-CsnMN a strong resistance-inducing activity, while both chitinase hydrolysates, 671-TvChi and 671-ChiG, were as inactive in this assay as was the parent chitosan polymer 651 (**Figure 5C**). Interestingly, we observed an all-or-none effect in all treatments: individual plants were either fully resistant or fully susceptible to the virus, the difference between successfully treated plants and control plants or plants treated with inactive chitosans lay in the relative number of infected *versus* healthy plants, i.e. chitosan treatment reduced disease incidence, but not disease severity. When we tested the paCOS mixtures with DP 4, 8, and 12 of the chitosanase hydrolysate 671-CsnMN individually, all reduced the infection ratio, but the inhibition was significant only for the DP 12 oligomers (**Figure 5D**). Taken together, these results clearly show that the chitosanase hydrolysate 671-CsnMN has a proven plant disease protecting effect.

**Figure 5:**
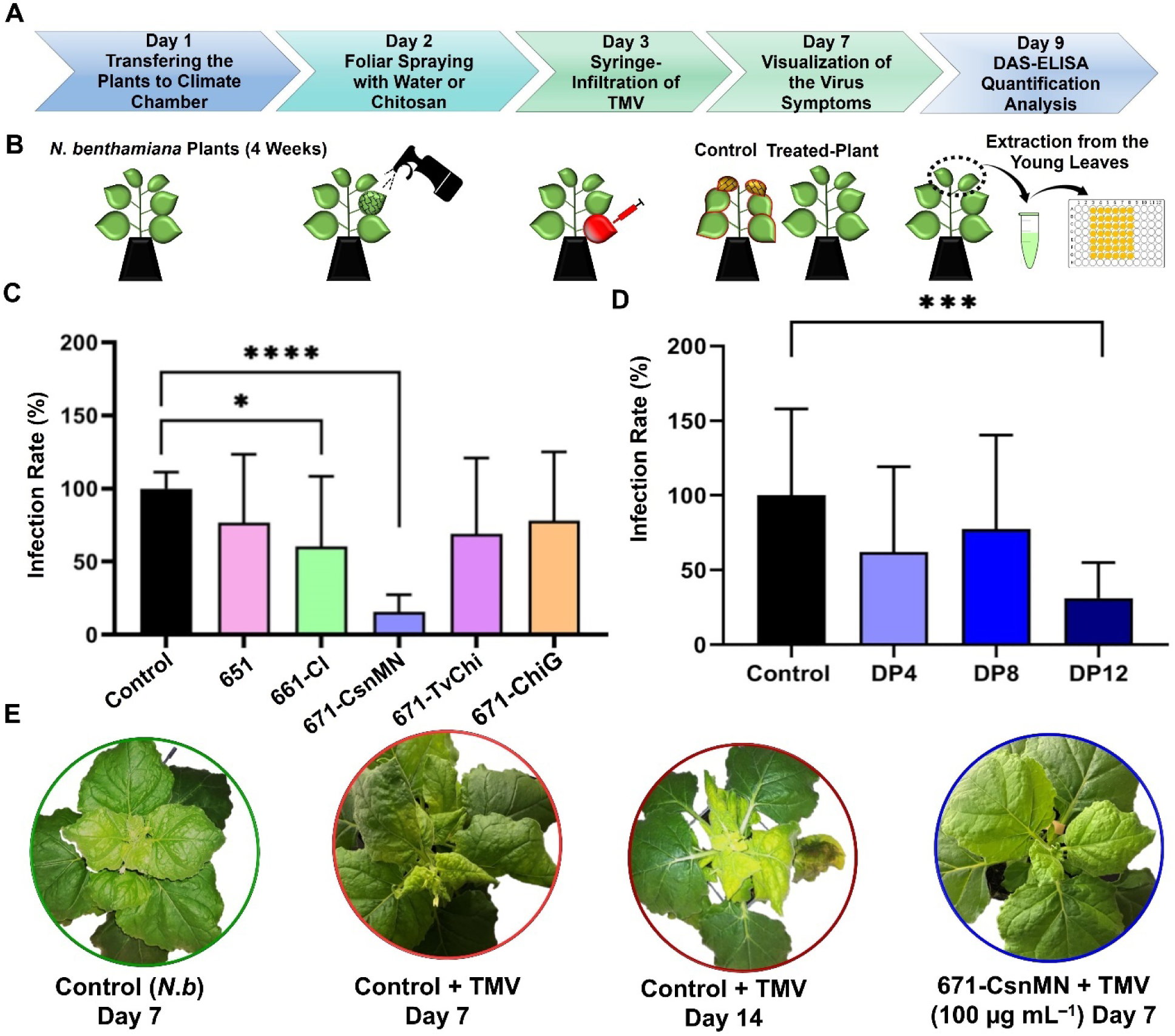
Antiviral activities of the chitosan polymer (651), its acid (661) and enzymatic (671) hydrolysates, and their oligomers with different DP in *N*. *benthamiana*. A) Timeline and B) schematic illustration of the antiviral assay. C) Viral infection in plants treated with different chitosan solutions at a concentration of 100 μg mL^−1^ (N = 2-3, n = 4-6), determined by quantifying the viral titer in the apex leaves using a DAS-ELISA kit for TMV. D) Influence of DP on the antiviral properties of chitosan oligomers SEC-purified from 671-CsnMN hydrolysate at 100 μg mL^−1^ (N = 3, n = 3-6). E) Optical appearance of the control, TMV-inoculated, and 671-CsnMN-treated as well as TMV-inoculated plants 7 or 14 days after treatment. Data are presented as the mean ± SD.

### 3.5. Transcriptomic analysis of treated and non-treated plants

To gain a more in-depth insight into the response of tobacco plants to treatment with the most strongly elicitor-and priming-active as well as resistance-inducing chitosanase hydrolysate, 671-CsnMN, we analyzed the influence of chitosan-treatment and TMV-inoculation on gene expression in the treated, local leaf and in the plant apex, respectively. In a preliminary experiment, we examined the expression of selected defense-related genes coding for pathogenesis-related protein 10 (PR10), nonexpressor of pathogenesis-related protein 1 (NPR1), enhanced disease susceptibility 1 (EDS1), respiratory burst oxidative homolog B (RBOHB), nitrate reductase (NR), and N requirement gene 1 (NRG1) 2 and 24 h after chitosan treatment in the treated, local leaf (**Supplementary Figure S8**). We found a slight up-regulation (log2-fold change ca. 1) of RBOHB and down-regulation (log2-fold change ca.-1) of NR only 2 h, but not 24 h after chitosan treatment. Based on these results, we performed a comprehensive RNAseq analysis in *N*. *benthamiana* plants 2 h after treatment with the chitosan hydrolysate CsnMN or water as a control in the treated leaf, to reveal genes involved in triggering systemic resistance induction. A second timepoint for RNAseq analysis was chosen 7 d after TMV-inoculation of chitosan-or water-pretreated plants in the plant apex, to reveal genes involved in performing the systemic resistance reaction. Principal component analysis (PCA, **Supplementary Figure S9**A) and Pearson’s correlation coefficient analysis (**Supplementary Figure S9**B) indicated that these treatment groups were well-separated into the defined clusters along the first principal component, which accounted for 72 % of the variance between these groups. Two hours after chitosan treatment, 525 genes were significantly up-regulated and 81 genes were significantly down-regulated in the chitosan-treated leaves compared to water-treated control plants (log2 ≥1 or ≤-1) (**Figure 6A**). One week after TMV inoculation, 1,466 genes were more strongly expressed in the apex of chitosan-pretreated and, thus, resistant plants (*nota bene:* only plants visibly resistant were harvested for RNAseq analysis) than in water-pretreated and, thus, susceptible control plants, while 3,060 genes were more strongly expressed in the susceptible than in the resistant plants (**Figure 6C**).

**Figure 6:**
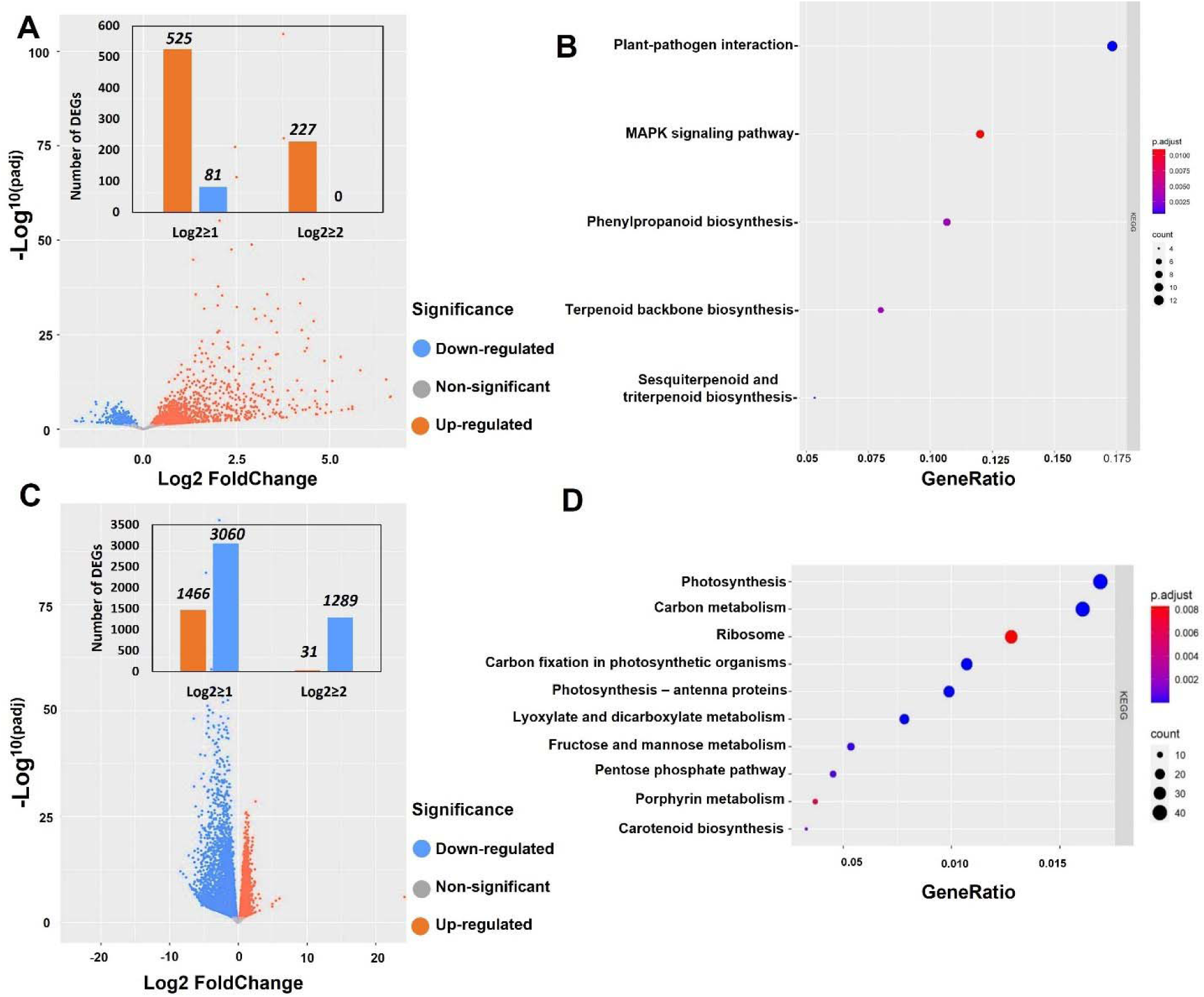
DEG analysis of *N. benthamiana* plants treated with 671-CsnMN chitosan hyrolysate at a concentration of 100 µg mL^□1^ compared to dH_2_O-treated control plants. Plants were foliar-sprayed with the chitosan solution in two different groups, without and with TMV inoculation, and harvested 2 hpt (chitosan-treated leaf) or 7 dpi (apex leaves), respectively. Volcano plots of the DEGs identified A) 2 hpt and C) 7 dpi. DEGs with *p-*values below 0.05 and log2 >1 (up-regulated genes) or log2 < 1 (down-regulated genes) were considered statistically significant. KEGG over-representation analysis (ORA) of the up-regulated genes B) 2 hpt and D) 7 dpi. the *p*-value of false discovery rate (FDR) was set to <0.05 with log2 ≥1. Custom R scripts (enrichKEGG functions from the clusterProfiler package) were used to calculate the ORAs of each sample. The defined dominant pathways are aligned to the annotation database of *N. benthamiana* genes. The rich factor of the genes is color-coded (blue to red), and the size of dots indicates the number of DEGs.

Next, we performed a Kyoto encyclopedia of genes and genomes (KEGG) analysis to identify the functional pathways affected by 671-CsnMN-treatment and TMV-inoculation. As the KEGG pathway analysis of *N. benthamiana* is unavailable online, we used the genome database of *N. attenuata* as the most similar genotype in this family (Pandey et al., 2008; Schiavinato et al., 2020). A comparison of the DEGs of *N. benthamiana* and *N. attenuata* is given in **Supplementary Figure S10** for the genes which are up-regulated by chitosan treatment 2 h post treatment in the local leaf. These genes belong to five different KEGG pathways, including plant-pathogen interaction, mitogen-activated protein kinase (MAPK) signalling pathway, phenylpropanoid biosynthesis, terpenoid backbone biosynthesis, and sesqui-and triterpenoid biosynthesis, (**Figure 6B**). Clearly, chitosan treatment induced secondary plant metabolism. In contrast, the ten KEGG pathways identified as being more strongly expressed in chitosan-pretreated, resistant plants compared to water-pretreated, susceptible plants one week after TMV-inoculation all belong to primary metabolism, most prominently to photosynthesis and carbon metabolism, which apparently is down-regulated or collapsing during TMV infection (**Figure 6D**).

We also used the gene ontology (GO) annotation of the DEGs of *N. benthamiana* in the 2 hpt and 7 dpi samples, using the programme R. Thirty Biological Processes (BP) and twelve Molecular Functions (MF) were enriched by chitosan-treatment 2 hpt (**Figure 7A**). The most dominant BPs belong to the terms “defense response” and “response to biotic stimulus”, while “sequence-specific DNA-binding” and “heme-binding” are identified as the main MFs. When the GO analysis was performed on the 2 hpt sample using the Plantregmap database, the up-regulated DEGs (log2 ≥2) belong to 97 GO terms, including 71 BFs, 23 MFs, and 3 Cellular Components (CC) (**Supplementary Figure S11**), in good agreement with the first GO analysis performed using R. The genes induced most strongly by chitosan-treatment were Patatin-like protein 2 ((Niben101Scf06277g00007), Somatic Embryogenesis Receptor Kinase (SERK) (Niben101Scf00821g16001), and Terpene Synthase-5 (Niben101Scf01683g03005). We were particularly interested in receptor like kinases, several of which were induced by chitosan treatment, including the chitin receptor CERK1, suggesting that this might be involved in the perception of the chitosan oligomers released by the chitosanase CsnMN, as shown previously in *Arabidopsis* (Miya et al., 2007). A detailed list of DEGs is presented in **Supplementary Table S2**.

**Figure 7:**
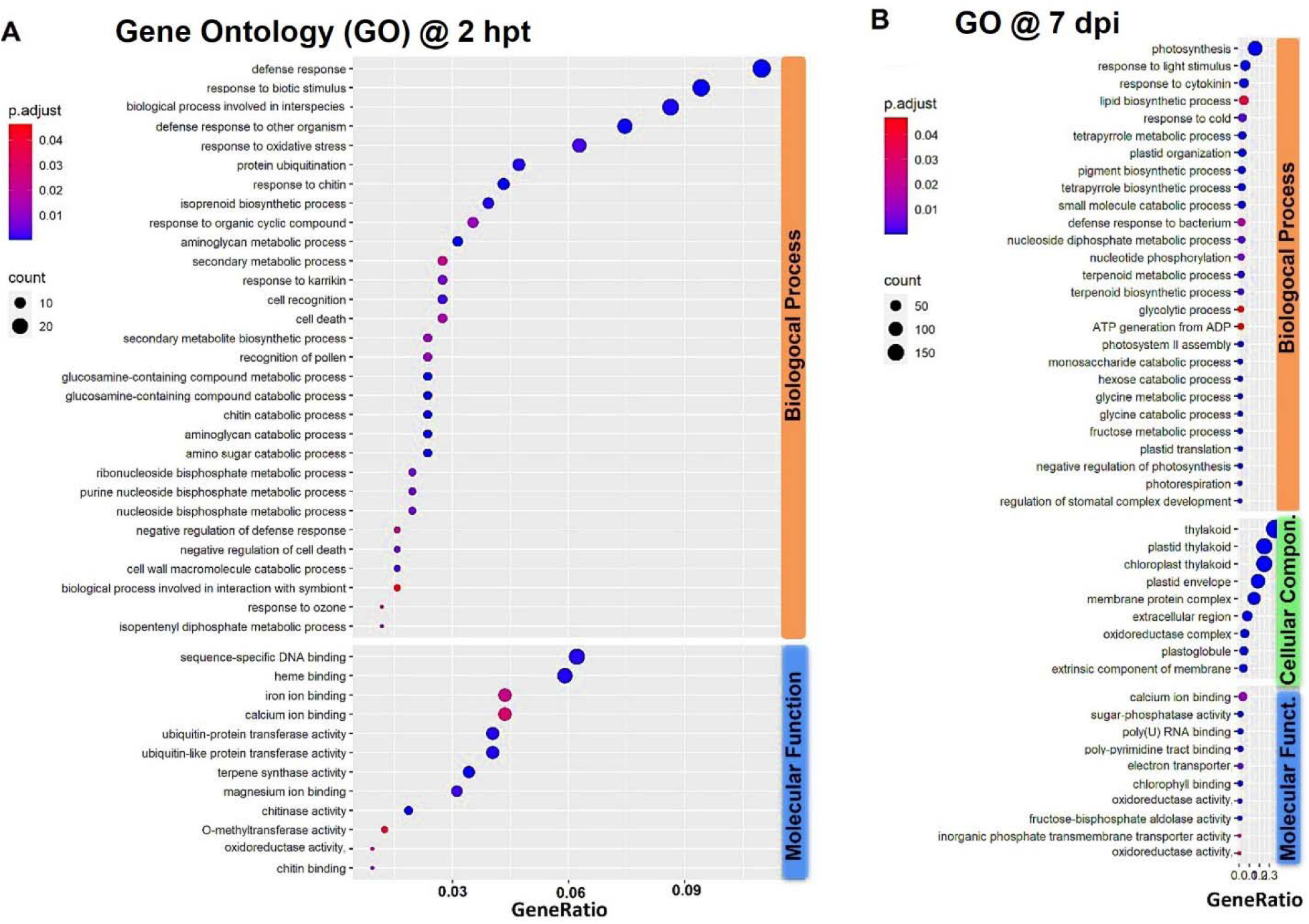
Gene ontology over-representation analysis of *N. benthamiana* plants treated with 671-CsnMN chitooligomer mixture at a concentration of 100 µg mL^□1^. Plants were foliar-sprayed with the chitooligomers in two different groups, A) without and B) with TMV inoculation, and harvested 2 hpt or 7 dpi, respectively. Each group contains different sub-ontologies, including biological processes, molecular functions, and cellular compounds. The statistically significant GO terms are aligned to the enriched relative background annotation of the *N. benthamiana* genes and listed in the y-axis of this plot. The size and color of the dots on the x-axis indicate the number of genes and the degree of rich factor, respectively (FDR <0.05 and log2 ≥1).

In the apex of the TMV-inoculated plants 7 dpi, the DEGs that were more highly expressed in the CsnMN-treated and, thus, resistant plants belong to 27 different BPs, 10 MFs, and 9 CCs. The most dominant enriched BPs are “photosynthesis” and “response to light stimulus”, while “thylakoid” and “calcium ion binding” are the main DEGs in the groups of CC and MF, respectively (**Figure 7B**). Again, these data agree with the GO analysis using the Plantregmap database where the DEGs (log2 ≥1.2) belonged to 31 GO terms, including 12 BFs, 14 MFs, and 5 CCs (**Supplementary Figure S12**). A detailed list of DEGs is given in **Supplementary Table S3** and **Supplementary Figure S13**.

We validated the accuracy of the RNAseq data using RT-qPCR, selecting ten putatively upregulated DEGs related to defense and photosynthesis (**Figure 8A**, **Supplementary Figures S14** and **S15**). In all cases, RT-qPCR verified the RNAseq results.

**Figure 8:**
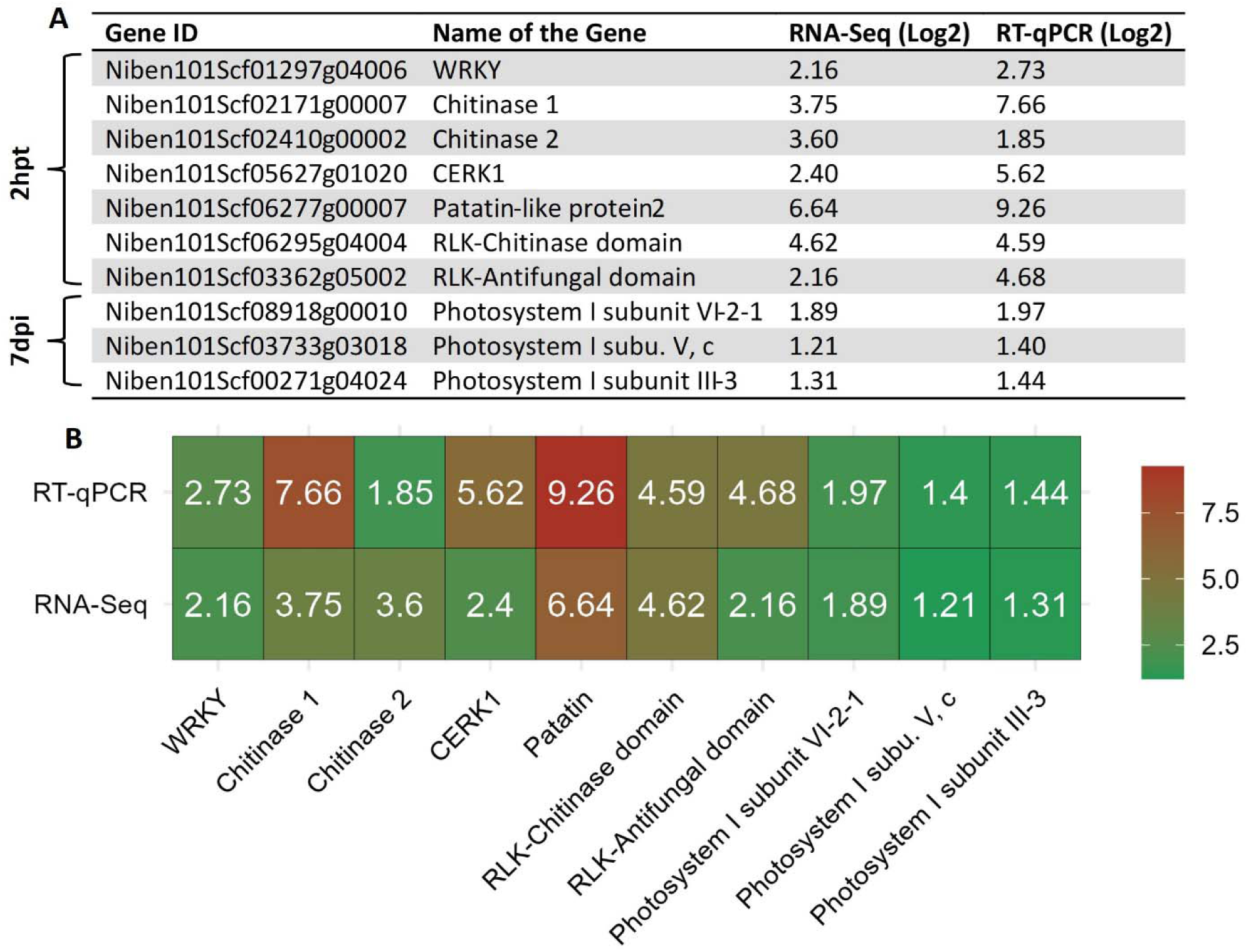
Validation of transcriptomic data obtained using RNA-Seq by RT-qPCR. A) Gene expression levels of selected DEGs. B) Color-coded heat map of RNA-Seq (N = 1, n = 3 pooled from three plants) and RT-qPCR (N = 1-2, n = 3-6) data. The two datasets are in agreement with each other (*p* = 0.0748).

### 3.6. Potential defence mechanisms induced by 671-CsnMN

To gain further insight into the plant’s transcriptomic responses to chitosan treatment and TMV infection, we performed gene concept network (Cnetplot) analyses (**Supplementary Figures S16** to **S23**) which reveal the linkages between DEGs and enriched KEGG or GO pathways. These data provide multiple valuable clues for further research, but we will highlight only selected aspects here. We found the known chitin/chitosan binding receptor CERK1 to be upregulated two hours after 671-CsnMN treatment, suggesting its involvement in the recognition of the chitosanase-produced chitosan oligomers. In addition to CERK1, several other protein kinases were also induced which, consequently, might also be involved in chitosan perception or chitosan-induced signal transduction (**Supplementary Figure S24)**. Also, other genes involved in ethylene-triggered, calcium, and MAPK signaling as well as WRKY transcription factors and three different chitinases were induced, indicating the elicitation of PAMP-triggered immunity (PTI).

Analysis of the enriched GO-terms of the 671-CsnMN-treated plants seven days after TMV inoculation mainly showed a massive breakdown of primary metabolism in the water-treated, susceptible control plants compared to the chitosan-treated, resistant plants. Interestingly, we found no induction of disease resistance-related terms in these samples which were taken from the apex of the plants, perhaps suggesting that chitosan-induced resistance prevented the virus from reaching the apex.

## 4. Conclusion

While there is ample evidence in literature that chitosans and chitosan oligomer mixtures can have plant protective activities, we still have only a very limited understanding of structure-function relationships of these bioactivities (Cord-Landwehr et al., 2020; Wattjes et al., 2020). In particular, the influence of the production route on the structure and function of chitosan oligomers have rarely been studied (Basa et al., 2020; Richter et al., 2025). Here, we clearly show that chitosan-hydrolyzing enzymes with different cleavage specificities can generate chitosan oligomer mixtures with different, partially defined acetylation patterns, and these differ strongly in their biological activities. Hydrolysates produced by chitinase are elicitor-and priming-inactive and do not protect tobacco plants from viral infection, presumably because these enzymes degrade bioactive GlcNAc-rich stretches within the chitosan polymer. In contrast, hydrolysates produced by chitosanase which leaves these GlcNAc-rich stretches intact have even stronger phytostimulatory activities than the original chitosan polymer. Importantly, the elicitor-and priming-active chitosanase products were able to protect tobacco plants from viral infection, while the elicitor-and priming-inactive chitinase products were not, suggesting a strong correlation between these activities, so that simple bioassays for elicitor/priming activity may be used to screen for complex plant protective properties. Seen together with the results of our parallel study on antibacterial and antifungal, elicitor and priming, as well as gene induction activities of chitosan hydrolysates in *Arabidopsis*, our study clearly shows that the partial PA-control that can easily be obtained by enzymatic hydrolysis - provided a structurally well-characterized chitosan polymer is used as a substrate and a functionally well-characterized and highly pure enzyme is used for hydrolysis - suffices to produce easily water-soluble chitosan oligomers with excellent and distinct biological activities. The observation that the hydrolysate obtained by the GH8 chitosanase CsnMN, which shows the highest activities in all phytostimulatory assays, also most strongly induces resistance against viral infection suggests that this approach is well-suited for the development of reliable agro-biologics.

## Data Availability

All raw and processed data are available from the corresponding authors upon reasonable request.

## Authorship Contribution

**S.K.**: Conceptualization, Methodology, Formal Analysis, Validation, Visualization, Writing – Original Draft, Writing – Review & Editing. **R.S.**: Methodology, Formal Analysis, Review & Editing. **J.R.**: Formal Analysis, Review & Editing. **N.Y.N.C.**: Formal Analysis, Review & Editing. **S.C.-L.**: Formal Analysis, Review & Editing. **C.R.**: Formal Analysis, Review & Editing **A.R.**: Formal Analysis, Validation, Writing – Original Draft, Writing – Review & Editing. **B.M.M.**: Conceptualization, Investigation, Resources, Formal Analysis, Supervision, Writing – Original Draft, Writing – Review & Editing. All authors have read and agreed to its content.

## Supporting information

Electronic Supplementary Information

## Acknowledgements

This work was supported financially through a doctoral fellowship to Soofia Khanahmadi from German Academic Exchange Service (DAAD). Chitosans 651 and 661-Cl were kindly provided by Dr. Dominique Gillet (Gillet Chitosan, Plumaudan, France). We would like to thank Sabrina Stritzel for technical assistance, and Dirk Schmidt and Sascha Ahrens for plant cultivation and maintenance.

