## Supplementary material for "Immunoengineered Chitosanase-Produced Chitosan Oligomers for Elevating Plant Resistance to Viral Infection": Electronic Supplementary Information

*Main, Germany*

*^c^ Institute of Cardiovascular Sciences, St. Boniface Hospital Albrechtsen Research Centre, Department of Physiology and Pathophysiology, Max Rady College of Medicine, Rady Faculty of Health Sciences, University of Manitoba, Winnipeg, Manitoba R2H 2A6, Canada*

Content:

*Supplementary Figures S1 to S24*

*Supplementary Tables S1 to S3*

**Correspondence:**

Prof Dr Bruno M. Moerschbacher

Institute for Biology and Biotechnology of Plants

University of Münster

Schlossplatz 8, 48143 Münster, Germany

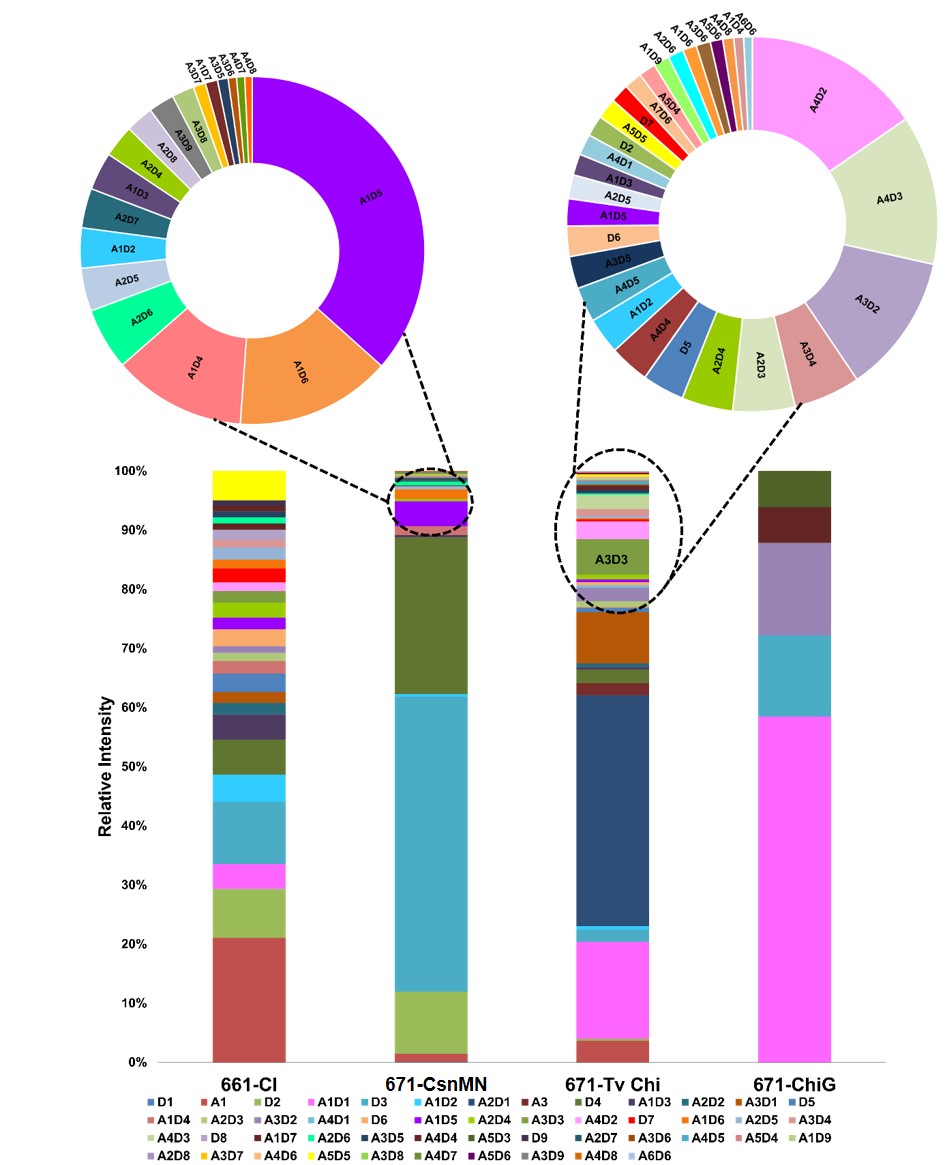

**Supplementary Figure S1****:** Relative intensity of the oligomeric products obtained by chemical (661-Cl produced by hydrochloric acid) and enzymatic (671-CsnMN produced by GH8 chitosanase, 671-TvChi by GH18 chitinase, 671-ChiG by GH19 chitinase) hydrolysis of chitosan polymer 651 (*F*_A_ 0.2) based on the UHPLC-ESI-MS analysis. A = GlcNAc, D = GlcN.

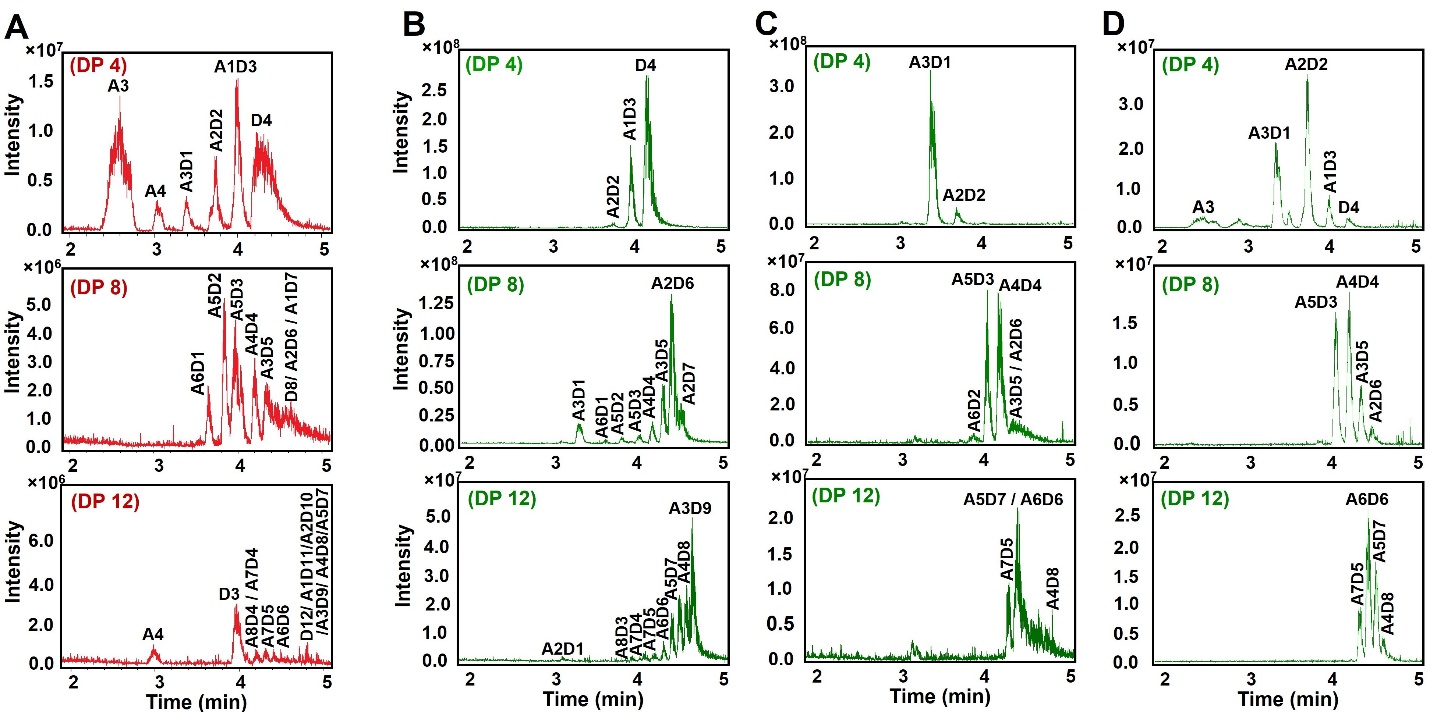

**Supplementary Figure S2:** HILIC-MS analysis of SEC purified tetra-, octa-, and dodecamer fractions of A) 661-Cl, B) 671-CsnMN, C) 671-TvChi, and D) 671-ChiG. The base peak chromatogram between minute 2 and minute 5 of the 10 min HILIC run are shown.

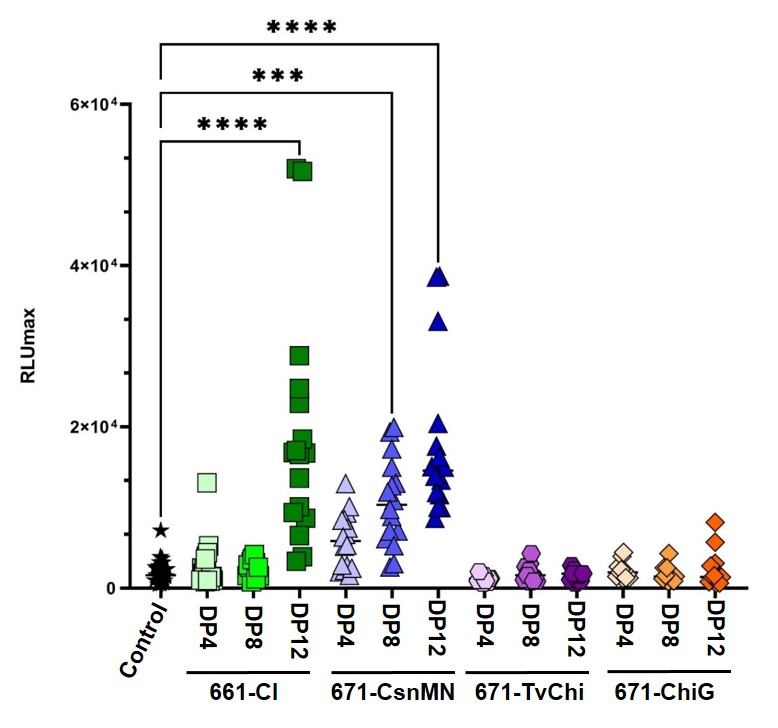

**Supplementary Figure S3:** Oxidative burst measurement of the SEC-purified oligomers at the concentration of 10 µg mL^‒1^, compared to control groups (ddH_2_O) in *N. benthamiana* seedlings (N = 3-5, n = 5-6).

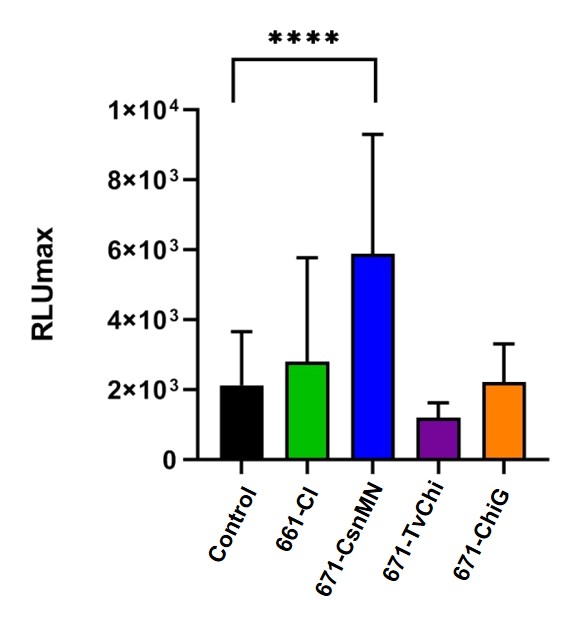

**Supplementary Figure S4:** Comparison of the eliciting activity of the SEC-purified DP 4 oligomers at the concentration of 10 µg mL^‒1^, compared to control groups (ddH_2_O) in seedlings of *N. benthamiana*. Data are presented as mean ± SD (N = 3, n = 6).

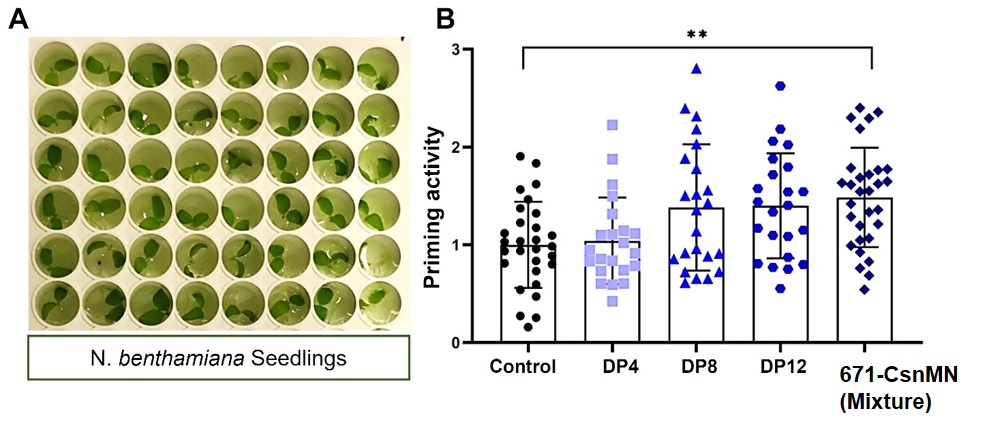

**Supplementary Figure S5:** Assessment of priming activity of chitosan 671-CsnMN and its SEC purified tetra, octa, and dodecamers in 6-day-old seedling of *N. benthamiana*. A) Digital image of representative seedlings after immersion in each well of the microtiter plates. B) The oxidative burst was measured after adding flg22 (0.025 ng mL^‒1^) to the seedlings treated with oligomers for 24 hours. The obtained values are normalized to the ddH_2_O-treated seedlings (control). Data are presented as mean ± SD (N=4-5, n=5-6).

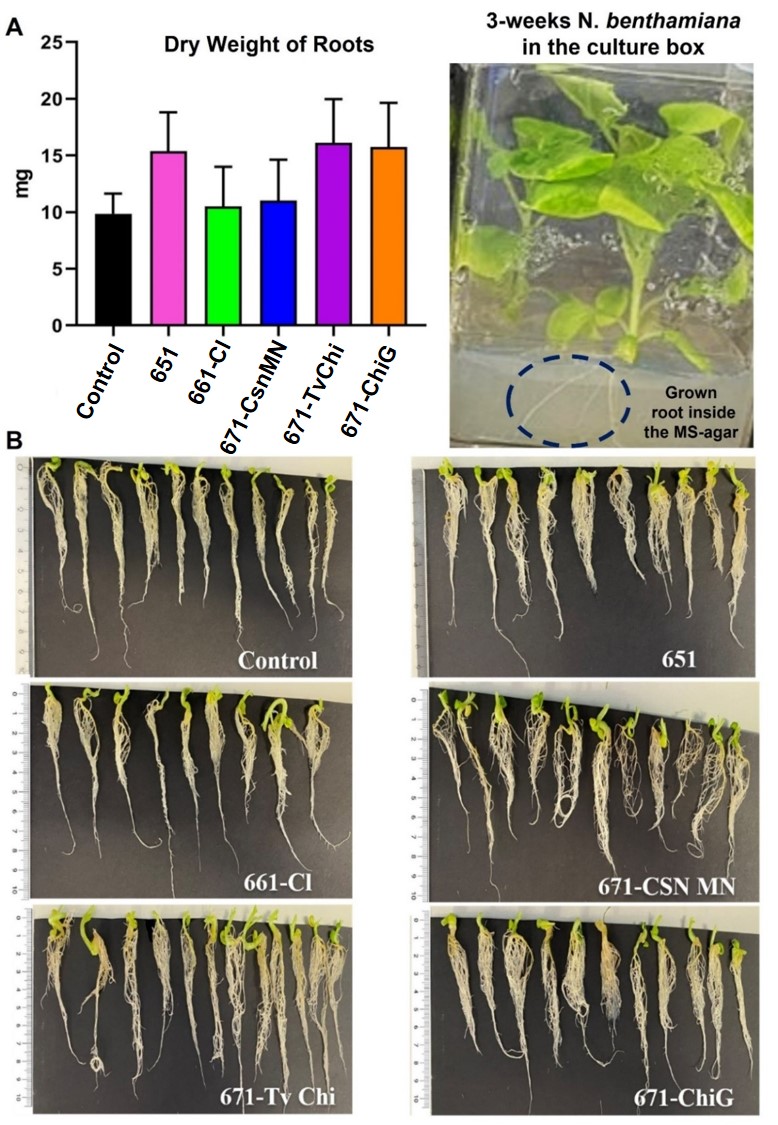

**Supplementary Figure S6:** Root growth analysis of the cultured *N.* *benthamiana* inside the vessel boxes. A) weight measurement of the dried roots after three weeks of growth inside the MS agar with and without chitosaccharides at the concentration of 100 µg mL^‒1^. No significant differences are observed in the root weight of the 671-CsnMN-treated plants compared to the control plants. This result further supports the biocompatibility of 671-CsnMN to the tested plants. B) Morphology and the length of the *N. benthamiana* roots.

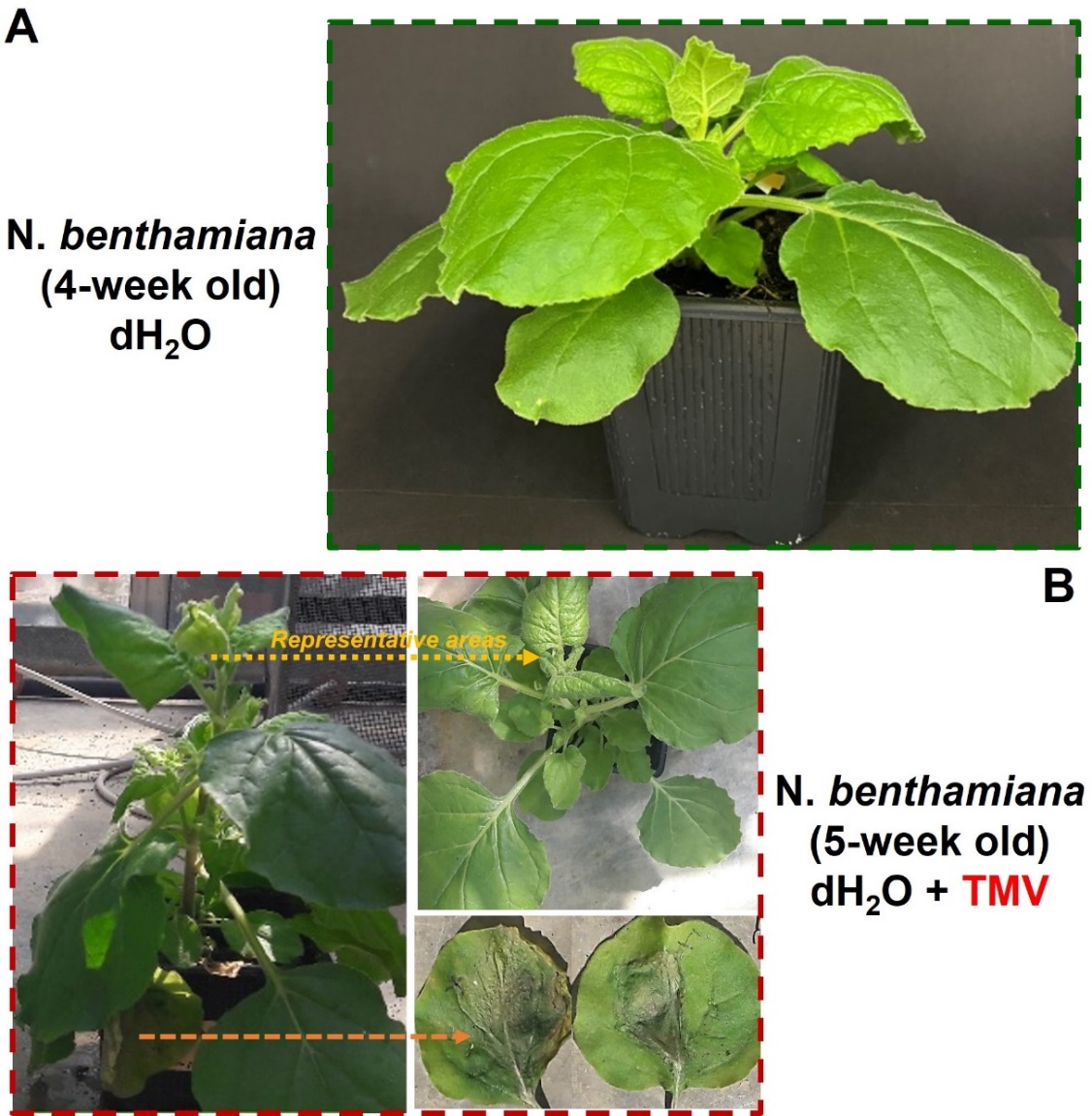

**Supplementary Figure S7:** Qualitative digital imaging of the *N. benthamiana* A) before and B) after inoculation with TMV for seven days. Our image depicted an infection sign in the local leaves of the TMV-inoculated plants (after syringe infiltration), and systemic chlorosis and curling symptoms are observed in these plans.

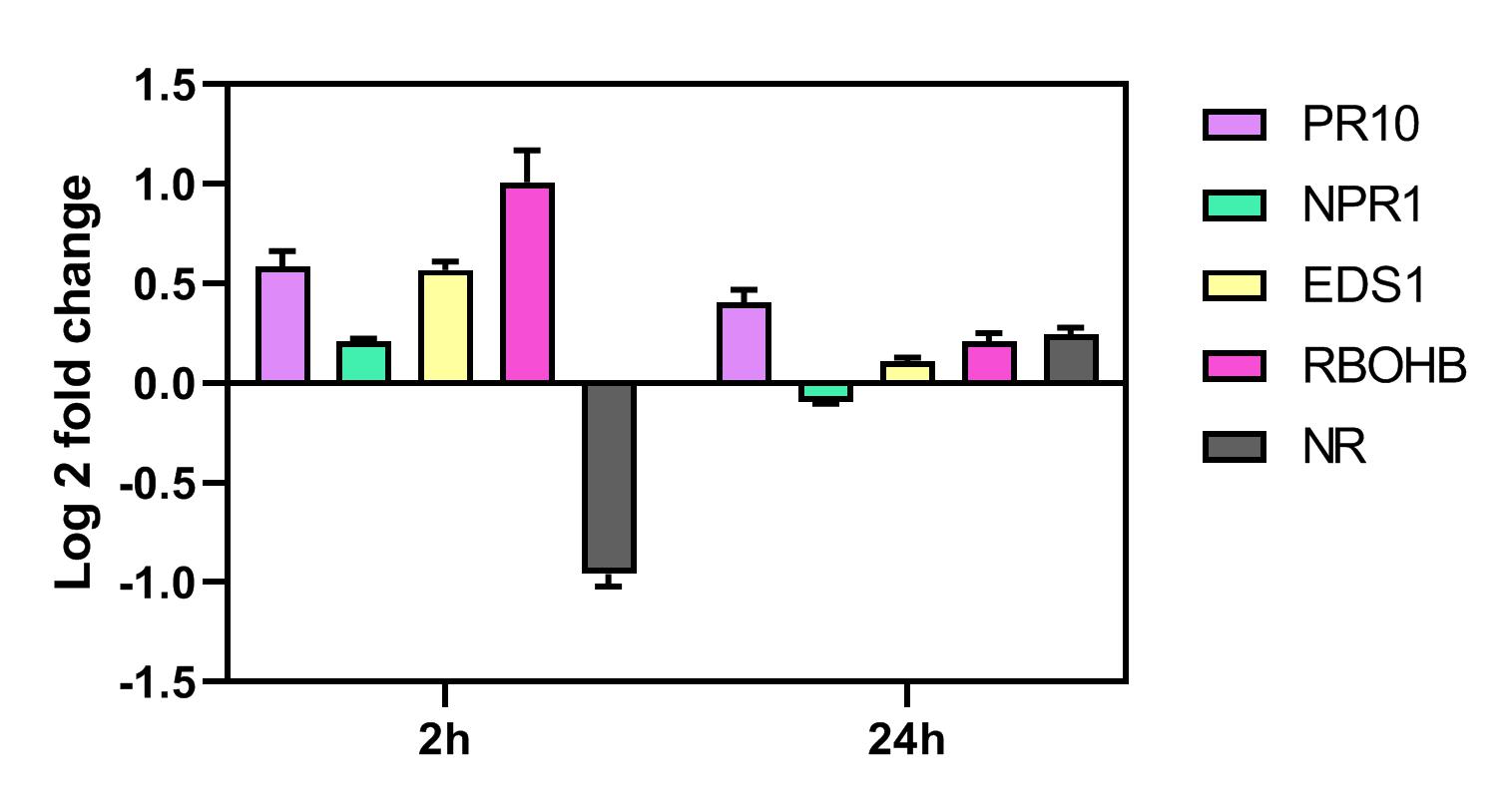

**Supplementary Figure S8:** Gene expression level of *N.* *benthamiana* after treatment with 671-CsnMN at the concentration of 100 µg mL^‒1^ after 2 and 24 hours*.* GAPDH is used as a reference gene in this assay*.* This data is presented as mean ± SD (N=1, n=3-9).

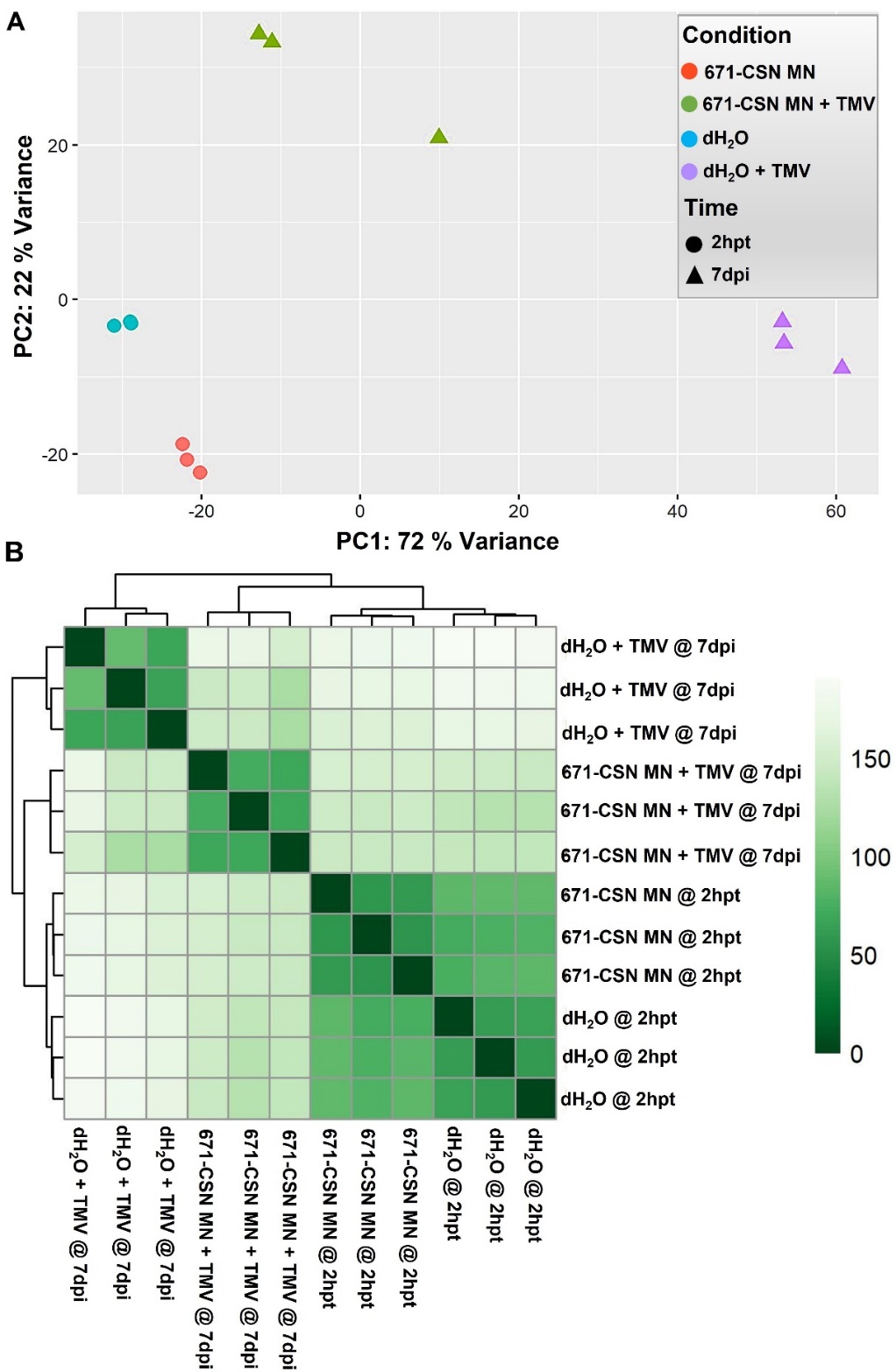

**Supplementary Figure S9:** A) Pearson’s correlation heatmap and B) Principle component analysis (PCA) of the identified genes of the treated *N.* *benthamiana* plants with chitosan 671-CsnMN at a concentration of 100 µg mL^‒1^ and pure ddH_2_O as the control group. Plants were foliar-sprayed with the chitooligomers in two different groups, without and with TMV inoculation, and harvested 2 hpt or 7 dpi, respectively. In the plotted Pairwise Pearson’s correlation, the coefficients presented in rows and columns of the graph are clustered based on their gene expression similarity. Furthermore, the calculated data in these samples are corroborated by performing a variance stabilizing transformation (VST) analysis using the VST package of the R program.

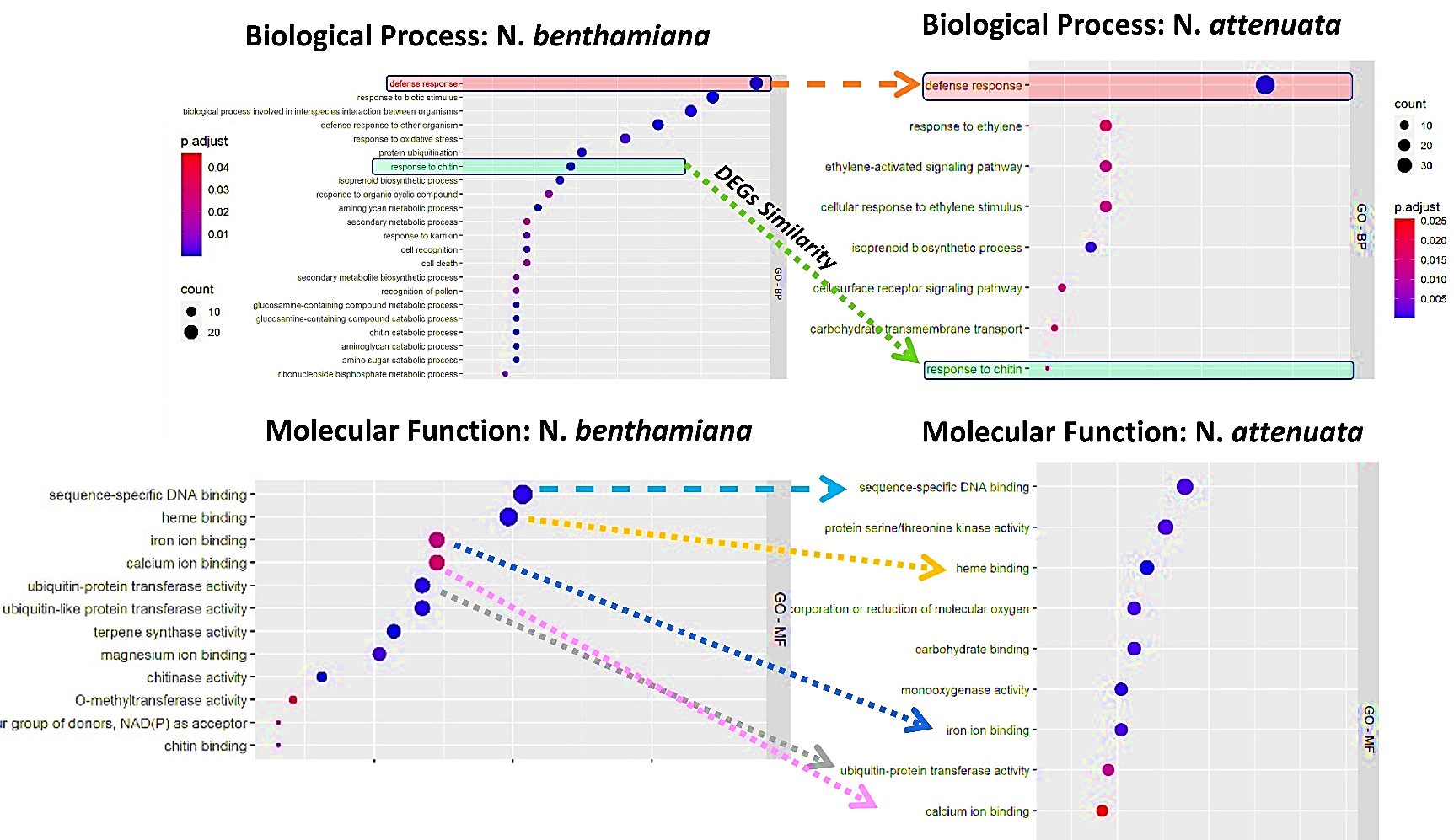

**Supplementary Figure S10:** GO over-representation analysis (ORA) similarities of the upregulated genes after treatment with 671-CsnMN and when the genes are comparably aligned to *N.* *benthamiana* and *N.* *attenuata* (FDR < 0.05 and log2 ≥ 1). The ORAs for each sub-ontology of BP: Biological Process, and MF: Molecular Function are computed using custom R scripts based on the enrichGO functions from the clusterProfiler package and ggplot2 for visualization. Terms which are significantly enriched relative to the background annotation of each reference genome are shown on the y-axis. The size and color of the dots on the x-axis indicate the number of genes and the degree of rich factor enrichment, respectively.

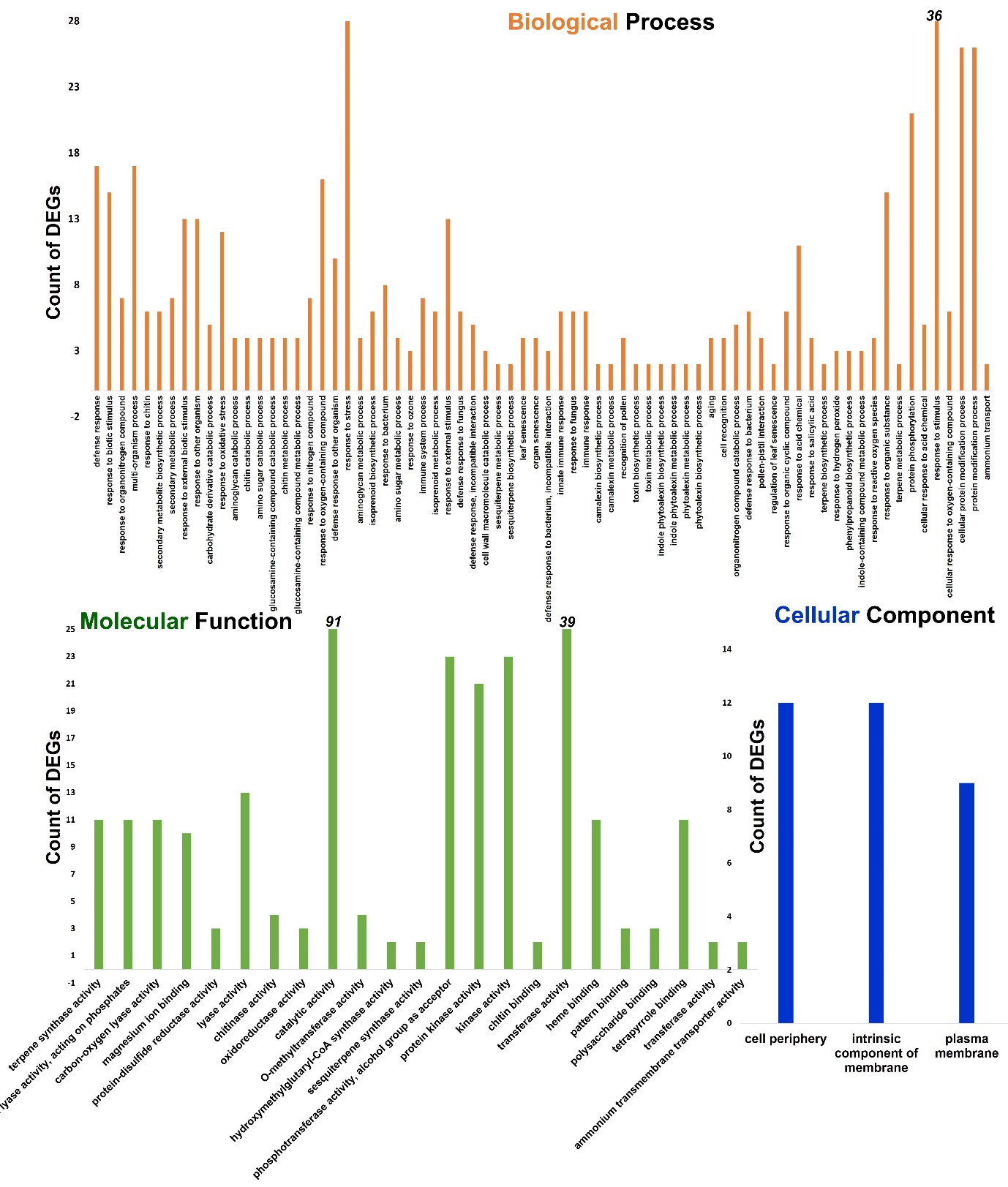

**Supplementary Figure S11:** GO analysis of the upregulated genes after treatment with 671-CsnMN (FDR < 0.05 and log2 ≥ 2). The DEGs of each sub-ontology (biological process, molecular function, and cellular component) are identified using Plantregmap data analysis. Terms which are significantly enriched relative to the background annotation of *N.* *benthamiana* genes are shown on the x-axis. The y-axis presents the numbers of DEGs in each GO of these samples (2hpt).

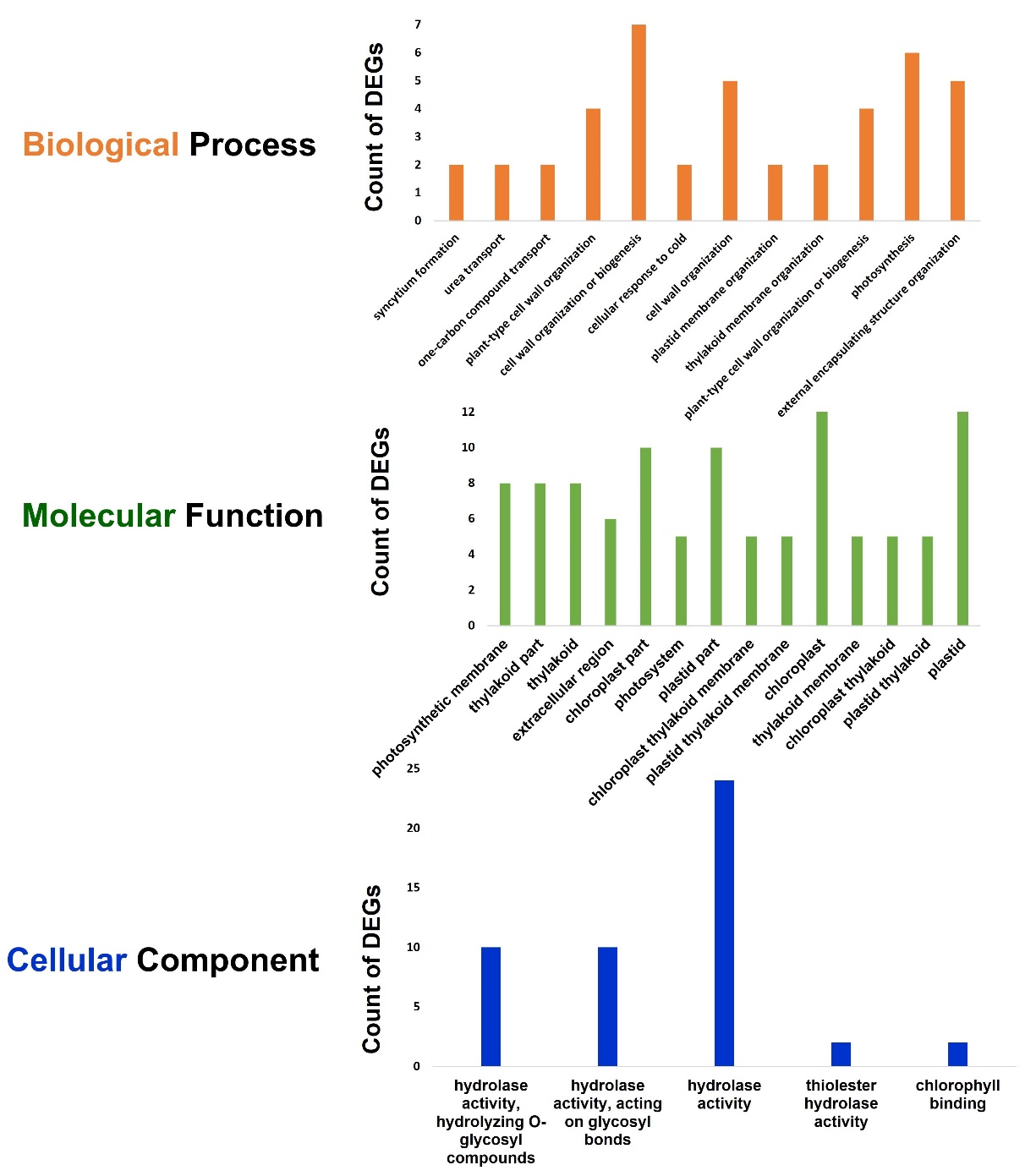

**Supplementary Figure S12:** GO analysis of the upregulated genes after treatment with 671-CsnMN and TMV inoculation (FDR < 0.05 and log2 ≥ 1.2). The DEGs of each sub-ontology (biological process, molecular function, and cellular component) are identified using Plantregmap data analysis. Terms which are significantly enriched relative to the background annotation of *N.* *benthamiana* genes are shown on the x-axis. The y-axis presents the numbers of DEGs in each GO of these samples (7 dpi).

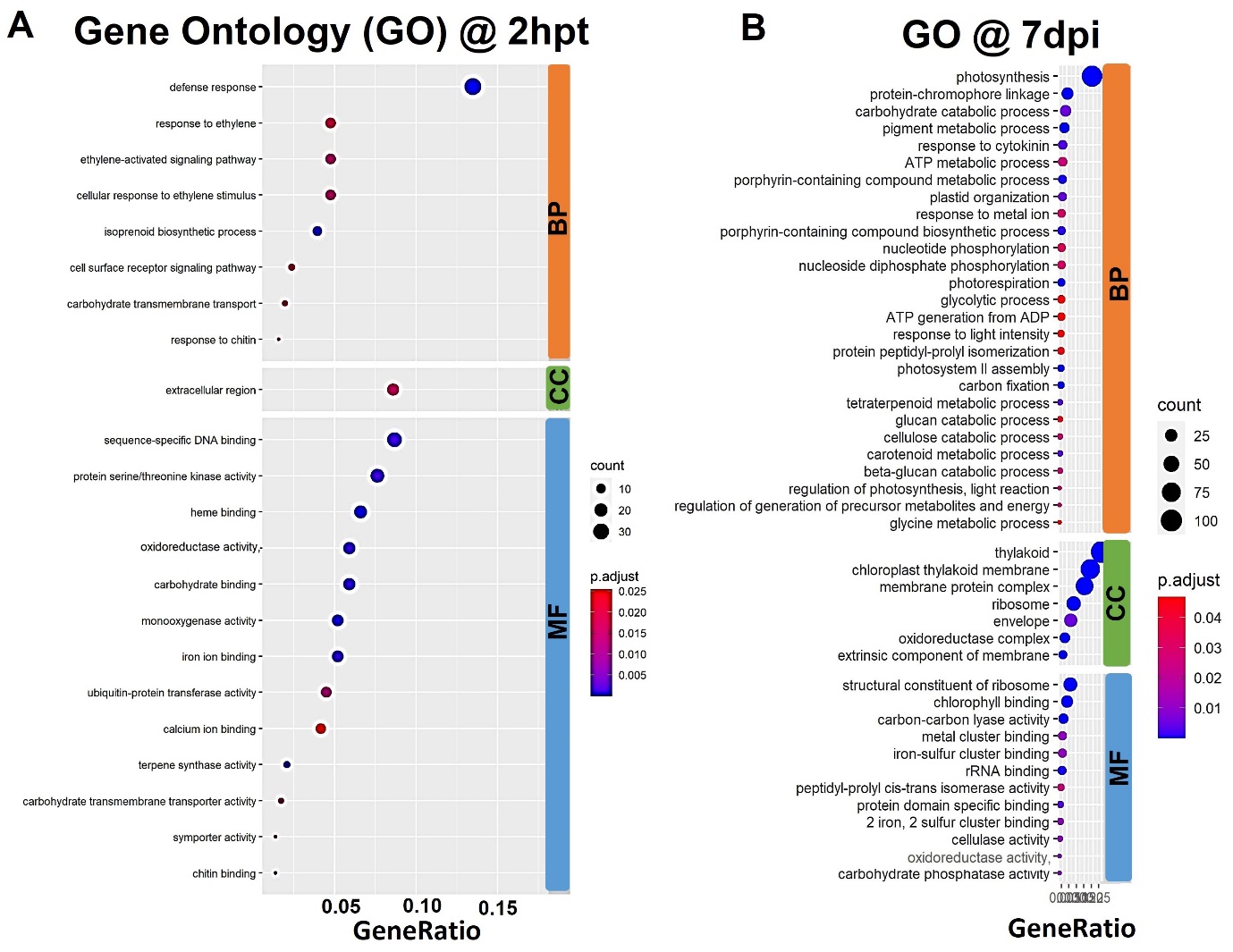

**Supplementary Figure S13:** GO analysis of the upregulated genes in the 671-CsnMN-treated plants A) without (2 hpt) and B) with TVM inoculation (7 dpi). The DEGs of each sub-ontology (biological process (BP), molecular function (MF), and cellular component (CC)) have been identified using R scripts based on the enrichGO functions from the clusterProfiler package. ggplot2 was used for data visualization of these samples. The statistically significant GO terms are aligned to the enriched relative background annotation of the *N.* *attenuata* genes and listed in the y-axis of this plot. The size and color of the dots on the x-axis indicate the number of genes and the degree of rich factor, respectively (FDR < 0.05 and log2 ≥ 1).

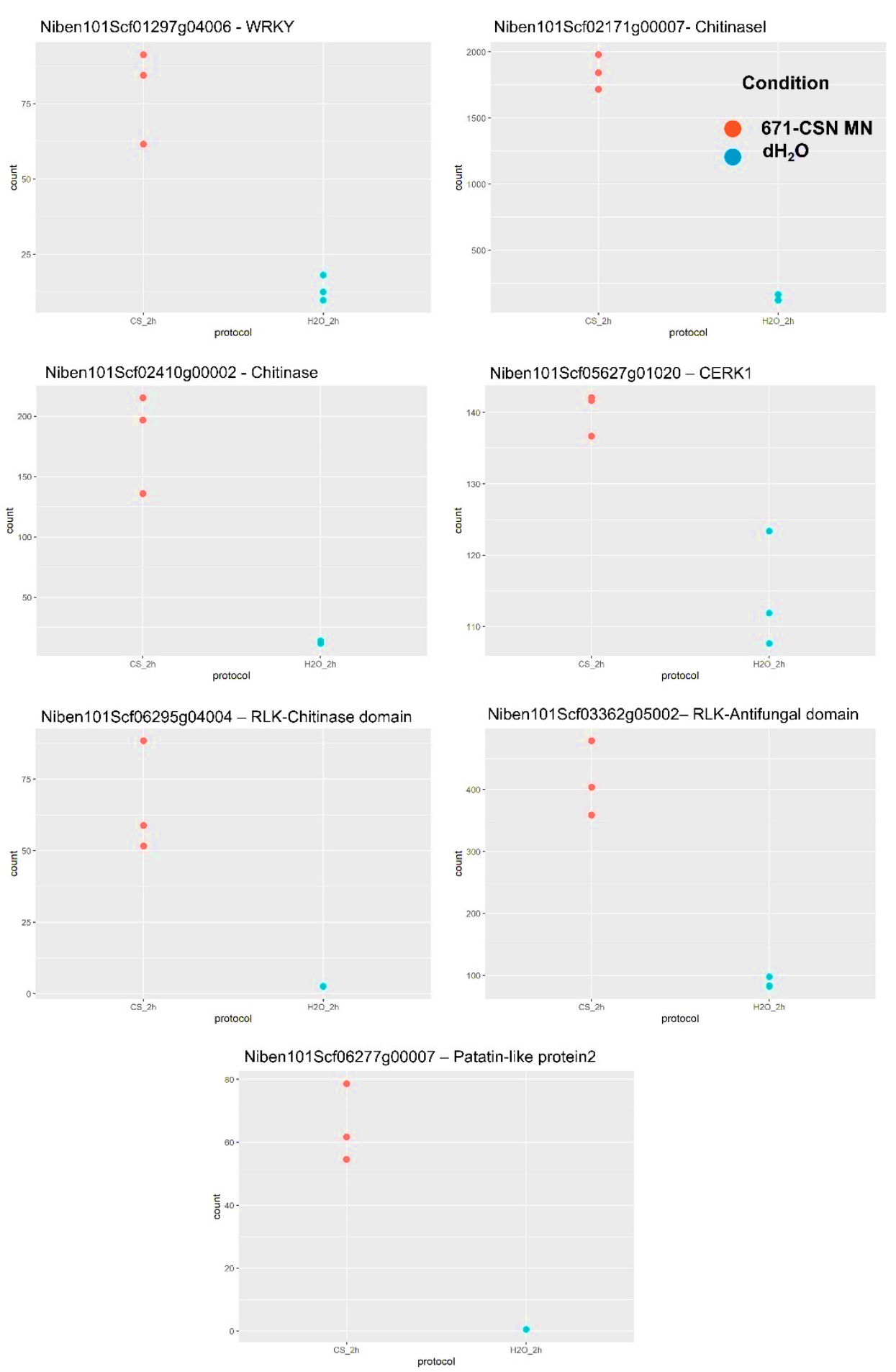

**Supplementary Figure S14:** Comparison of the read count of the RT-qPCR selected genes between the 671-CsnMN-treated plants and control groups (2 hpt).

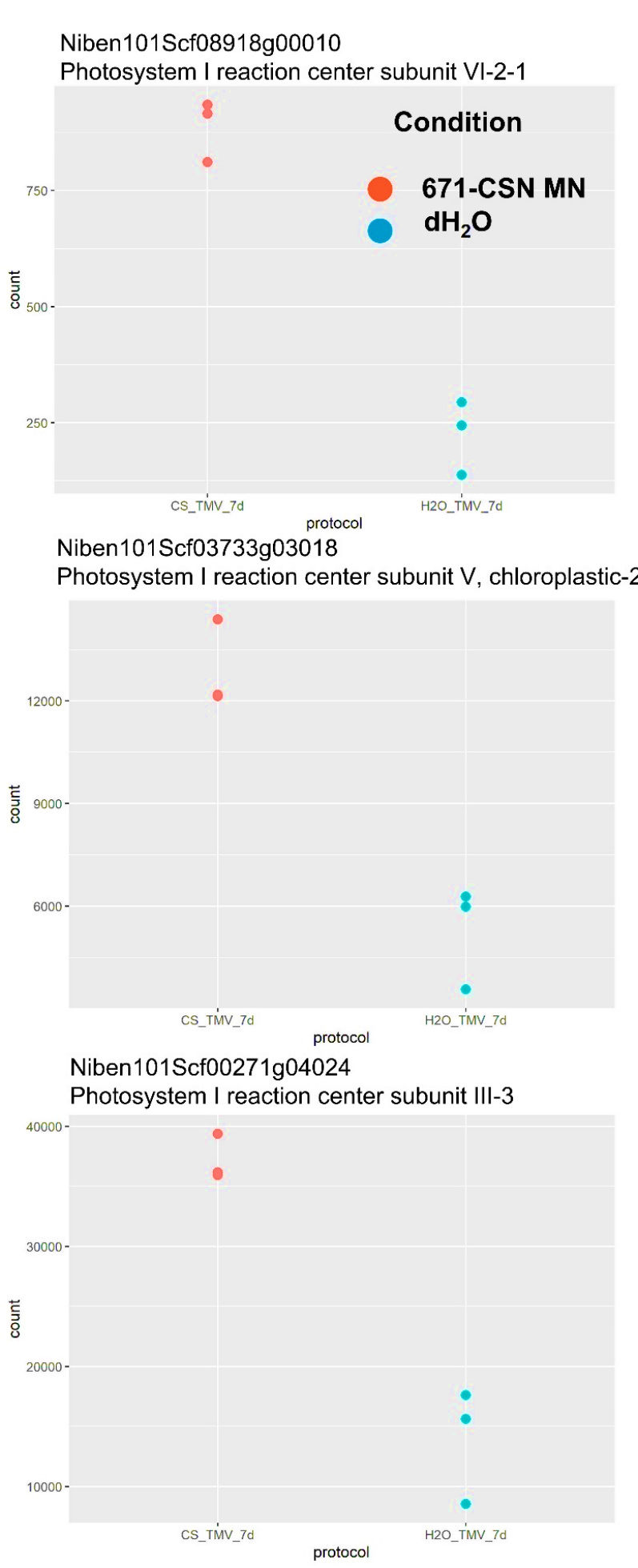

**Supplementary Figure S15:** Comparison of the read count of the RT-qPCR selected genes between the 671-CsnMN-treated plants and control groups after TMV inoculation (7 dpi).

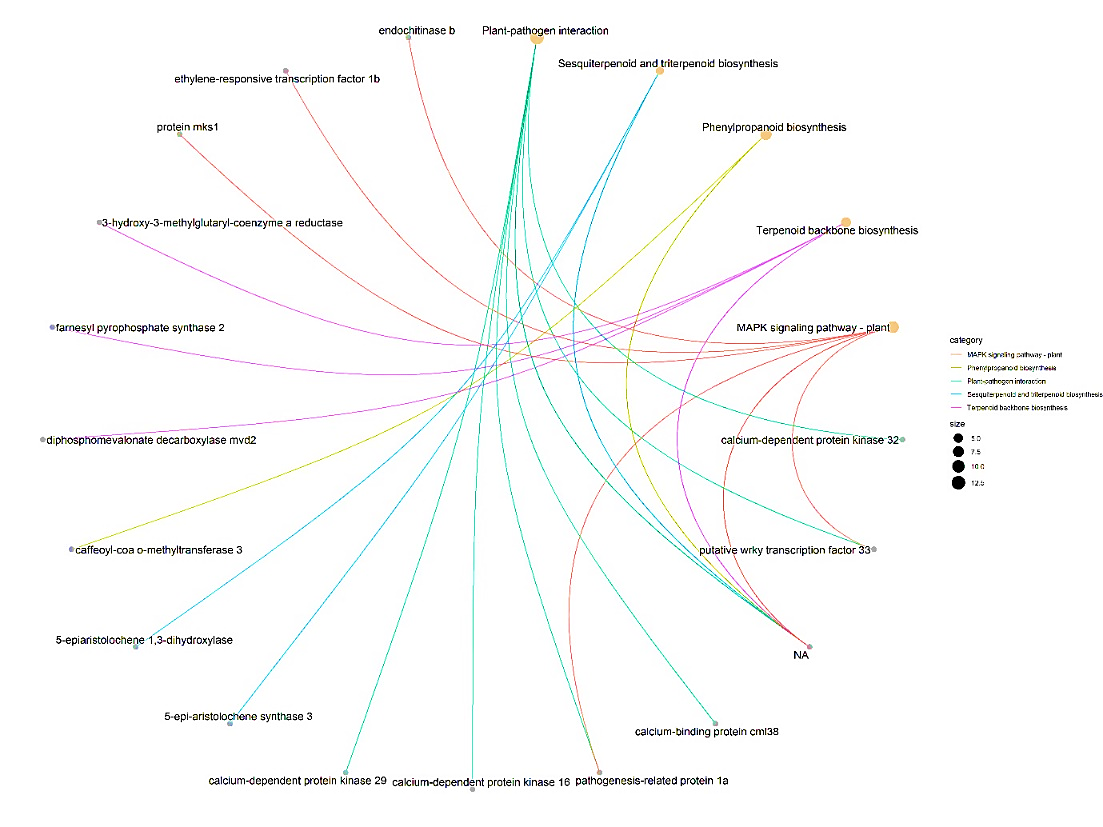

**Supplementary Figure S16:** Cnetplot of the enriched KEGG pathways in the 671-CsnMN-treated plants (2hpt). The enriched-KEGG pathways of these plants are compared to the control group. The size of the dots shows the number of DEGs which belong to the enriched KEGG. The lines depict the predicted interaction and their linkages. This data is derived from log2fold change based on the DESeq2 analysis and was generated using clusterProfiler R package (P < 0.05, FDR < 0.05, log2 ≥ 1).

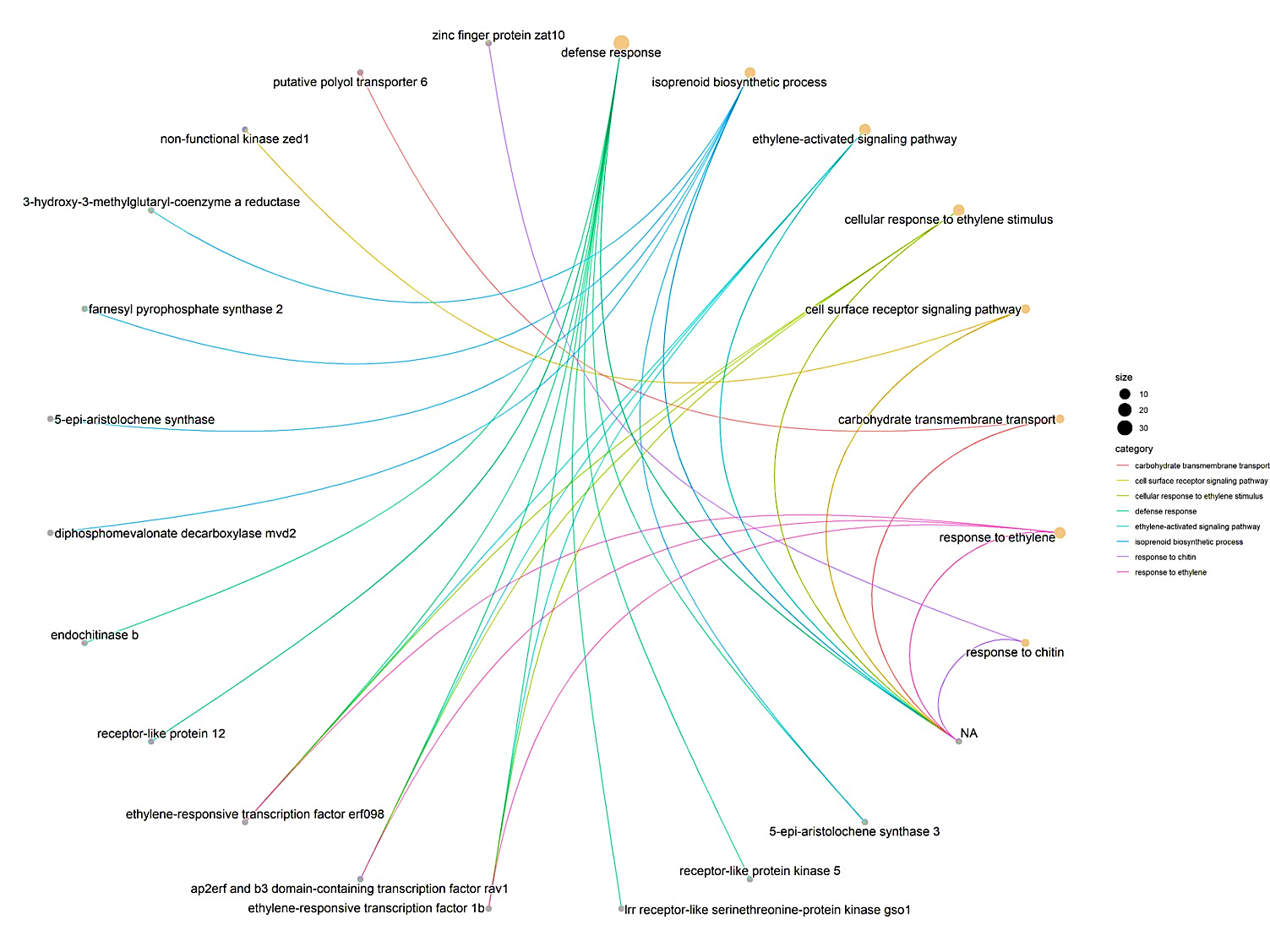

**Supplementary Figure S17:** Cnetplot of the enriched GO biological process (BP) in the 671-CsnMN-treated plants (2hpt). The gene ontology of these plants are compared to the control group. The size of the dots shows the number of DEGs which belong to the enriched BP. The lines depict the predicted interaction and their linkages. This data is derived from log2fold change based on the DESeq2 analysis and was generated using clusterProfiler R package (P < 0.05, FDR < 0.05, log2 ≥ 1).

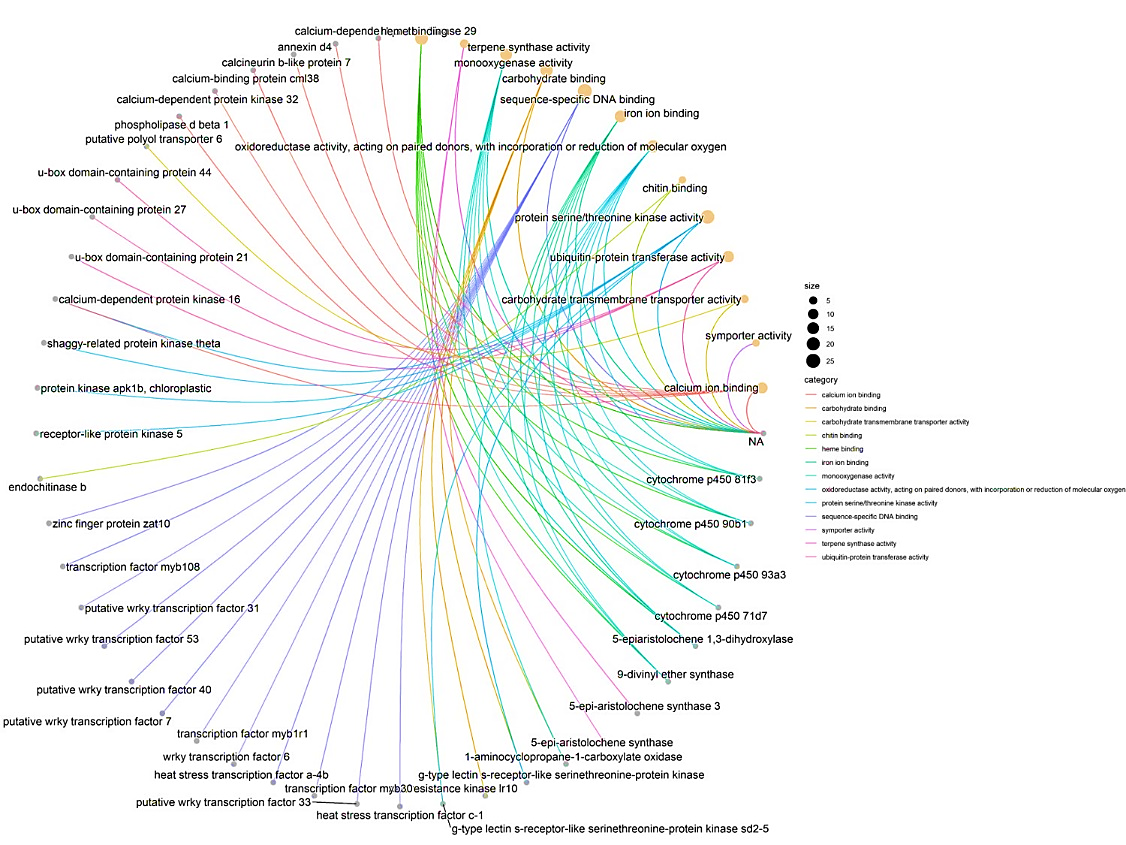

**Supplementary Figure S18:** Cnetplot of the enriched GO molecular function (MF) in the 671-CsnMN-treated plants (2hpt). The gene ontology of these plants are compared to the control group. The size of the dots shows the number of DEGs which belong to the enriched MF. The lines depict the predicted interaction and their linkages. This data is derived from log2fold change based on the DESeq2 analysis and was generated using clusterProfiler R package (P < 0.05, FDR < 0.05, log2 ≥ 1)**.**

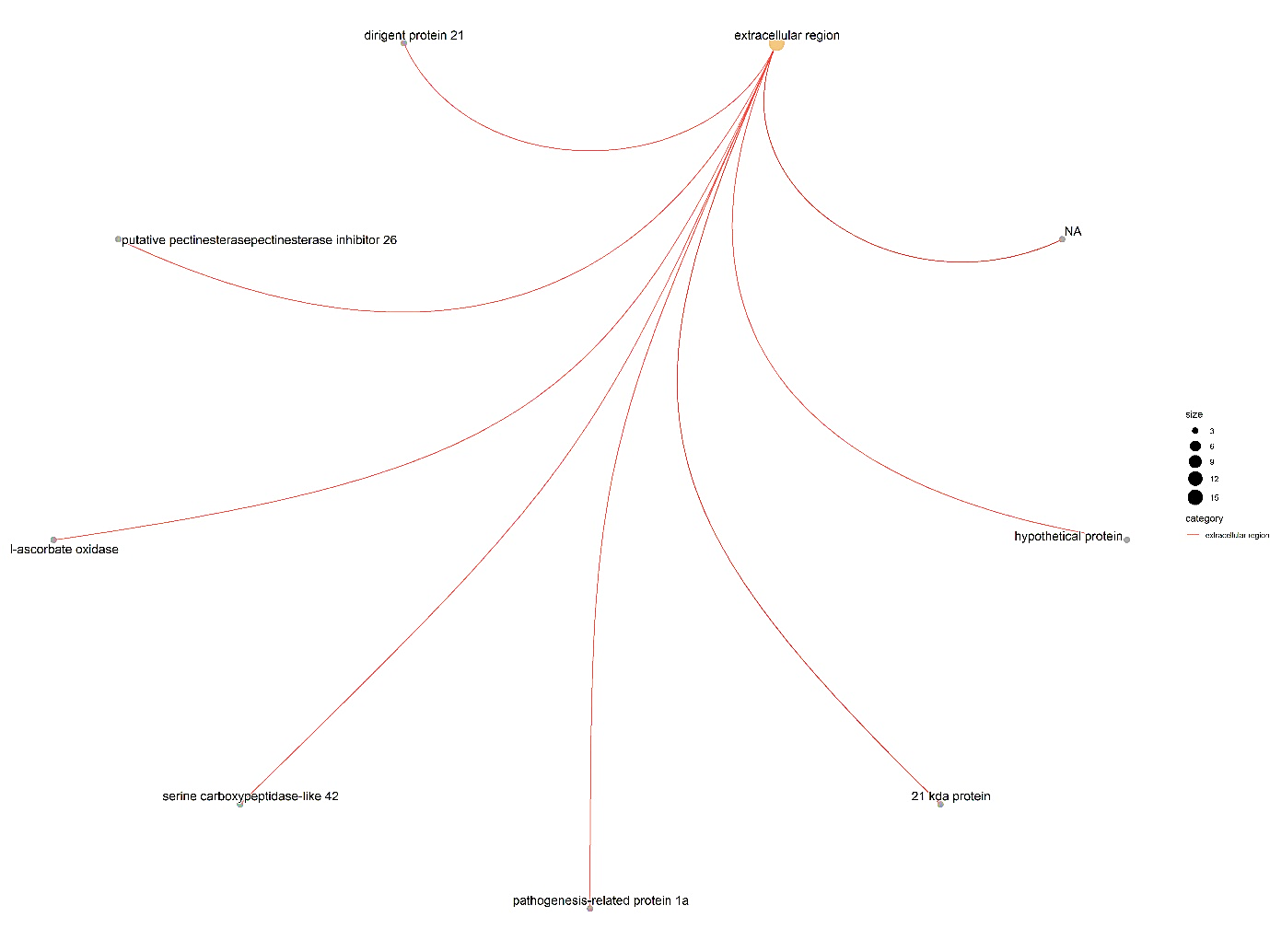

**Supplementary Figure S19:** Cnetplot of the enriched GO cellular component (CC) in the 671-CsnMN-treated plants (2hpt). The gene ontology of these plants are compared to the control group. The size of the dots shows the number of DEGs which belong to the enriched CC. The lines depict the predicted interaction and their linkages. This data is derived from log2fold change based on the DESeq2 analysis and was generated using clusterProfiler R package (P < 0.05, FDR < 0.05, log2 ≥ 1)**.**

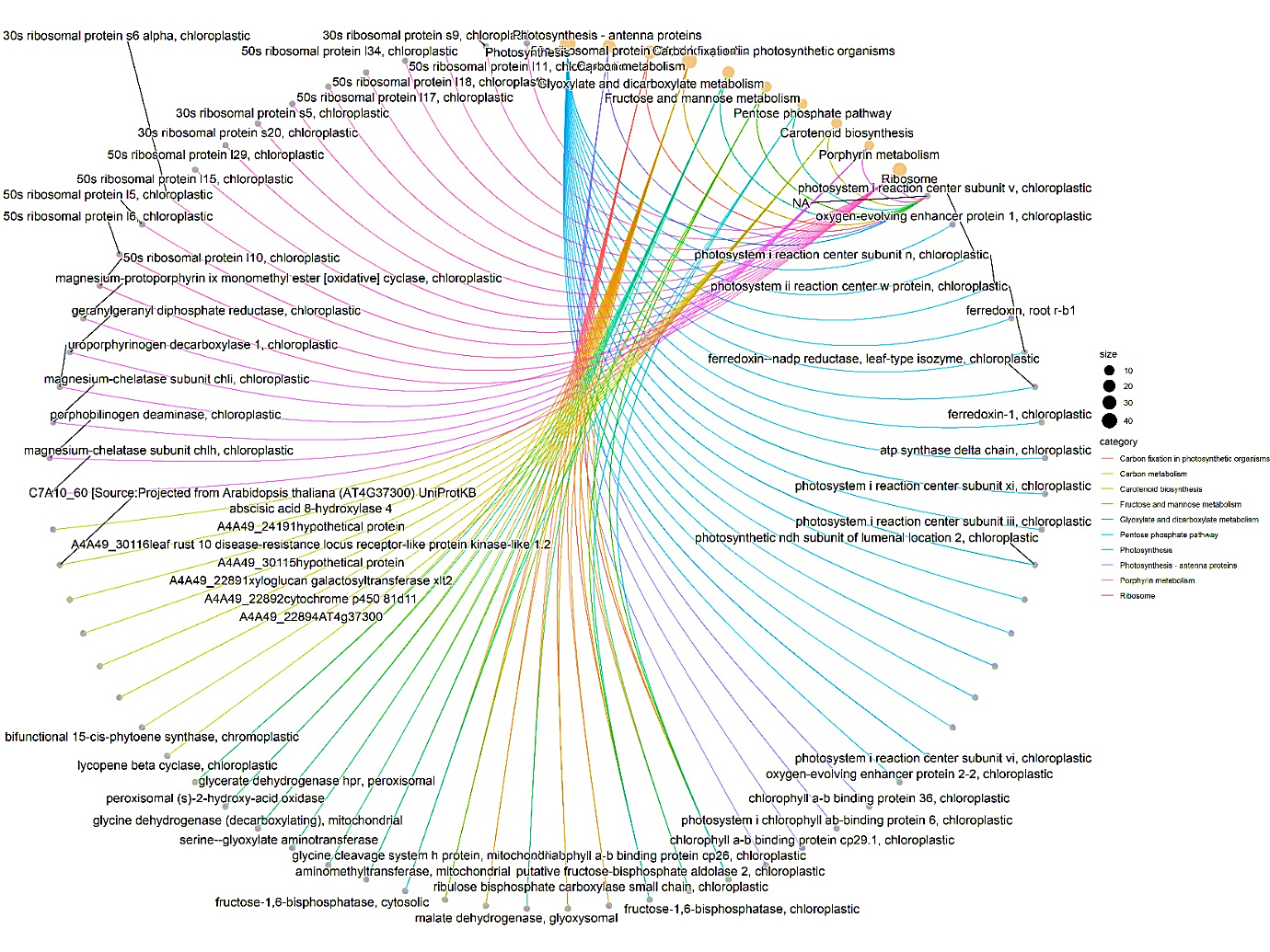

**Supplementary Figure S20:** Cnetplot of the enriched KEGG pathways in the 671-CsnMN-treated plants after TMV inoculation (7 dpi). The enriched-KEGG pathways of these plants are compared to the control group. The size of the dots shows the number of DEGs which belong to the enriched KEGG. The lines depict the predicted interaction and their linkages. This data is derived from log2fold change based on the DESeq2 analysis and was generated using clusterProfiler R package (P < 0.05, FDR < 0.05, log2 ≥ 1).

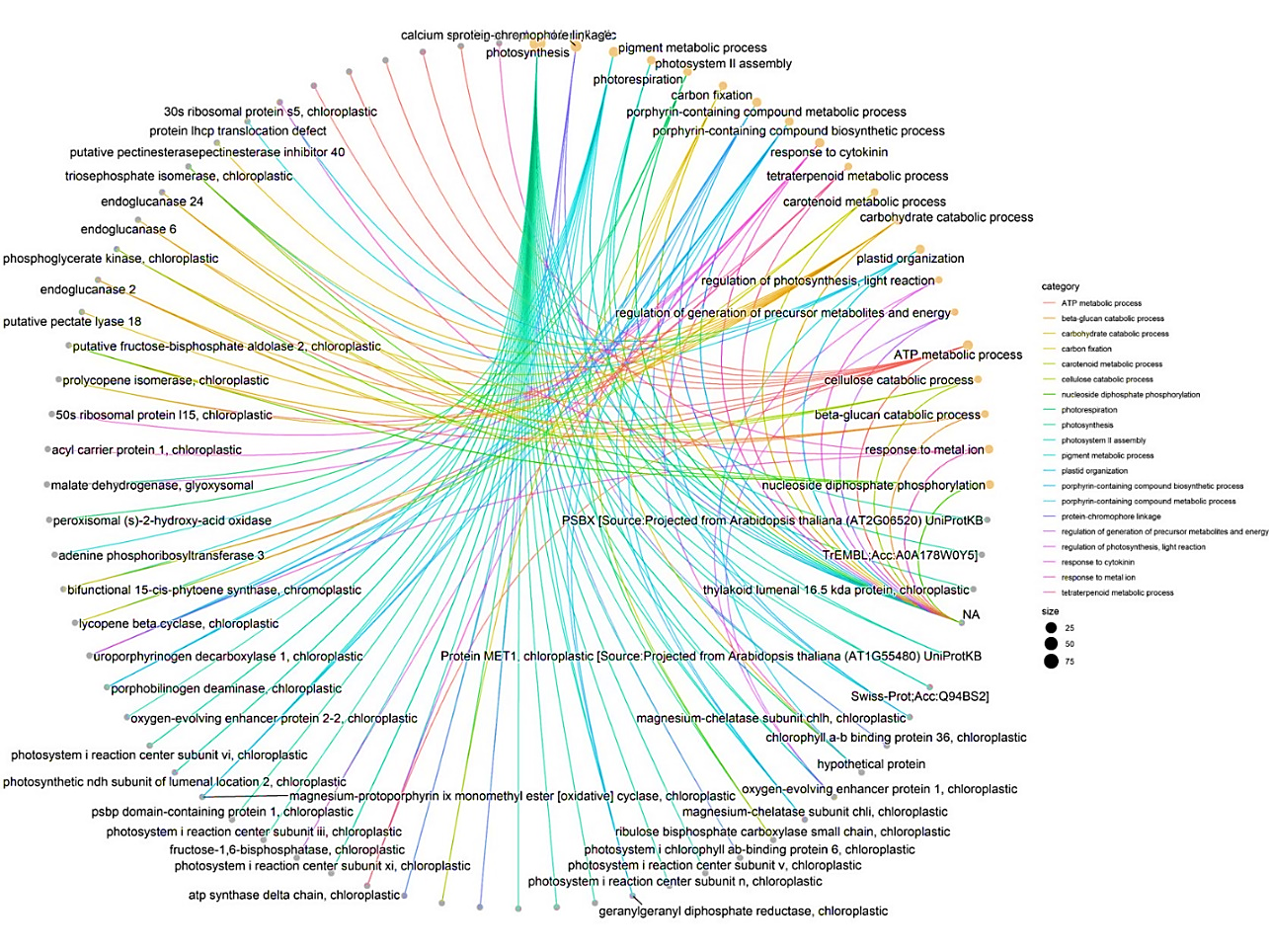

**Supplementary Figure S21:** Cnetplot of the enriched GO biological process (BP) in the 671-CsnMN-treated plants after TMV inoculation (7 dpi). The gene ontology of these plants are compared to the control group. The size of the dots shows the number of DEGs which belong to the enriched BP. The lines depict the predicted interaction and their linkages. This data is derived from log2fold change based on the DESeq2 analysis and was generated using clusterProfiler R package (P < 0.05, FDR < 0.05, log2 ≥ 1).

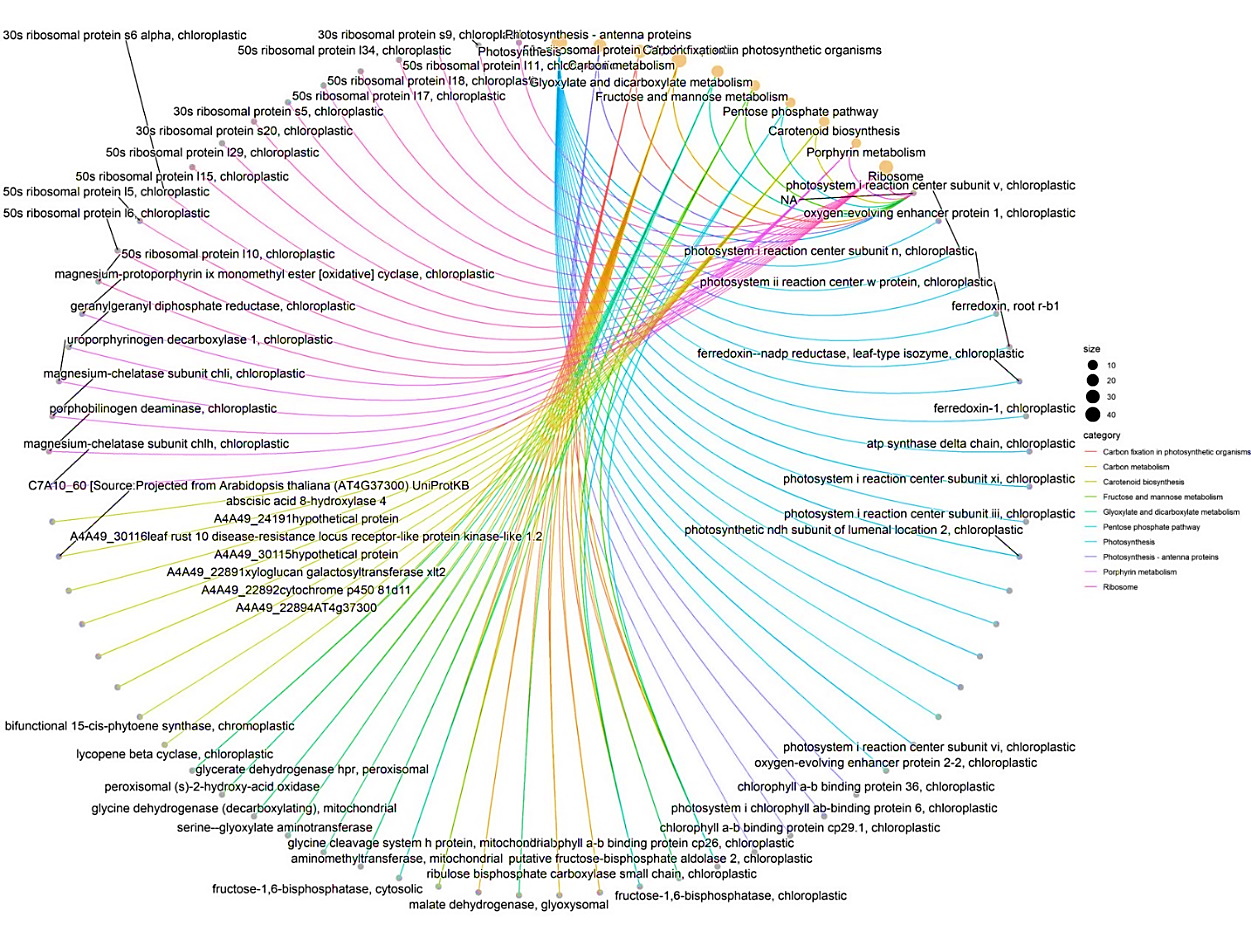

**Supplementary Figure S22:** Cnetplot of the enriched GO molecular function (MF) in the 671-CsnMN-treated plants after TMV inoculation (7 dpi). The gene ontology of these plants are compared to the control group. The size of the dots shows the number of DEGs which belong to the enriched MF. The lines depict the predicted interaction and their linkages. This data is derived from log2fold change based on the DESeq2 analysis and was generated using clusterProfiler R package (P < 0.05, FDR < 0.05, log2 ≥ 1)**.**

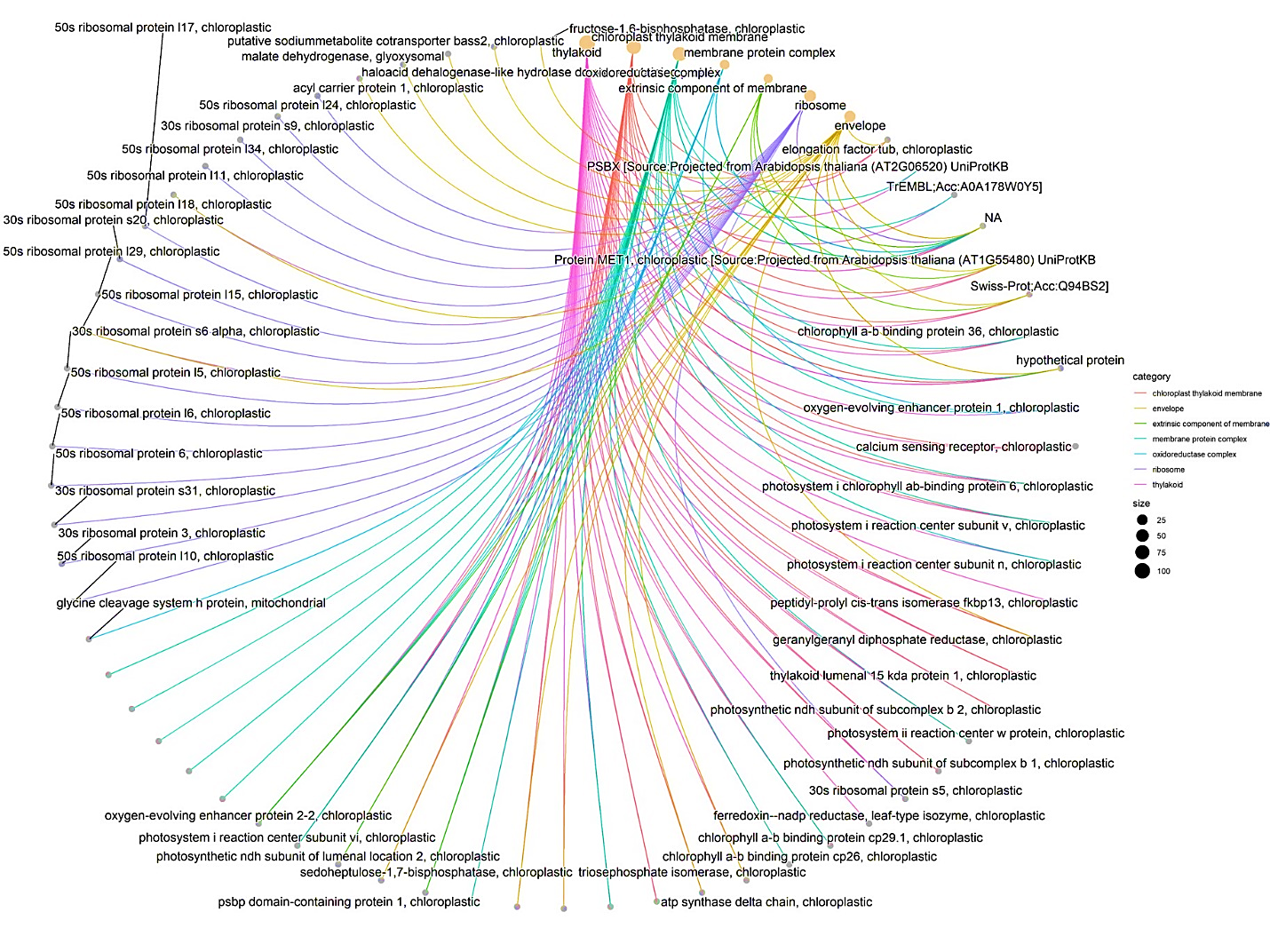

**Supplementary Figure S23:** Cnetplot of the enriched GO cellular component (CC) in the 671-CsnMN-treated plants after TMV inoculation (7 dpi). The gene ontology of these plants are compared to the control group. The size of the dots shows the number of DEGs which belong to the enriched CC. The lines depict the predicted interaction and their linkages. This data is derived from log2fold change based on the DESeq2 analysis and was generated using clusterProfiler R package (P < 0.05, FDR < 0.05, log2 ≥ 1)**.**

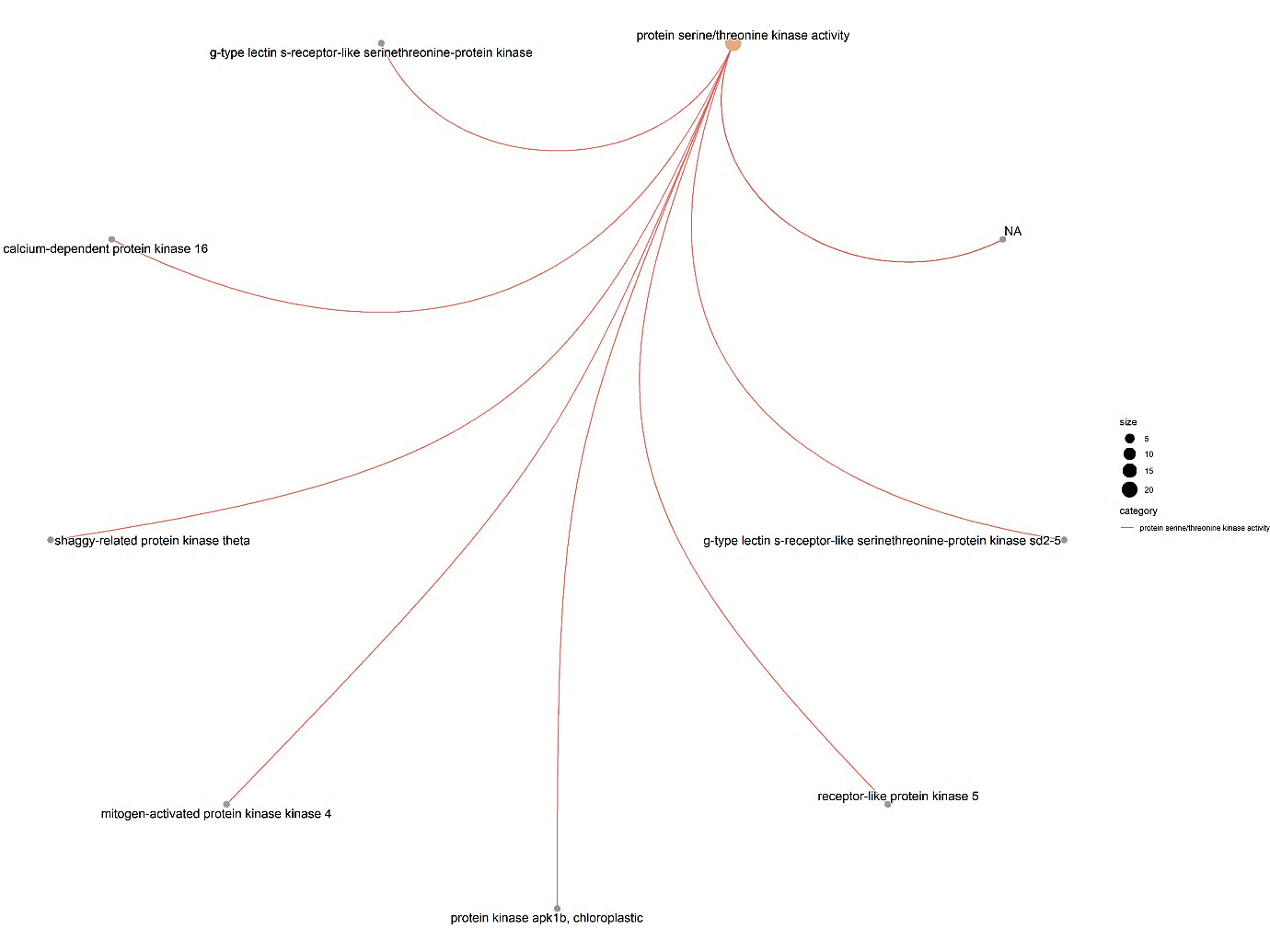

**Supplementary Figure S24:** Cnetplot of the enriched protein serine/threonine kinase activity genes upregulated in the 671-CsnMN-treated plants (2hpt). This plot was derived from log2fold change based on the DESeq2 analysis and was generated using the clusterProfiler R package (P < 0.05, FDR < 0.05, log2 ≥ 1).

**Table S1:** The used primers for RT-qPCR measurement in the current study.

| **Gene** | **Forward Primer** | **Reverse Primer** | **Amplicon(bp)** |
| --- | --- | --- | --- |
| GAPDH^1^ | AGCTCAAGGTTAAGGATGAC | TGGCCAAGGGAGCAAGGCAA | 350 |
| NPR1 | GCCCAATAAGCTCCCTTTTAGGA | TGGTACAGGTGTTGTATGTATGCT | 131 |
| RBOHB | AACAACCTCGGATACATTAT | TGTAAATAGACCAGCCATAA | 246 |
| NR | CACCTCCAACGCTAAAGAA | TGCCAGCATTGATGAGAAT | 195 |
| EDS1 | ATGGTGAGAATTGAAGAGG | ACTGTACCAACCATCAACAGA | 180 |
| PR10 | CGACAAACTTGAGTTCGTAACC | GCTTGATCTTTGCCTTCACT | 180 |
| NRG1 | CTGCTCTTGGTCCAGTTTTTG | AATTTGTAAAGATAAAGCA | 187 |
| WRKY | GGTTTCAGTGATCAAAGAATG | GAAGAGTATTGGAATTGTCAAAC | 196 |
| Chitinase 1 | GAACAATCTTGGCACAAACC | GCAGCTTGATCTGCTATTCC | 199 |
| Chitinase 2 | ATGAAATCATCCATCCCAAG | CAGGACAATTAGCATCGTTG | 268 |
| CERK1 | CCTGAGGATAACTTGACGG | GATATCCCAGCTATTGCTCC | 191 |
| Patatin-like protein | GCAAGAAAATGTTGGTTCTGTC | CAATATCCATAGATGCAGCCTC | 264 |
| RLK-chitinase domain | ATGGCATTCAAACACATTTTC | GAAATCAATAGCAATTCTCCAATC | 525 |
| RLK-antifungal domain | ATGGATTCTCCCGTCAAATG | GACAAGTCTTTCATGCATTCACC | 266 |
| Photosystem I subunit VI-2-1 | ACAAATCCAACTCTCCCAAAG | CTGGTGCTGCTGACACTG | 120 |
| Photosystem I V, chloroplastic2 | CTTGAGCACAGGACTCTCAC | CAACTATGTTGAAACCAACAGG | 185 |
| Photosystem I subunit III-3 | CTCACCACACTCCATCTCC | GAAGACAAAGCAAGTGCAG | 267 |

^1^ Reference gene

**Table S2:** The upregulated genes in the 671-CsnMN-treated *N.* *bethamiana*, 2 hours after foliar spraying (Log2 ≥ 2).

| **Gene ID** | **baseMean** | **log2FoldChange** | **lfcSE** | **pvalue** | **padj** |
| --- | --- | --- | --- | --- | --- |
| Niben101Scf06277g00007.1 | 17.5759799 | 6.6369596 | 1.10713077 | 1.0938E-11 | 2.4532E-09 |
| Niben101Scf00821g16001.1 | 97.1748044 | 6.62822868 | 1.11701431 | 1.5524E-11 | 3.3983E-09 |
| Niben101Scf06245g01017.1 | 76.5224857 | 6.51851489 | 0.84665688 | 1.6431E-16 | 7.9038E-14 |
| Niben101Scf05124g01018.1 | 30.4026323 | 5.82746122 | 0.68402552 | 4.8689E-19 | 2.7384E-16 |
| Niben101Scf01683g03005.1 | 12.0297092 | 5.61031932 | 1.26598127 | 4.6059E-08 | 4.6006E-06 |
| Niben101Scf08390g03006.1 | 21.4422002 | 5.61008339 | 1.13721814 | 8.4324E-09 | 1.0366E-06 |
| Niben101Scf04418g07001.1 | 7.62748353 | 5.50461752 | 1.22890232 | 4.6465E-08 | 4.6285E-06 |
| Niben101Scf03861g00008.1 | 7.15600535 | 5.32196164 | 1.24303109 | 1.0095E-07 | 9.4147E-06 |
| Niben101Ctg06260g00001.1 | 42.3704855 | 5.29883671 | 0.55743068 | 9.6969E-23 | 7.8778E-20 |
| Niben101Scf04555g01001.1 | 23.0835097 | 5.07299631 | 0.65530144 | 4.3321E-16 | 1.9081E-13 |
| Niben101Scf02405g05001.1 | 8.02819878 | 5.03889664 | 1.28918379 | 3.3214E-07 | 2.7785E-05 |
| Niben101Scf04933g02001.1 | 15.9262456 | 4.93814609 | 0.72828573 | 4.6478E-13 | 1.2776E-10 |
| Niben101Scf02511g04008.1 | 17.8881507 | 4.91641592 | 0.9163676 | 2.0636E-09 | 2.9289E-07 |
| Niben101Scf01587g00006.1 | 7.49594253 | 4.89525568 | 1.34116724 | 6.3623E-07 | 4.907E-05 |
| Niben101Scf10067g02017.1 | 56.9835202 | 4.8585836 | 0.52535526 | 1.2411E-21 | 9.0745E-19 |
| Niben101Scf11922g01019.1 | 10.3715067 | 4.74741 | 1.00186148 | 3.8094E-08 | 3.912E-06 |
| Niben101Scf04362g07011.1 | 61.260211 | 4.67046802 | 0.85289257 | 1.2768E-09 | 1.8975E-07 |
| Niben101Ctg15035g00004.1 | 140.22577 | 4.67042884 | 0.93206076 | 1.2542E-08 | 1.4791E-06 |
| Niben101Scf11512g00011.1 | 4.75840365 | 4.66562773 | 1.57845846 | 2.2024E-06 | 0.00014655 |
| Niben101Scf00927g08010.1 | 12.5996361 | 4.63452747 | 1.43432953 | 1.6434E-06 | 0.00011335 |
| Niben101Scf06295g04004.1 | 42.0115992 | 4.6154046 | 0.67164691 | 2.617E-13 | 7.4163E-11 |
| Niben101Scf00551g06025.1 | 13.7796956 | 4.60871792 | 0.79230262 | 1.9903E-10 | 3.4982E-08 |
| Niben101Scf09334g02006.1 | 55.5872976 | 4.57220291 | 0.39260921 | 1.4686E-32 | 2.4405E-29 |
| Niben101Scf00212g03016.1 | 10.1795442 | 4.45924789 | 0.89681961 | 1.6088E-08 | 1.8437E-06 |
| Niben101Scf11922g01018.1 | 37.8211994 | 4.43305624 | 0.41550662 | 7.9527E-28 | 1.0383E-24 |
| Niben101Scf08510g01007.1 | 39.4622035 | 4.38722189 | 0.43460135 | 3.1179E-25 | 3.2567E-22 |
| Niben101Scf07725g00004.1 | 239.562786 | 4.3608401 | 0.9455531 | 7.4272E-08 | 7.2407E-06 |
| Niben101Scf00072g05003.1 | 301.859826 | 4.33364217 | 0.4388389 | 2.7643E-24 | 2.6594E-21 |
| Niben101Scf11183g00001.1 | 79.5001091 | 4.30017033 | 0.31353583 | 4.7884E-44 | 1.945E-40 |
| Niben101Scf12789g00006.1 | 365.359832 | 4.27920709 | 0.92912619 | 7.8414E-08 | 7.5439E-06 |
| Niben101Scf01084g01009.1 | 70.364963 | 4.25278099 | 0.38154074 | 3.9431E-30 | 6.0064E-27 |
| Niben101Scf03706g00007.1 | 96.2171409 | 4.24761715 | 0.61150766 | 1.5108E-13 | 4.4542E-11 |
| Niben101Ctg09679g00001.1 | 88.4447504 | 4.20900107 | 0.33519362 | 1.9605E-37 | 5.1194E-34 |
| Niben101Scf02398g00005.1 | 21.0415184 | 4.12848253 | 0.53126964 | 3.2804E-16 | 1.4805E-13 |
| Niben101Ctg07683g00002.1 | 13.6680022 | 4.12601768 | 0.78766401 | 4.7992E-09 | 6.38E-07 |
| Niben101Scf03993g06006.1 | 585.41017 | 4.0938794 | 1.02657309 | 5.9656E-07 | 4.6206E-05 |
| Niben101Ctg15860g00004.1 | 9.1538791 | 4.05989335 | 0.87501606 | 7.353E-08 | 7.1874E-06 |
| Niben101Scf02069g00017.1 | 14.8603383 | 4.02097409 | 0.67154519 | 7.4388E-11 | 1.4164E-08 |
| Niben101Scf00072g05004.1 | 203.653013 | 3.99099012 | 0.8221933 | 3.046E-08 | 3.2183E-06 |
| Niben101Scf03292g03005.1 | 13.2453205 | 3.91795465 | 0.73602164 | 3.165E-09 | 4.3336E-07 |
| Niben101Scf21242g00003.1 | 24.1009766 | 3.88203858 | 0.56402739 | 2.191E-13 | 6.3071E-11 |
| Niben101Scf07372g00014.1 | 13.7399089 | 3.84770034 | 1.79772559 | 1.3334E-05 | 0.00070547 |
| Niben101Scf01729g01015.1 | 383.228329 | 3.77101875 | 0.19830934 | 5.4669E-82 | 9.993E-78 |
| Niben101Scf08921g02018.1 | 286.90408 | 3.76132352 | 1.72846849 | 1.4468E-05 | 0.00075562 |
| Niben101Scf02171g00007.1 | 396.707518 | 3.75333887 | 0.16958506 | 6.651E-110 | 2.432E-105 |
| Niben101Scf03164g03009.1 | 3.69594316 | 3.71639226 | 1.95303736 | 1.8001E-05 | 0.00090644 |
| Niben101Scf02936g04013.1 | 15.3722638 | 3.69264876 | 0.88524817 | 4.2176E-07 | 3.434E-05 |
| Niben101Scf03993g05005.1 | 669.863587 | 3.67326759 | 1.28206716 | 6.826E-06 | 0.00039736 |
| Niben101Scf06538g02009.1 | 8.49755381 | 3.65010792 | 0.98353626 | 1.4468E-06 | 0.00010172 |
| Niben101Scf01111g01003.1 | 218.425549 | 3.6146965 | 0.29505126 | 6.4407E-36 | 1.3081E-32 |
| Niben101Scf07355g02001.1 | 26.7776279 | 3.61431055 | 0.60907793 | 1.0362E-10 | 1.923E-08 |
| Niben101Scf05619g00015.1 | 142.166916 | 3.60845355 | 0.37317245 | 1.5439E-23 | 1.3766E-20 |
| Niben101Scf02410g00002.1 | 184.04399 | 3.59899234 | 0.61513609 | 1.7087E-10 | 3.062E-08 |
| Niben101Scf00372g05013.1 | 23.1208739 | 3.58832703 | 0.66408762 | 2.159E-09 | 3.0475E-07 |
| Niben101Scf02819g00006.1 | 72.3214646 | 3.58807688 | 0.32707053 | 2.0306E-29 | 2.7495E-26 |
| Niben101Scf00870g00001.1 | 11.2698552 | 3.57592636 | 0.86869969 | 5.177E-07 | 4.0877E-05 |
| Niben101Scf10986g00001.1 | 43.7019921 | 3.5690253 | 0.37065571 | 2.2799E-23 | 1.9383E-20 |
| Niben101Scf02351g08017.1 | 7.51118047 | 3.53776877 | 0.94973187 | 1.4721E-06 | 0.0001031 |
| Niben101Scf03830g02010.1 | 14.2360673 | 3.49200036 | 0.7486866 | 7.6816E-08 | 7.4687E-06 |
| Niben101Scf10126g00007.1 | 19.0643704 | 3.48146605 | 0.83461313 | 4.5254E-07 | 3.644E-05 |
| Niben101Scf03202g12016.1 | 40.2962244 | 3.48097985 | 0.51186014 | 3.8537E-13 | 1.0837E-10 |
| Niben101Scf00158g04018.1 | 31.0118778 | 3.46262644 | 0.48048708 | 2.1405E-14 | 7.1139E-12 |
| Niben101Scf10688g01013.1 | 139.386673 | 3.43532008 | 0.29630434 | 1.6632E-32 | 2.6437E-29 |
| Niben101Scf00960g15004.1 | 108.05393 | 3.40478284 | 0.43429924 | 1.706E-16 | 8.0996E-14 |
| Niben101Scf00679g00001.1 | 12.3420239 | 3.39928083 | 0.59823155 | 4.8008E-10 | 7.7316E-08 |
| Niben101Scf03422g04036.1 | 38.6443773 | 3.3981753 | 0.3915244 | 1.504E-19 | 9.0134E-17 |
| Niben101Scf04143g01004.1 | 20.2457737 | 3.39790717 | 0.5801178 | 1.7258E-10 | 3.0776E-08 |
| Niben101Scf07725g01004.1 | 119.118688 | 3.39702855 | 1.15372117 | 7.2182E-06 | 0.00041953 |
| Niben101Scf03985g04006.1 | 16.5589394 | 3.35455308 | 0.61813356 | 2.0308E-09 | 2.9001E-07 |
| Niben101Scf11175g00005.1 | 26.1406772 | 3.34982313 | 0.57058265 | 1.6195E-10 | 2.926E-08 |
| Niben101Scf02819g00005.1 | 147.494911 | 3.328015 | 0.25636021 | 5.8797E-40 | 1.9541E-36 |
| Niben101Scf03810g07006.1 | 30.7861009 | 3.28227003 | 0.47248338 | 1.4744E-13 | 4.3821E-11 |
| Niben101Scf01521g00004.1 | 53.1546385 | 3.28047365 | 0.35628146 | 1.3003E-21 | 9.321E-19 |
| Niben101Scf02407g03010.1 | 179.494489 | 3.26475058 | 0.27461482 | 5.2532E-34 | 9.6023E-31 |
| Niben101Scf03532g01002.1 | 5.51659354 | 3.26057609 | 1.6500784 | 3.0001E-05 | 0.00142068 |
| Niben101Scf01795g09001.1 | 19.8760394 | 3.24127851 | 0.52794401 | 3.297E-11 | 6.8484E-09 |
| Niben101Scf04015g01004.1 | 134.526536 | 3.24116364 | 0.42270505 | 7.0303E-16 | 2.9206E-13 |
| Niben101Scf03602g01007.1 | 435.816904 | 3.21334506 | 0.86926506 | 1.7891E-06 | 0.00012249 |
| Niben101Scf02547g00007.1 | 25.6586209 | 3.18397366 | 0.4877746 | 2.7767E-12 | 6.6347E-10 |
| Niben101Scf03600g02023.1 | 47.6952408 | 3.16538402 | 0.49990364 | 1.013E-11 | 2.2861E-09 |
| Niben101Scf00149g10020.1 | 54.5215237 | 3.12000404 | 0.30875971 | 2.219E-25 | 2.3859E-22 |
| Niben101Scf06650g03009.1 | 122.29923 | 3.10212785 | 0.68058138 | 1.3335E-07 | 1.2097E-05 |
| Niben101Scf04296g00015.1 | 40.433548 | 3.09577352 | 0.35515312 | 1.2488E-19 | 7.6091E-17 |
| Niben101Scf08273g02004.1 | 22.8774281 | 3.09203615 | 0.48270116 | 6.6019E-12 | 1.5275E-09 |
| Niben101Scf04594g02003.1 | 34.859765 | 3.0546036 | 0.60173751 | 1.4159E-08 | 1.6485E-06 |
| Niben101Scf00789g00009.1 | 190.333514 | 3.02984933 | 0.25816887 | 3.7491E-33 | 6.5266E-30 |
| Niben101Scf00163g22004.1 | 41.3221108 | 3.01628177 | 0.77721564 | 1.2077E-06 | 8.6569E-05 |
| Niben101Scf02736g02001.1 | 52.5949665 | 3.01580607 | 0.46048966 | 2.7224E-12 | 6.5477E-10 |
| Niben101Scf06275g04013.1 | 190.066701 | 3.0095535 | 0.57747484 | 7.4375E-09 | 9.3116E-07 |
| Niben101Scf03503g00003.1 | 33.5168789 | 2.99723263 | 0.51070419 | 2.0147E-10 | 3.5073E-08 |
| Niben101Scf05787g01001.1 | 41.974718 | 2.99215798 | 1.13100863 | 1.4706E-05 | 0.00076259 |
| Niben101Scf07477g02005.1 | 19.5921898 | 2.97778839 | 0.52105444 | 5.013E-10 | 8.0029E-08 |
| Niben101Scf03385g02011.1 | 1276.46757 | 2.97709102 | 0.24303594 | 7.9681E-36 | 1.5331E-32 |
| Niben101Scf05675g01018.1 | 163.511993 | 2.97450884 | 0.58630409 | 1.4984E-08 | 1.728E-06 |
| Niben101Ctg14168g00004.1 | 14.3145314 | 2.9709535 | 0.66866035 | 2.1468E-07 | 1.8866E-05 |
| Niben101Scf03191g04001.1 | 115.409173 | 2.96948178 | 0.39545041 | 2.9421E-15 | 1.1088E-12 |
| Niben101Scf10436g03001.1 | 35.8429574 | 2.94947736 | 0.43978689 | 9.9897E-13 | 2.5718E-10 |
| Niben101Scf07063g00001.1 | 106.159893 | 2.94801058 | 0.305255 | 2.266E-23 | 1.9383E-20 |
| Niben101Scf00557g03004.1 | 54.576833 | 2.94401295 | 0.39631006 | 5.5695E-15 | 1.9962E-12 |
| Niben101Scf04240g03001.1 | 675.676589 | 2.91177285 | 0.19173138 | 2.172E-53 | 1.3234E-49 |
| Niben101Scf20576g01017.1 | 15.3788198 | 2.90951039 | 0.56930552 | 1.2974E-08 | 1.5153E-06 |
| Niben101Scf00700g00005.1 | 40.9915823 | 2.90848713 | 0.4078652 | 5.2419E-14 | 1.624E-11 |
| Niben101Scf03791g01002.1 | 16.2874477 | 2.90581967 | 0.79443067 | 2.1772E-06 | 0.00014524 |
| Niben101Ctg14008g00002.1 | 198.162973 | 2.90384549 | 0.31041831 | 4.3888E-22 | 3.2744E-19 |
| Niben101Scf00372g05012.1 | 20.7917059 | 2.89632559 | 0.91485179 | 6.1838E-06 | 0.00036639 |
| Niben101Scf02370g02008.1 | 261.557498 | 2.89169151 | 0.5533722 | 7.4373E-09 | 9.3116E-07 |
| Niben101Scf01109g03004.1 | 111.346053 | 2.88881215 | 0.47071496 | 4.3405E-11 | 8.7669E-09 |
| Niben101Scf02513g00012.1 | 34.0152619 | 2.87942597 | 0.48177329 | 1.1665E-10 | 2.143E-08 |
| Niben101Scf00573g00010.1 | 52.9827836 | 2.86766275 | 0.67277788 | 4.1213E-07 | 3.3706E-05 |
| Niben101Scf01237g00013.1 | 7.96353624 | 2.86670894 | 1.18290074 | 2.2351E-05 | 0.00109824 |
| Niben101Scf07829g01008.1 | 33.7808761 | 2.84281077 | 0.68086812 | 5.472E-07 | 4.2745E-05 |
| Niben101Scf05151g03018.1 | 15.2271274 | 2.8237003 | 0.62673936 | 1.7969E-07 | 1.5945E-05 |
| Niben101Scf02877g02005.1 | 388.929332 | 2.82333677 | 1.08637542 | 1.7314E-05 | 0.00087546 |
| Niben101Scf03861g00009.1 | 17.6769467 | 2.81914041 | 0.65664819 | 3.777E-07 | 3.1169E-05 |
| Niben101Scf02601g04010.1 | 7.20296112 | 2.81761855 | 1.3116317 | 3.4734E-05 | 0.00159722 |
| Niben101Scf00994g00001.1 | 94.4324856 | 2.79129315 | 0.6676094 | 5.4475E-07 | 4.2645E-05 |
| Niben101Scf05368g06017.1 | 8.13482797 | 2.78489748 | 1.73682799 | 6.9064E-05 | 0.00286778 |
| Niben101Scf04869g03002.1 | 669.200258 | 2.76407478 | 0.82534701 | 4.4895E-06 | 0.00027677 |
| Niben101Scf00149g10001.1 | 18.6219552 | 2.76273773 | 0.63670699 | 3.2897E-07 | 2.7647E-05 |
| Niben101Scf12515g00025.1 | 49.9520216 | 2.75750101 | 0.43261109 | 1.106E-11 | 2.4654E-09 |
| Niben101Scf01001g07015.1 | 16.0753377 | 2.75290405 | 1.24252859 | 3.2788E-05 | 0.00152364 |
| Niben101Scf05938g00001.1 | 15.0357477 | 2.74104575 | 0.73991258 | 2.0351E-06 | 0.00013701 |
| Niben101Scf00520g06006.1 | 23.7080859 | 2.72862335 | 0.51495653 | 5.6945E-09 | 7.4075E-07 |
| Niben101Scf00109g02027.1 | 90.6210221 | 2.72104263 | 0.96526219 | 1.2488E-05 | 0.00066972 |
| Niben101Scf04286g01010.1 | 17.6341247 | 2.71690361 | 0.53238137 | 1.4939E-08 | 1.728E-06 |
| Niben101Scf01374g15011.1 | 530.982434 | 2.70665589 | 0.46952386 | 4.7044E-10 | 7.6099E-08 |
| Niben101Ctg16217g00002.1 | 29.6201002 | 2.70505504 | 0.44666176 | 8.4962E-11 | 1.601E-08 |
| Niben101Scf07608g00007.1 | 48.4338362 | 2.69823529 | 0.40379253 | 1.5483E-12 | 3.7988E-10 |
| Niben101Scf02203g05002.1 | 345.288308 | 2.68564727 | 0.28572922 | 3.8669E-22 | 2.9451E-19 |
| Niben101Scf08799g00001.1 | 120.984482 | 2.671827 | 0.25945197 | 5.1664E-26 | 5.9023E-23 |
| Niben101Scf09822g00004.1 | 456.969773 | 2.66155456 | 0.53364446 | 2.6188E-08 | 2.8409E-06 |
| Niben101Scf01398g00006.1 | 367.931661 | 2.64194236 | 0.58440237 | 1.7832E-07 | 1.5861E-05 |
| Niben101Scf10846g00019.1 | 33.5985748 | 2.62106897 | 0.67954323 | 1.4157E-06 | 9.9722E-05 |
| Niben101Scf01980g02002.1 | 17.8186759 | 2.61538686 | 0.5268791 | 2.9612E-08 | 3.1654E-06 |
| Niben101Scf02353g06039.1 | 31.5726099 | 2.59325749 | 0.4294485 | 1.047E-10 | 1.9332E-08 |
| Niben101Scf11706g00004.1 | 39.1686628 | 2.58158867 | 0.34807086 | 9.5347E-15 | 3.3197E-12 |
| Niben101Scf04988g02019.1 | 67.9465895 | 2.57864178 | 0.30732617 | 3.9797E-18 | 2.1086E-15 |
| Niben101Scf04973g02010.1 | 19.3224207 | 2.5758092 | 1.92232182 | 0.00010229 | 0.00393629 |
| Niben101Scf11706g00010.1 | 29.7028335 | 2.56762845 | 0.40626757 | 1.9009E-11 | 4.1121E-09 |
| Niben101Scf04570g06004.1 | 38.6932331 | 2.56557894 | 0.41023934 | 2.8836E-11 | 6.1566E-09 |
| Niben101Scf04881g01002.1 | 9.85499858 | 2.56004649 | 0.80320443 | 6.5867E-06 | 0.00038589 |
| Niben101Scf02115g04004.1 | 126.948087 | 2.55702499 | 1.17875667 | 3.9482E-05 | 0.00177319 |
| Niben101Scf02786g03020.1 | 12.6858953 | 2.54425662 | 0.94486932 | 1.6682E-05 | 0.00084956 |
| Niben101Scf03925g01010.1 | 65.8775001 | 2.54074158 | 0.33561368 | 3.1469E-15 | 1.162E-12 |
| Niben101Scf19766g00001.1 | 207.993951 | 2.54037696 | 0.35205333 | 4.4388E-14 | 1.387E-11 |
| Niben101Scf05166g01006.1 | 45.0123621 | 2.53715535 | 0.35836156 | 1.1874E-13 | 3.5581E-11 |
| Niben101Scf14163g00004.1 | 177.476364 | 2.51742515 | 0.20296616 | 2.3289E-36 | 5.3211E-33 |
| Niben101Scf00522g01017.1 | 61.7702901 | 2.51303984 | 1.24512937 | 5.1124E-05 | 0.00221181 |
| Niben101Scf02383g01001.1 | 151.442057 | 2.50758557 | 0.6210576 | 8.6883E-07 | 6.4637E-05 |
| Niben101Scf00530g11015.1 | 20.0396475 | 2.50468382 | 0.45822585 | 2.7315E-09 | 3.7825E-07 |
| Niben101Scf00225g00008.1 | 974.051057 | 2.5046564 | 0.14124347 | 2.2157E-71 | 2.025E-67 |
| Niben101Scf00712g02011.1 | 242.245022 | 2.49963692 | 1.88888029 | 0.00011397 | 0.00432195 |
| Niben101Scf00618g01019.1 | 18.5789519 | 2.49571829 | 0.53684896 | 1.1202E-07 | 1.0368E-05 |
| Niben101Scf04383g01014.1 | 71.7503203 | 2.48729876 | 0.57118347 | 3.2367E-07 | 2.7265E-05 |
| Niben101Scf05137g01028.1 | 113.040266 | 2.48713193 | 0.37784949 | 3.797E-12 | 8.9556E-10 |
| Niben101Scf06236g03019.1 | 66.9958196 | 2.48055371 | 0.31352483 | 2.3674E-16 | 1.0955E-13 |
| Niben101Scf04652g00027.1 | 30.638719 | 2.48007605 | 0.40791847 | 8.9549E-11 | 1.6788E-08 |
| Niben101Scf02437g01001.1 | 6.29284977 | 2.47870679 | 2.031826 | 0.00011529 | 0.00435858 |
| Niben101Scf03885g10002.1 | 10.1514491 | 2.46859783 | 1.15997681 | 4.4241E-05 | 0.00195806 |
| Niben101Scf05848g05012.1 | 349.199966 | 2.4684479 | 0.13159958 | 1.7107E-79 | 2.0846E-75 |
| Niben101Scf00456g00003.1 | 63.1090436 | 2.42136256 | 0.27112368 | 4.4571E-20 | 2.8586E-17 |
| Niben101Scf14778g00021.1 | 80.9723258 | 2.41820294 | 0.51631202 | 9.7484E-08 | 9.1851E-06 |
| Niben101Scf10650g02017.1 | 537.81072 | 2.39256544 | 0.38247318 | 3.2644E-11 | 6.8484E-09 |
| Niben101Scf01918g04014.1 | 52.8076386 | 2.39198173 | 0.34866979 | 6.4201E-13 | 1.7132E-10 |
| Niben101Scf06825g00036.1 | 19.5881001 | 2.37164988 | 0.70566428 | 4.6908E-06 | 0.00028629 |
| Niben101Scf06009g00025.1 | 541.990895 | 2.36891342 | 0.15751122 | 4.885E-52 | 2.5512E-48 |
| Niben101Scf07792g02034.1 | 21.6203295 | 2.3679244 | 0.50377106 | 9.04E-08 | 8.5618E-06 |
| Niben101Scf01176g01025.1 | 60.3401132 | 2.36692217 | 0.34133905 | 3.9423E-13 | 1.1002E-10 |
| Niben101Scf07123g01015.1 | 39.3799731 | 2.36581827 | 0.77539324 | 8.9476E-06 | 0.00050016 |
| Niben101Scf04362g07009.1 | 81.7848782 | 2.36023507 | 1.70297712 | 0.00013089 | 0.00484329 |
| Niben101Scf09870g00017.1 | 102.210032 | 2.35743457 | 2.09813975 | 0.00013004 | 0.0048166 |
| Niben101Scf01109g03005.1 | 34.6768794 | 2.3573168 | 0.44410672 | 6.21E-09 | 7.9658E-07 |
| Niben101Scf00307g01028.1 | 29.5966539 | 2.35444107 | 0.92379437 | 2.2958E-05 | 0.00112357 |
| Niben101Scf23113g00026.1 | 86.4055112 | 2.35299457 | 0.483372 | 4.5671E-08 | 4.5869E-06 |
| Niben101Scf07788g04025.1 | 54.9825241 | 2.33130376 | 0.36330518 | 1.2315E-11 | 2.7286E-09 |
| Niben101Scf11684g00005.1 | 68.4185878 | 2.32225844 | 0.31618784 | 2.1937E-14 | 7.2251E-12 |
| Niben101Scf00214g07004.1 | 92.8966236 | 2.31959534 | 0.69595663 | 4.9833E-06 | 0.00030313 |
| Niben101Scf01739g03006.1 | 27.2150576 | 2.31917574 | 0.47284146 | 3.8679E-08 | 3.9609E-06 |
| Niben101Scf00213g01002.1 | 216.005332 | 2.30532695 | 0.26813674 | 9.863E-19 | 5.3817E-16 |
| Niben101Scf03062g02017.1 | 233.921797 | 2.30390722 | 0.22729581 | 4.9234E-25 | 4.8646E-22 |
| Niben101Scf00773g08003.1 | 108.738708 | 2.29628263 | 0.30184246 | 3.1337E-15 | 1.162E-12 |
| Niben101Scf01241g02004.1 | 400.154154 | 2.29518378 | 0.26418582 | 4.5539E-19 | 2.6013E-16 |
| Niben101Scf03200g01014.1 | 18.8520572 | 2.29467809 | 0.65620167 | 3.4325E-06 | 0.00021938 |
| Niben101Scf00592g07009.1 | 23.3907093 | 2.29329231 | 0.44450435 | 1.2396E-08 | 1.4665E-06 |
| Niben101Scf11689g03003.1 | 129.506827 | 2.28802458 | 0.33657727 | 1.0481E-12 | 2.6795E-10 |
| Niben101Scf02918g00003.1 | 1335.27373 | 2.28730647 | 0.77770526 | 1.1221E-05 | 0.00061166 |
| Niben101Scf05916g00004.1 | 63.1180218 | 2.27922478 | 0.30461087 | 8.1571E-15 | 2.8674E-12 |
| Niben101Scf01696g06043.1 | 146.867162 | 2.27546404 | 0.30191201 | 5.4362E-15 | 1.9677E-12 |
| Niben101Scf00031g01005.1 | 524.190702 | 2.27362142 | 0.21593575 | 8.6042E-27 | 1.0485E-23 |
| Niben101Scf04046g00002.1 | 41.4532275 | 2.27049242 | 0.43338052 | 8.4497E-09 | 1.0366E-06 |
| Niben101Scf01719g08001.1 | 1946.38234 | 2.26970016 | 0.39903494 | 8.7415E-10 | 1.3427E-07 |
| Niben101Scf09230g02002.1 | 33.0054147 | 2.26767047 | 0.7288741 | 7.9762E-06 | 0.00045569 |
| Niben101Scf01322g03005.1 | 153.895162 | 2.26257489 | 0.22068068 | 1.5732E-25 | 1.7428E-22 |
| Niben101Scf05694g00009.1 | 93.3102685 | 2.25872767 | 0.40711465 | 1.8185E-09 | 2.6381E-07 |
| Niben101Scf14697g01004.1 | 741.50816 | 2.25629909 | 0.30841912 | 2.8291E-14 | 9.2174E-12 |
| Niben101Scf00975g01015.1 | 16.3438912 | 2.24561465 | 0.65062775 | 3.7837E-06 | 0.00023686 |
| Niben101Scf16105g00008.1 | 29.6337158 | 2.24248942 | 0.96203559 | 3.4986E-05 | 0.00160681 |
| Niben101Scf00317g07014.1 | 148.270487 | 2.23994214 | 0.31456771 | 1.1533E-13 | 3.4844E-11 |
| Niben101Scf08111g02011.1 | 119.761627 | 2.237893 | 0.426756 | 8.1128E-09 | 1.0046E-06 |
| Niben101Scf06509g02006.1 | 47.3406846 | 2.23788046 | 0.45747172 | 3.9162E-08 | 3.9991E-06 |
| Niben101Scf00428g09008.1 | 23.6948012 | 2.20881483 | 0.65316312 | 4.4088E-06 | 0.00027272 |
| Niben101Scf02348g05009.1 | 9.07215765 | 2.20833307 | 1.0203798 | 4.736E-05 | 0.00207104 |
| Niben101Scf05711g00021.1 | 127.670897 | 2.20259364 | 0.24815404 | 9.4435E-20 | 5.8515E-17 |
| Niben101Scf01212g03005.1 | 63.6530713 | 2.20056989 | 1.5383461 | 0.00014736 | 0.00526097 |
| Niben101Scf03925g02013.1 | 15.8805197 | 2.19809343 | 0.61524765 | 2.7983E-06 | 0.00018235 |
| Niben101Scf01495g00002.1 | 44.1231936 | 2.17266232 | 0.28249115 | 1.7616E-15 | 6.9247E-13 |
| Niben101Scf01297g04006.1 | 123.911397 | 2.16305327 | 0.39751866 | 2.9346E-09 | 4.0485E-07 |
| Niben101Scf03362g05002.1 | 329.311547 | 2.155146 | 0.31933733 | 1.4579E-12 | 3.6013E-10 |
| Niben101Scf01239g02002.1 | 962.92502 | 2.11688256 | 0.16244814 | 1.3969E-39 | 3.9282E-36 |
| Niben101Scf00539g05012.1 | 44.5176373 | 2.11223668 | 0.4125017 | 1.2771E-08 | 1.4965E-06 |
| Niben101Scf17561g00009.1 | 23.3175978 | 2.10882228 | 0.50516967 | 4.9015E-07 | 3.8869E-05 |
| Niben101Scf01432g06023.1 | 112.006388 | 2.1040834 | 0.26849241 | 5.6912E-16 | 2.4123E-13 |
| Niben101Scf06482g03003.1 | 59.0903285 | 2.08111376 | 0.28296184 | 2.1023E-14 | 7.051E-12 |
| Niben101Scf01249g03012.1 | 24.2656246 | 2.06958879 | 0.59817944 | 3.4881E-06 | 0.00022177 |
| Niben101Scf03925g01009.1 | 10.9312217 | 2.06841529 | 1.02251623 | 6.4298E-05 | 0.00269873 |
| Niben101Scf04099g05004.1 | 191.089691 | 2.06554714 | 0.33100531 | 3.2784E-11 | 6.8484E-09 |
| Niben101Scf01001g00003.1 | 2570.22119 | 2.06524261 | 1.46640111 | 0.00017162 | 0.00600954 |
| Niben101Scf11649g01009.1 | 13.2542252 | 2.05340274 | 1.47440698 | 0.000177 | 0.00612462 |
| Niben101Scf04015g00001.1 | 80.9122737 | 2.05085747 | 0.29326867 | 2.5999E-13 | 7.4163E-11 |
| Niben101Scf00370g03023.1 | 439.806944 | 2.04556015 | 0.12623286 | 8.8731E-60 | 6.4876E-56 |
| Niben101Scf02172g02008.1 | 120.600738 | 2.04149129 | 0.21327037 | 1.5739E-22 | 1.2509E-19 |
| Niben101Scf00160g06027.1 | 30.0207012 | 2.03522843 | 0.52306032 | 1.0809E-06 | 7.8713E-05 |
| Niben101Scf02513g05010.1 | 222.797722 | 2.03182351 | 0.18127158 | 6.2136E-30 | 9.0862E-27 |
| Niben101Scf01580g05004.1 | 79.3006896 | 2.0196126 | 0.36566979 | 1.5913E-09 | 2.3363E-07 |
| Niben101Scf04003g08001.1 | 552.85594 | 2.0091551 | 0.14919828 | 4.5117E-42 | 1.6494E-38 |
| Niben101Scf07165g00009.1 | 36.4706349 | 2.00349661 | 0.67979252 | 1.0922E-05 | 0.00060044 |
| Niben101Scf14679g00002.1 | 493.146066 | 2.00005232 | 0.15942506 | 7.4737E-37 | 1.8215E-33 |

**Table S3:** The up-regulated genes in the 671-CsnMN-treated *N. bethamiana*, seven days after TMV inoculation (Log2≥1.5).

| **Gene ID** | **baseMean** | **log2FoldChange** | **lfcSE** | **pvalue** | **padj** |
| --- | --- | --- | --- | --- | --- |
| Niben101Scf15817g01009.1 | 15.2869224 | 24.2154794 | 7.13475942 | 9.2699E-08 | 9.1812E-07 |
| Niben101Scf02452g05005.1 | 28.5401454 | 5.99106876 | 1.44575706 | 2.1933E-07 | 2.0199E-06 |
| Niben101Scf00607g03026.1 | 44.8636566 | 5.46348757 | 1.39797621 | 8.1979E-07 | 6.7313E-06 |
| Niben101Ctg07034g00003.1 | 199.882202 | 4.96297064 | 2.63300545 | 2.9221E-05 | 0.00017468 |
| Niben101Scf01160g00002.1 | 6.48643548 | 4.95711956 | 1.51441907 | 5.603E-06 | 3.9275E-05 |
| Niben101Scf00131g01009.1 | 16.2860272 | 3.17554425 | 0.73105915 | 2.4058E-07 | 2.1963E-06 |
| Niben101Ctg04867g00002.1 | 4.18917919 | 3.12488651 | 1.91509317 | 0.00032784 | 0.00151822 |
| Niben101Scf08294g01001.1 | 17.5186496 | 2.96741463 | 0.60674336 | 1.8648E-08 | 2.0791E-07 |
| Niben101Scf00135g03010.1 | 11.8553349 | 2.76239793 | 0.94936225 | 1.3415E-05 | 8.6646E-05 |
| Niben101Scf12404g00002.1 | 3071.556 | 2.51235834 | 0.21924862 | 8.5963E-32 | 2.551E-29 |
| Niben101Scf08926g05003.1 | 18.7559261 | 2.51078927 | 0.6343163 | 2.0394E-07 | 1.8913E-06 |
| Niben101Scf05290g00013.1 | 5.62701667 | 2.50389268 | 1.76110646 | 0.00076132 | 0.00320132 |
| Niben101Scf02077g12021.1 | 57.1237532 | 2.45951941 | 0.41844269 | 1.6236E-11 | 3.1267E-10 |
| Niben101Scf04563g02007.1 | 220.549236 | 2.43212189 | 0.34409435 | 8.1071E-15 | 2.8682E-13 |
| Niben101Scf01578g00012.1 | 59.8919039 | 2.33997696 | 0.41322405 | 2.6005E-11 | 4.8144E-10 |
| Niben101Scf06799g04001.1 | 57.5119317 | 2.33663507 | 0.40011258 | 9.214E-12 | 1.8751E-10 |
| Niben101Scf05044g00015.1 | 33.5072917 | 2.31627303 | 0.43028895 | 1.0732E-10 | 1.7924E-09 |
| Niben101Scf02829g01014.1 | 13.9402071 | 2.31585649 | 0.95489453 | 4.5749E-05 | 0.00026143 |
| Niben101Scf06584g00004.1 | 486.471918 | 2.27864802 | 0.42763055 | 1.1795E-10 | 1.9608E-09 |
| Niben101Scf02160g00026.1 | 4.13481941 | 2.26344246 | 1.76196266 | 0.00118985 | 0.00475105 |
| Niben101Scf00143g03005.1 | 7.80380108 | 2.20880426 | 0.98439273 | 8.2104E-05 | 0.00044193 |
| Niben101Scf01817g01001.1 | 22.6603356 | 2.20537437 | 0.70341539 | 2.9776E-06 | 2.2025E-05 |
| Niben101Scf03289g02002.1 | 18.6017334 | 2.18238446 | 0.62558049 | 6.6442E-07 | 5.5718E-06 |
| Niben101Scf01784g05005.1 | 39.365142 | 2.15318758 | 0.4105791 | 1.0169E-10 | 1.7062E-09 |
| Niben101Scf00978g06005.1 | 684.092526 | 2.08869536 | 0.23255802 | 7.0765E-23 | 8.8492E-21 |
| Niben101Scf07812g00013.1 | 603.116731 | 2.08034684 | 0.24503785 | 5.2785E-21 | 5.0947E-19 |
| Niben101Scf02854g11002.1 | 54.5426965 | 2.05481237 | 0.40828851 | 2.3401E-10 | 3.6942E-09 |
| Niben101Scf00339g16007.1 | 165.837649 | 2.05465889 | 0.23602892 | 6.4291E-22 | 6.9852E-20 |
| Niben101Scf02401g02009.1 | 6.25569874 | 2.04411221 | 1.07144739 | 0.00025605 | 0.00121868 |
| Niben101Scf05483g08004.1 | 771.043774 | 2.02547467 | 0.48286441 | 1.9083E-08 | 2.1256E-07 |
| Niben101Scf08604g00003.1 | 49.8828788 | 2.00105926 | 0.3447414 | 1.9421E-12 | 4.4848E-11 |
| Niben101Scf10409g02009.1 | 2020.12878 | 1.98841571 | 0.28202831 | 3.3808E-16 | 1.5226E-14 |
| Niben101Scf01550g00008.1 | 11.4650755 | 1.97554015 | 0.78213648 | 2.909E-05 | 0.00017401 |
| Niben101Scf02572g02024.1 | 2874.33743 | 1.97372397 | 0.24610698 | 1.4713E-19 | 1.1405E-17 |
| Niben101Scf03422g01002.1 | 1186.58747 | 1.96958357 | 0.24542676 | 1.3823E-19 | 1.08E-17 |
| Niben101Scf05692g04001.1 | 63.7827673 | 1.96216077 | 0.4968344 | 6.0269E-08 | 6.1312E-07 |
| Niben101Scf04187g00031.1 | 44.8472713 | 1.95730513 | 0.41819744 | 1.3716E-09 | 1.8698E-08 |
| Niben101Scf05886g01005.1 | 7.66112125 | 1.95535907 | 1.01500941 | 0.00024945 | 0.00119151 |
| Niben101Scf05554g05007.1 | 3.28113801 | 1.95103757 | 1.71349957 | 0.00207848 | 0.007747 |
| Niben101Scf00482g02014.1 | 21.8794386 | 1.94338299 | 0.53580746 | 2.779E-07 | 2.5054E-06 |
| Niben101Scf03127g00002.1 | 3.04252195 | 1.92924808 | 1.70622011 | 0.00215759 | 0.00800244 |
| Niben101Scf07231g04011.1 | 26.7681489 | 1.92628665 | 0.45596915 | 1.5007E-08 | 1.7017E-07 |
| Niben101Scf10999g00004.1 | 179.802595 | 1.91343835 | 0.23855483 | 1.2116E-19 | 9.5411E-18 |
| Niben101Scf00823g00001.1 | 1.9673775 | 1.90876783 | 1.82891875 | 0.00240506 | 0.00880012 |
| Niben101Scf02011g03011.1 | 133.359057 | 1.89831277 | 0.29363684 | 1.934E-14 | 6.3508E-13 |
| Niben101Scf02122g02016.1 | 7.29799673 | 1.89407975 | 0.94464456 | 0.00019779 | 0.00096875 |
| Niben101Scf08918g00010.1 | 293.683434 | 1.89293343 | 0.22973585 | 1.831E-20 | 1.6362E-18 |
| Niben101Scf04328g00049.1 | 5.51889163 | 1.88669234 | 1.38896026 | 0.00142341 | 0.0055586 |
| Niben101Ctg14049g00002.1 | 3180.21687 | 1.8763352 | 0.38203451 | 3.6911E-10 | 5.6217E-09 |
| Niben101Scf09589g00006.1 | 27.1790993 | 1.86478606 | 0.54432122 | 7.1524E-07 | 5.9502E-06 |
| Niben101Scf02983g02002.1 | 10.9796244 | 1.84166552 | 0.97710662 | 0.0003072 | 0.00143363 |
| Niben101Scf01383g00009.1 | 2.56151781 | 1.82732045 | 1.84233817 | 0.0027753 | 0.00995873 |
| Niben101Scf02195g04002.1 | 2.21843841 | 1.82623415 | 1.77701432 | 0.00274984 | 0.00988343 |
| Niben101Scf39395g00004.1 | 49.4374972 | 1.81992332 | 0.43384866 | 1.8735E-08 | 2.088E-07 |
| Niben101Scf06286g03008.1 | 133.800523 | 1.81877652 | 0.33647321 | 2.1565E-11 | 4.0642E-10 |
| Niben101Scf04113g17024.1 | 2.55618934 | 1.80567942 | 1.75654402 | 0.00283694 | 0.01014696 |
| Niben101Scf00879g03005.1 | 47.7346995 | 1.79010765 | 0.36920127 | 6.0044E-10 | 8.8076E-09 |
| Niben101Scf06203g02004.1 | 9.98790583 | 1.78972186 | 0.9336336 | 0.00028853 | 0.00135574 |
| Niben101Scf06529g06013.1 | 168.139998 | 1.7815592 | 0.2276697 | 7.4631E-19 | 5.1794E-17 |
| Niben101Scf01998g10004.1 | 392.101444 | 1.77706257 | 0.26952133 | 9.6161E-15 | 3.3397E-13 |
| Niben101Scf00875g07011.1 | 10.245184 | 1.7751965 | 0.81332688 | 0.00011739 | 0.00060799 |
| Niben101Scf01716g00015.1 | 458.50937 | 1.76289398 | 0.2378441 | 2.2625E-17 | 1.2444E-15 |
| Niben101Scf01188g00028.1 | 5.00980872 | 1.75077469 | 1.60664454 | 0.00286833 | 0.01024909 |
| Niben101Scf03004g00014.1 | 4.49499848 | 1.74321733 | 1.28365294 | 0.00163006 | 0.00625794 |
| Niben101Scf06305g01018.1 | 18.1220117 | 1.73419322 | 0.64318413 | 1.8325E-05 | 0.00011476 |
| Niben101Scf08041g02005.1 | 52.759048 | 1.73251169 | 0.37476868 | 2.4907E-09 | 3.2452E-08 |
| Niben101Scf06216g01014.1 | 94.5744558 | 1.73165606 | 0.3382123 | 1.5565E-10 | 2.5336E-09 |
| Niben101Scf09112g03002.1 | 103.681512 | 1.73129855 | 0.29577162 | 1.7728E-12 | 4.142E-11 |
| Niben101Scf01374g15012.1 | 285.130884 | 1.7150352 | 0.26882482 | 5.8929E-14 | 1.7793E-12 |
| Niben101Scf03114g07010.1 | 60.9934485 | 1.71086246 | 0.32927489 | 1.1026E-10 | 1.8361E-09 |
| Niben101Scf01063g09001.1 | 49.2386157 | 1.70675511 | 0.41620177 | 4.0372E-08 | 4.2477E-07 |
| Niben101Scf04407g02027.1 | 127.438326 | 1.70007591 | 0.26436203 | 4.5664E-14 | 1.4098E-12 |
| Niben101Scf00369g05011.1 | 3784.21557 | 1.68423424 | 0.27736948 | 5.676E-13 | 1.445E-11 |
| Niben101Scf05867g02007.1 | 38.5057987 | 1.67091902 | 0.46305398 | 4.6763E-07 | 4.0303E-06 |
| Niben101Scf02751g05008.1 | 1.65231517 | 1.66956146 | 1.73538295 | 0.00364556 | 0.01258957 |
| Niben101Scf03634g06014.1 | 9.99720755 | 1.66857867 | 0.8790298 | 0.00035301 | 0.0016212 |
| Niben101Scf02821g14008.1 | 2.03344982 | 1.66842675 | 1.80353907 | 0.00364526 | 0.01258957 |
| Niben101Scf03283g04012.1 | 2729.97901 | 1.66459707 | 0.23153231 | 2.5233E-16 | 1.1654E-14 |
| Niben101Scf00560g09010.1 | 2.22428152 | 1.66324985 | 1.68811935 | 0.00364587 | 0.01258957 |
| Niben101Scf10262g00005.1 | 16.8817342 | 1.65985224 | 0.86256151 | 0.0003283 | 0.00151982 |
| Niben101Scf04049g00024.1 | 43.4713583 | 1.65570995 | 0.39178485 | 2.6973E-08 | 2.9298E-07 |
| Niben101Scf05584g04007.1 | 180.646851 | 1.65337713 | 0.2602182 | 1.0954E-13 | 3.1657E-12 |
| Niben101Scf05252g02010.1 | 330.979489 | 1.65210988 | 0.30552405 | 4.4836E-11 | 7.9739E-10 |
| Niben101Scf02922g00017.1 | 20.548245 | 1.65159722 | 0.48018251 | 1.0572E-06 | 8.5024E-06 |
| Niben101Scf00819g11005.1 | 82.7541852 | 1.65051812 | 0.53079552 | 4.2656E-06 | 3.0602E-05 |
| Niben101Scf01160g01003.1 | 4444.65601 | 1.64823718 | 0.25327895 | 3.9978E-14 | 1.244E-12 |
| Niben101Scf16582g00010.1 | 16.7620041 | 1.63885241 | 0.67227707 | 5.835E-05 | 0.00032546 |
| Niben101Scf03961g00004.1 | 15592.8336 | 1.62770879 | 0.23918002 | 5.946E-15 | 2.1613E-13 |
| Niben101Scf10889g01017.1 | 125.492036 | 1.61481893 | 0.27920822 | 6.0776E-12 | 1.2767E-10 |
| Niben101Scf02824g03028.1 | 24.6020232 | 1.61245368 | 0.48752172 | 2.1493E-06 | 1.6314E-05 |
| Niben101Scf00607g03013.1 | 8049.51294 | 1.60515868 | 0.24076723 | 1.9687E-14 | 6.4594E-13 |
| Niben101Scf08263g01001.1 | 2.20312136 | 1.60500539 | 1.6356755 | 0.00401905 | 0.01368102 |
| Niben101Scf04714g00004.1 | 876.570023 | 1.60482719 | 0.26494371 | 1.1773E-12 | 2.8444E-11 |
| Niben101Scf03057g00001.1 | 4.70242844 | 1.60287243 | 1.26409217 | 0.00239606 | 0.00877122 |
| Niben101Scf02211g02012.1 | 60.8628086 | 1.58338542 | 0.2661203 | 2.8378E-12 | 6.3864E-11 |
| Niben101Scf07034g08030.1 | 28.7590363 | 1.58259117 | 0.42753643 | 4.5332E-07 | 3.9163E-06 |
| Niben101Scf04287g08001.1 | 250.03684 | 1.57121629 | 0.31953565 | 1.297E-09 | 1.7743E-08 |
| Niben101Scf00328g00001.1 | 3.94401967 | 1.56804323 | 1.31709331 | 0.00299846 | 0.01066138 |
| Niben101Scf02128g00016.1 | 43.4326167 | 1.56516582 | 0.38955774 | 1.2005E-07 | 1.1607E-06 |
| Niben101Scf01742g01001.1 | 161.987149 | 1.5645134 | 0.32503588 | 2.3728E-09 | 3.1028E-08 |
| Niben101Scf00123g04002.1 | 474.215632 | 1.56396667 | 0.25125065 | 5.8904E-13 | 1.4967E-11 |
| Niben101Scf13600g01002.1 | 106.897892 | 1.56214176 | 0.40779728 | 2.8577E-07 | 2.5693E-06 |
| Niben101Scf03839g12003.1 | 551.255425 | 1.56175506 | 0.25171149 | 6.8694E-13 | 1.7322E-11 |
| Niben101Scf00537g00005.1 | 338.613573 | 1.55798497 | 0.280909 | 4.2044E-11 | 7.5143E-10 |
| Niben101Scf02853g06014.1 | 574.040261 | 1.55467842 | 0.27867091 | 3.5736E-11 | 6.4628E-10 |
| Niben101Scf02451g01004.1 | 271.914105 | 1.55394834 | 0.27403053 | 2.082E-11 | 3.9389E-10 |
| Niben101Scf07036g02002.1 | 12.9646639 | 1.55326013 | 0.63668895 | 7.2077E-05 | 0.00039333 |
| Niben101Ctg06858g00001.1 | 19.0023708 | 1.55093985 | 0.70960926 | 0.00017115 | 0.00085139 |
| Niben101Scf01546g00008.1 | 116.294249 | 1.54903503 | 0.28688992 | 1.0722E-10 | 1.7915E-09 |
| Niben101Scf05250g02013.1 | 1636.77808 | 1.54656999 | 0.24331666 | 2.9758E-13 | 7.9633E-12 |
| Niben101Scf01980g04003.1 | 79.835809 | 1.54406881 | 0.28776234 | 1.3598E-10 | 2.2373E-09 |
| Niben101Scf04220g01008.1 | 16.8261303 | 1.5432719 | 0.62964887 | 7.1365E-05 | 0.00038998 |
| Niben101Scf03370g04008.1 | 22.8596649 | 1.54302279 | 0.70168489 | 0.00016702 | 0.00083282 |
| Niben101Scf01687g02010.1 | 45.9000786 | 1.53748917 | 0.35443203 | 3.1886E-08 | 3.4205E-07 |
| Niben101Scf14803g00004.1 | 495.401283 | 1.53630285 | 0.18941719 | 7.2402E-19 | 5.0511E-17 |
| Niben101Scf06578g02001.1 | 469.351705 | 1.53424242 | 0.18768049 | 4.3673E-19 | 3.1236E-17 |
| Niben101Scf03481g00014.1 | 679.00759 | 1.53110577 | 0.17068658 | 4.4334E-22 | 4.9108E-20 |
| Niben101Scf09089g01026.1 | 1868.23479 | 1.52847415 | 0.15582845 | 1.58E-25 | 2.6766E-23 |
| Niben101Scf00247g05011.1 | 180.507206 | 1.51859786 | 0.20795776 | 5.2337E-16 | 2.25E-14 |
| Niben101Scf02907g07021.1 | 23.7985089 | 1.51734774 | 0.54736564 | 2.5152E-05 | 0.00015224 |
| Niben101Scf06405g00006.1 | 468.321989 | 1.51693933 | 0.20652451 | 3.8793E-16 | 1.7198E-14 |
| Niben101Scf07103g03015.1 | 136.147611 | 1.51617295 | 0.27121118 | 4.831E-11 | 8.5383E-10 |
| Niben101Scf02063g02014.1 | 690.98707 | 1.51068819 | 0.14428106 | 2.3113E-28 | 5.0225E-26 |
| Niben101Scf02264g06029.1 | 54.4455314 | 1.50675864 | 0.28871904 | 4.3787E-10 | 6.5782E-09 |
| Niben101Scf02749g04008.1 | 14641.047 | 1.50487678 | 0.24852832 | 3.2339E-12 | 7.1923E-11 |
| Niben101Scf06684g04001.1 | 72.4214609 | 1.50442419 | 0.26656748 | 3.9712E-11 | 7.1167E-10 |
| Niben101Scf05749g00010.1 | 1483.22647 | 1.50015063 | 0.35836721 | 8.5694E-08 | 8.5236E-07 |
